# JointPRS: A Data-Adaptive Framework for Multi-Population Genetic Risk Prediction Incorporating Genetic Correlation

**DOI:** 10.1101/2023.10.29.564615

**Authors:** Leqi Xu, Geyu Zhou, Wei Jiang, Haoyu Zhang, Yikai Dong, Leying Guan, Hongyu Zhao

## Abstract

Genetic prediction accuracy for non-European populations is hindered by the limited sample size of Genome-wide association studies (GWAS) data in these populations. Additionally, it is challenging to tune model parameters with a small tuning dataset for methods that require tuning data, which is often the case for non-European samples. To address these challenges, we propose JointPRS, a novel, data-adaptive framework that simultaneously models multiple populations using GWAS summary statistics. JointPRS incorporates genetic correlation structures into the prediction framework, enabling accurate performance even without individual-level tuning data. Additionally, it uniquely employs a data-adaptive approach, providing a robust solution when only a small tuning dataset is available. Through extensive simulations and real data applications to 22 quantitative traits and four binary traits in five continental populations (European (EUR); East Asian (EAS); African (AFR); South Asian (SAS); and Admixed American (AMR)) evaluated using the UK Biobank (UKBB) and All of Us (AoU), we demonstrate that JointPRS outperforms six other state-of-art methods across three different data scenarios (no tuning data, tuning and testing data from the same cohort, and tuning and testing data from different cohorts) for most traits in non-European populations, while maintaining model simplicity and computational efficiency.

## 1 Introduction

Polygenic Risk Scores (PRS), the weighted sum of risk alleles across a collection of genetic variants, have seen active development for predicting complex traits in recent years. PRS have demonstrated their ability to identify individuals with high disease risk, which can be applied for disease monitoring, prevention, and treatment [1–4]. However, most PRS have been developed for European populations, due to the dominance of large European cohorts in Genome-Wide Associations Studies (GWAS) [5], with non-European populations’ PRS exhibiting reduced accuracy in comparison [6]. Several factors contribute to the reduced prediction accuracy of PRS in non-European populations. First, both GWAS summary statistics for model training and individual-level datasets for parameter tuning are limited in these populations, hindering the development of population-specific PRS that rely solely on non-European datasets. Additionally, distinctive genetic structures, such as different linkage disequilibrium (LD) patterns and varying numbers of single nucleotide polymorphisms (SNPs) between European and non-European populations, constrain the transferability of European PRS prediction models to non-European populations. This gap in genetic risk prediction performance can exacerbate health disparities [6, 7], underscoring the urgent need to improve PRS prediction accuracy for non-European populations.

To address this need, there has been an increase in GWAS focused on non-European populations [8–16], complemented by the development of various models tailored for multi-population PRS predictions [17–29]. These models can be categorized into two types: “auto methods” and “tune methods”. Auto methods, which utilize only GWAS summary statistics, can provide robust performance with or without individual-level tuning data [20]. However, neglecting individual-level tuning data when available may miss useful information that could further improve PRS prediction accuracy. But tune methods require an individual-level tuning dataset, limiting their applications in various data scenarios.

When individual-level tuning data are available, tune methods can enhance the prediction accuracy of PRS. However, their performance is significantly influenced by the quality and sample size of the tuning data, a challenge particularly pronounced in non-European populations. The quality of tuning datasets can be compromised by confounding factors and measurement errors of the phenotype being predicted and the chosen tuning dataset. Additionally, various factors such as socio-economic status, age, and sex in the individual-level dataset can lead to varying PRS prediction accuracy across individuals [30]. Including individuals who are not effectively predicted by PRS due to these factors can adversely affect tuning weights and overall model performance. Small sample sizes further exacerbate the issue, potentially leading to unstable performance when PRS are applied to new datasets. Consequently, there remains a need for methods that not only provide accurate and robust predictions without tuning data but also can effectively utilize individual-level tuning data to achieve both efficiency and robustness.

In addition to these challenges, cross-population genetic correlations has proven to be valuable in existing multi-population PRS methods. However, there are limitations in these methods. For instance, SDPRX [20] relies on genetic-correlation estimates from external software like Popcorn [31], which can lead to model mismatch and less accurate PRS predictions. XPASS [17] employs the Method of Moments for estimating cross-population genetic correlations, potentially resulting in unconstrained estimates outside the valid range of [−1, 1]. Furthermore, methods like MUSSEL and PROSPER [22, 23] treat genetic correlations as tuning parameters within their models, restricting their applicability when tuning data are not available. Thus, challenges remain in effectively incorporating cross-population genetic correlations within multi-population PRS models.

Moreover, most published studies evaluate method performance by splitting a single individual-level data cohort into tuning and testing datasets [17, 19–24, 29]. However, in real-world PRS applications, such as predicting phenotypes for individuals in a medical center, tuning data from the same cohort are often unavailable. Instead, we may have no tuning data at all or only tuning data from other cohorts. Therefore, there is a need to benchmark PRS methods under diverse data scenarios, including no tuning data, tuning and testing data from the same cohort, and tuning and testing data from different cohorts. This comprehensive evaluation, which remains largely unexplored, is essential to ensure the robustness and applicability of multi-population PRS methods in practical settings.

In this study, we introduce JointPRS, a Bayesian framework that employs sparse distribution assumptions for genetic variant effect sizes, jointly models multiple populations, and incorporates chromosome-wise cross-population genetic correlations. JointPRS uses a continuous shrinkage (CS) prior to flexibly account for varying sparsity levels in the genetic variant effect sizes, which is also utilized in PRS-CSx [19]. However, JointPRS offers two distinct advantages: First, JointPRS incorporates cross-population genetic correlation structures, ensuring accurate performance even when only GWAS summary statistics are available for training. In contrast, PRS-CSx does not consider genetic correlations, limiting its prediction accuracy, particularly when tuning data are not available.

Second, when non-European individual-level tuning data are available, JointPRS adopts a data-adaptive approach that combines meta-analysis with tuning strategies.

This approach addresses the challenge of small non-European tuning datasets, resulting in more robust performance. Conversely, the tuning method in PRS-CSx relies on the quality of individual-level tuning data, leading to unstable PRS performances. By leveraging these innovations, JointPRS enhances the robustness and accuracy of PRS across diverse populations.

The efficacy of JointPRS is evaluated through extensive simulations and applications to real data, including 22 quantitative traits and four binary traits across five populations (European (EUR); East Asian (EAS); African (AFR); South Asian (SAS); and Admixed American (AMR)) using the UK Biobank (UKBB) [32] and All of Us (AoU) cohorts [33]. Evaluations are conducted under three data scenarios: [1] no tuning data, [2] tuning and testing data from the same cohort, and [3] tuning and testing data from different cohorts. JointPRS consistently outperforms other state-of-the-art methods [17, 19, 20, 22–24] across most traits in non-European populations and various data scenarios while maintaining model simplicity and achieving computational efficiency. Our results illustrate the contributions of appropriate sparse distribution, joint modeling of multiple populations, and genetic correlation structures to prediction accuracy. Additionally, our findings highlight the importance of the data-adaptive approach for robust predictions when only a small tuning dataset is available. Moreover, our simulation studies underscore the impact of tuning sample sizes on PRS prediction accuracy. In conclusion, JointPRS offers a robust and accurate apporach for genetic risk prediction in diverse populations, with or without tuning datasets. This work also enhances our understanding of the contributions of different types of information to impacting PRS prediction.

## 2 Results

### 2.1 Overview of JointPRS

JointPRS is a Bayesian high-dimensional prediction framework designed for multiple populations with an additional data-adaptive approach to determine how to use tuning datasets when they are available. This framework operates in two key steps (Figure 1): [1] model development for multiple populations, and [2] data-adaptive approach to select the optimal PRS between meta and tune versions when a tuning dataset is available. An independent testing dataset is used to evaluate prediction performance.

**Fig. 1.**
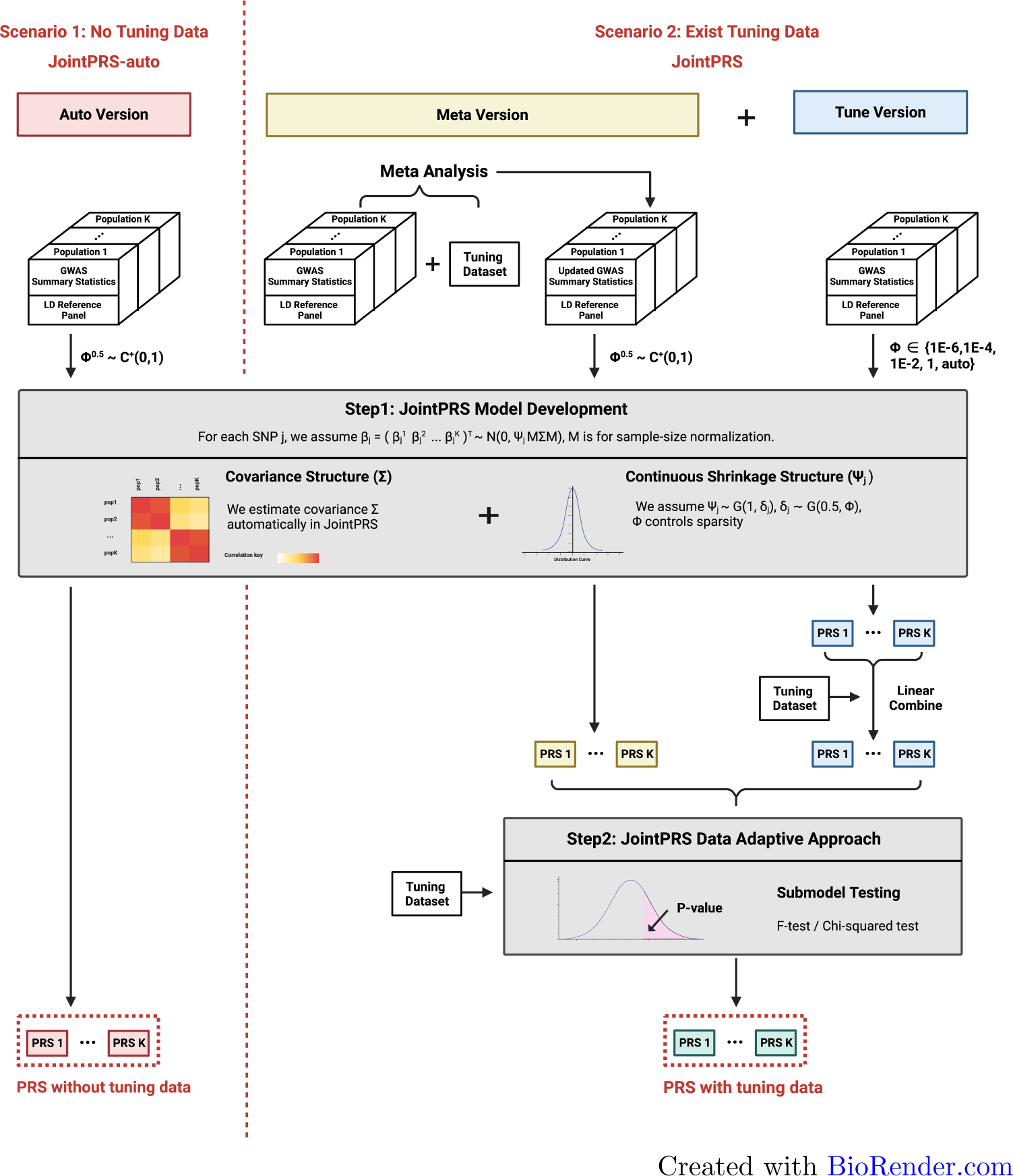
JointPRS Workflow. The pipeline of using JointPRS for PRS estimation under two scenarios: without and with a tuning dataset.

**Step1: JointPRS Model Development**. In Step 1, various versions of the Joint-PRS model are trained using appropriate training inputs. First, the JointPRS model is introduced, followed by an explanation of each version and their application scenarios.

#### JointPRS Model

The JointPRS model has the following form:

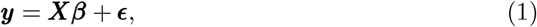

where

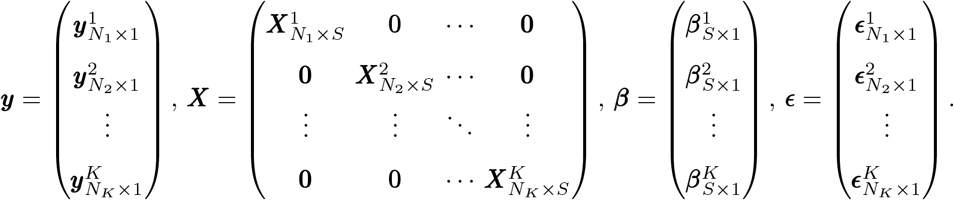

Here, *K* is the number of populations, *N*_*k*_ denotes the sample size for population *k*, and *S* represents the number of SNPs that are available in at least one population. The vectors ***y***^*k*^, ***X***^*k*^, ***β***^*k*^ and ***ϵ***^*k*^ correspond to the standardized phenotype vector, the column-standardized genotype matrix, the standardized effect size vector for SNPs, and residuals in population *k* respectively. There may be missing elements in the genotype matrix and effect size vector, which will be discussed in detail later.

When SNP *j* is available for all *K* populations, the effect size *β*_*j*_ for SNP *j* across *K* populations is modeled with a correlated Gaussian prior:

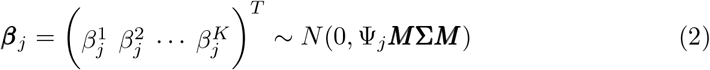

with

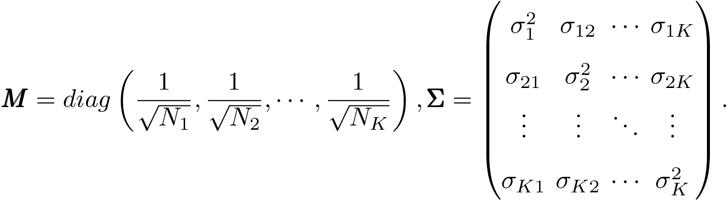

Here, Ψ_*j*_ represents the sample-size normalized effect size for SNP *j* shared across *K* populations. This shared effect, Ψ_*j*_, is modeled using a continuous shrinkage prior, as implemented in the PRS-CSx framework [19]. Specifically, Ψ_*j*_ follows a gamma distribution Ψ_*j*_ ∼ *G*(1, *δ*_*j*_), with 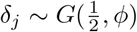. The global shrinkage parameter *ϕ* varies depending on the JointPRS version used, which will be detailed later. Additionally, we introduce the cross-population covariance matrix **Σ** in the model to capture shared patterns across populations. All parameters in the model are estimated automatically by the JointPRS algorithm using only GWAS summary statistics **(Methods)**.

For SNPs available in only a subset of populations, the model truncates to include only those populations with the SNP, using an index matrix **(Methods)**.

#### JointPRS Versions

The JointPRS modeling framework has three versions—auto, meta, and tune—each differing in terms of input data, model parameters, and modeling strategies (Table 1). Detailed illustrations and implementations of these different JointPRS versions are provided in the Methods section **(Methods)**.

**Table 1.** JointPRS model differences in different versions.

| JointPRS version | Input data | Model parameters | Modeling strategies |
| --- | --- | --- | --- |
| Auto version | original GWAS<br>summary statistics | $\phi^{\frac{1}{2}} \sim C^+(0, 1)$ | no parameter tuning |
| Meta version | meta-analyzed GWAS<br>summary statistics | $\phi^{\frac{1}{2}} \sim C^+(0, 1)$ | no parameter tuning |
| Tune version | original GWAS<br>summary statistics | $\phi^{\frac{1}{2}} \sim C^+(0, 1)$ or<br>$\phi \in \{10^{-6}, 10^{-4}, 10^{-2}, 1\}$ | linear combination<br>across populations and<br>optimal parameter selection |

When tuning data are unavailable, JointPRS employs the auto version to construct the PRS, referred to as JointPRS-auto. When tuning data are available, JointPRS uses a data-adaptive approach to select between the meta version and the tune version, as detailed in Step2. This data-adaptive PRS approach is referred to as JointPRS. JointPRS-auto and JointPRS together cover data scenarios without and with tuning data, with the latter adaptively deciding how the available tuning data are used.

**Step2: JointPRS Data-Adaptive Approach**. Step2 is performed only when the tuning data are available, aiming to select the optimal PRS between the meta version and the tune version. Directly evaluating the meta version using the tuning dataset can lead to overfitting, as the meta version already incorporates the tuning dataset into the original GWAS summary statistics during training. To mitigate this problem, we approximate the meta version’s performance by using the tuning data to evaluate the performance of the auto version, which employs the global shrinkage prior 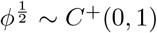 with the original GWAS summary statistics. Additionally, we evaluate the tune version by using 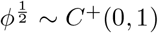, followed by a linear combination across populations, skipping the parameter selection step. The key difference in the evaluation of the two versions is whether a linear combination across populations is performed. This approach frames the choice between the meta version and the tune version as a model selection problem, using the tuning dataset to determine whether the linear combination across populations is beneficial. This selection is achieved through the F-test for continuous traits and chi-squared test for binary traits **(Methods)**.

### 2.2 Simulations

#### 2.2.1 Simulation Settings

##### Benchmarking Methods

We compared the performance of JointPRS with six other state-of-the-art multi-population PRS methods: XPASS [17], SDPRX [20], PRS-CSx [19], MUSSEL [22], PROSPER [23], and BridgePRS [24] **(Methods)**. These methods can be grouped into two categories: auto methods and tune methods (Figure S1). Auto methods, including JointPRS-auto, XPASS, SDPRX, and PRS-CSx-auto, can be implemented using only GWAS summary statistics. When a tuning dataset is available, XPASS and SDPRX incorporate the tuning data with GWAS summary statistics to increase sample size. Tune methods, including JointPRS, PRS-CSx, MUSSEL, PROSPER, and BridgePRS, require a tuning dataset with corresponding tuning strategies for application. We benchmarked seven methods across various simulation settings, including different sample sizes for training, tuning, and testing datasets and various genetic architectures of a continuous trait.

##### Genetic Dataset Sample Sizes Settings

For the European population, we used the UKBB genotype data, which include 311,600 unrelated samples for training in the simulation. Additionally, we used 10,000 simulated individuals for tuning and 10,000 for testing from a large simulation dataset [21]. Briefly, this simulated dataset comprises a total of 600,000 independent samples with reference LD based on the 1000 Genomes Project data [34]. There are 120,000 samples representing each of the five continental populations: EUR, EAS, AFR, SAS, and AMR.

For the non-European populations, we obtained all genotype datasets from the large simulated dataset [21]. We considered two training sample sizes: 80,000 and 15,000 samples. In each scenario, based on the availability and sample size of the tuning dataset, we examined the following situations:

1. **No tuning dataset available:** We compared the performance of methods that only require GWAS summary statistics, including JointPRS-auto, XPASS, SDPRX, and PRS-CSx-auto.
2. **Tuning dataset available:** We compared the performance of all methods: JointPRS, XPASS, SDPRX, PRS-CSx, MUSSEL, PROSPER, and BridgePRS, with varying tuning dataset sample sizes (500, 2,000, 5,000, and 10,000).

In all scenarios, we evaluated the performance of all methods using 10,000 testing individuals. For all methods, we jointly modeled the maximum number of populations each method could utilize: five populations for JointPRS(-auto), PRS-CSx(-auto), MUSSEL, and PROSPER, and two populations for XPASS, SDPRX, and BridgePRS.

##### Genetic Architecture Settings

We simulated the true effect sizes using a spike and slab model for five populations: EUR, EAS, AFR, SAS, and AMR:

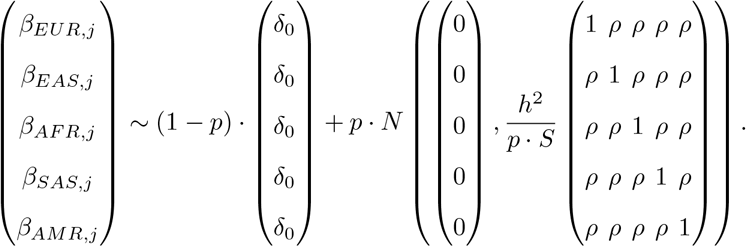

We considered four different sparsity levels for the proportion of causal SNPs: *p* = 0.1, 0.01, 0.001, and 5 × 10^*−*4^. The total heritability of common SNPs was set to *h*^2^ = 0.4, and the cross-population genetic correlation was set to *ρ* = 0.8 in our primary analysis. We also considered four genetic correlation settings *ρ* = 0, 0.2, 0.4, 0.6, and 0.8 with fixed casual SNPs proportion *p* = 0.1 and heritability *h*^2^ = 0.4 as our secondary analysis. The total number of SNPs was *S* = 1, 203, 063, based on the HapMap3 SNP list [35]. We generated five replicates of the effect sizes for each setting.

Using these parameters and effect sizes, we utilized GCTA-sim [36] to generate phenotypes for analysis in the training, tuning, and testing datasets. PLINK2 [37] was used to derive GWAS summary statistics in the training dataset. For the GWAS summary statistics provided for each method, we extracted SNPs common to all methods’ LD reference panels across all populations, resulting in 717, 985 SNPs. This intersection ensures that each method used the same number of SNPs in the GWAS data for fair comparisons.

#### 2.2.2 Simulation Results

Simulation results (Figures 2 - 4 and Figures S2 - S9; Table 2 and Tables S1 - S5) demonstrate that JointPRS consistently performs well across various tuning dataset scenarios, non-European population training sample sizes, and genetic architecture settings.

**Table 2.** Simulation Results for Counting Each Method having Top Two Performance Based on the Mean Value Across Five Replications When a Tuning Dataset is Available in the EAS, AFR, SAS, and AMR Populations.

| Pop | Method | $p = 0.1, 0.01$ | | | Method | $p = 0.001, 5 \times 10^{-4}$ | | |
| --- | --- | --- | --- | --- | --- | --- | --- | --- |
|  |  | Top2 | best | second best |  | Top2 | best | second best |
| EAS | JointPRS | 15 | 11 | 4 | JointPRS | 11 | 5 | 6 |
|  | XPASS | 0 | 0 | 0 | XPASS | 0 | 0 | 0 |
|  | SDPRX | 3 | 1 | 2 | SDPRX | 10 | 8 | 2 |
|  | PRS-CSx | 14 | 4 | 10 | PRS-CSx | 11 | 3 | 8 |
|  | MUSSEL | 0 | 0 | 0 | MUSSEL | 0 | 0 | 0 |
|  | PROSPER | 0 | 0 | 0 | PROSPER | 0 | 0 | 0 |
|  | BridgePRS | 0 | 0 | 0 | BridgePRS | 0 | 0 | 0 |
| AFR | JointPRS | 13 | 5 | 8 | JointPRS | 2 | 0 | 2 |
|  | XPASS | 0 | 0 | 0 | XPASS | 0 | 0 | 0 |
|  | SDPRX | 2 | 1 | 1 | SDPRX | 15 | 7 | 8 |
|  | PRS-CSx | 8 | 6 | 2 | PRS-CSx | 4 | 1 | 3 |
|  | MUSSEL | 9 | 4 | 5 | MUSSEL | 11 | 8 | 3 |
|  | PROSPER | 0 | 0 | 0 | PROSPER | 0 | 0 | 0 |
|  | BridgePRS | 0 | 0 | 0 | BridgePRS | 0 | 0 | 0 |
| SAS | JointPRS | 16 | 11 | 5 | JointPRS | 16 | 11 | 5 |
|  | XPASS | 0 | 0 | 0 | XPASS | 0 | 0 | 0 |
|  | SDPRX | 0 | 0 | 0 | SDPRX | 0 | 0 | 0 |
|  | PRS-CSx | 16 | 5 | 11 | PRS-CSx | 16 | 5 | 11 |
|  | MUSSEL | 0 | 0 | 0 | MUSSEL | 0 | 0 | 0 |
|  | PROSPER | 0 | 0 | 0 | PROSPER | 0 | 0 | 0 |
|  | BridgePRS | 0 | 0 | 0 | BridgePRS | 0 | 0 | 0 |
| AMR | JointPRS | 15 | 11 | 4 | JointPRS | 16 | 10 | 6 |
|  | XPASS | 0 | 0 | 0 | XPASS | 0 | 0 | 0 |
|  | SDPRX | 0 | 0 | 0 | SDPRX | 0 | 0 | 0 |
|  | PRS-CSx | 16 | 4 | 12 | PRS-CSx | 16 | 6 | 10 |
|  | MUSSEL | 1 | 1 | 0 | MUSSEL | 0 | 0 | 0 |
|  | PROSPER | 0 | 0 | 0 | PROSPER | 0 | 0 | 0 |
|  | BridgePRS | 0 | 0 | 0 | BridgePRS | 0 | 0 | 0 |

**Fig. 2.**
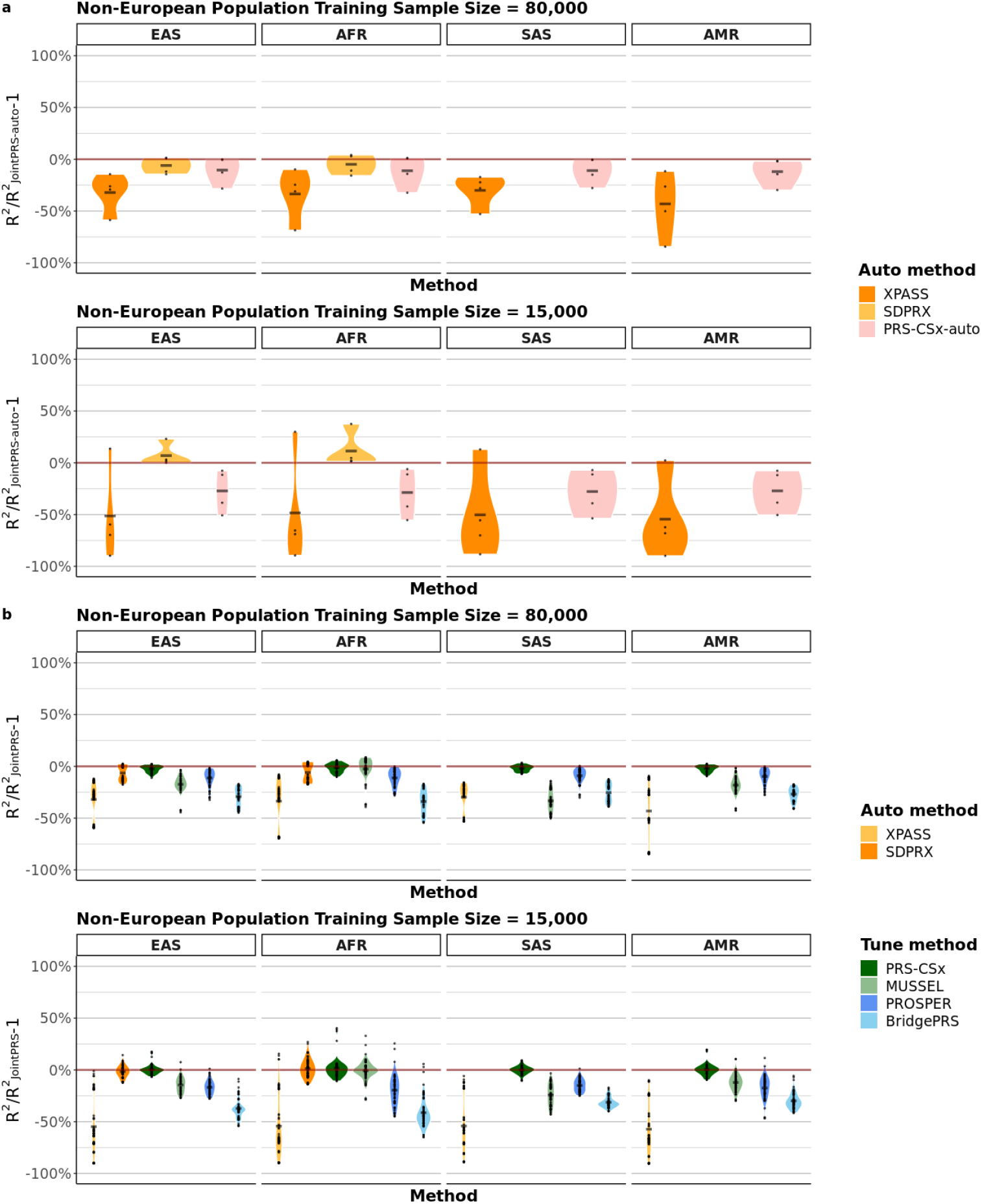
Relative Performances for Multi-population PRS Methods in Comparison to JointPRS across All Different Settings. **a-b**, The training dataset sample sizes for non-European populations were set to *N*_train_ = 80, 000 and 15, 000. The y-axis was calculated as the *R*^2^(other methods)*/R*^2^(JointPRS-auto) −1 or *R*^2^(other methods)*/R*^2^(JointPRS) −1 for each method based on the simulation setting. (**a**) represents the simulation results when there is no tuning dataset. For each method (XPASS, SDPRX, and PRS-CSx-auto) in non-European each population (EAS, AFR, SAS, and AMR), each dot represents the mean value for one causal SNPs proportion (*p* = 0.1, 0.01, 0.001, and 5×10^*−*4^) across five replicates. (**b**) represents the simulation results when there is a tuning dataset. For each method (XPASS, SDPRX, and PRS-CSx-auto) in non-European each population (EAS, AFR, SAS, and AMR), each dot represents the mean value for one causal SNPs proportion (*p* = 0.1, 0.01, 0.001, and 5 × 10^*−*4^) and one tuning dataset sample size (*N*_tune_ = 500, 2, 000, 5, 000, and 10, 000) across five replicates. Here, SDPRX cannot provide predictions for SAS and AMR as it does not provide the corresponding LD reference panels.

When a tuning dataset is not available, the results (Figure 2a and Figure 3a; Table S1) show that JointPRS-auto outperforms other auto methods (XPASS, SDPRX, and PRS-CSx-auto) with large non-European training sample sizes (*N*_train_ = 80, 000) and high genetic correlations (*ρ* = 0.8) across four causal SNP proportions (*p* = 0.1, 0.01, 0.001, and 5 × 10^*−*4^) for four non-European populations (EAS, AFR, SAS, and AMR). Additionally, JointPRS-auto ranks second among auto methods with small non-European training sample sizes (*N*_train_ = 15, 000) in two non-European populations (EAS and AFR) and first in the remaining two non-European populations (SAS and AMR) (Figure 3b; Table S1).

**Fig. 3.**
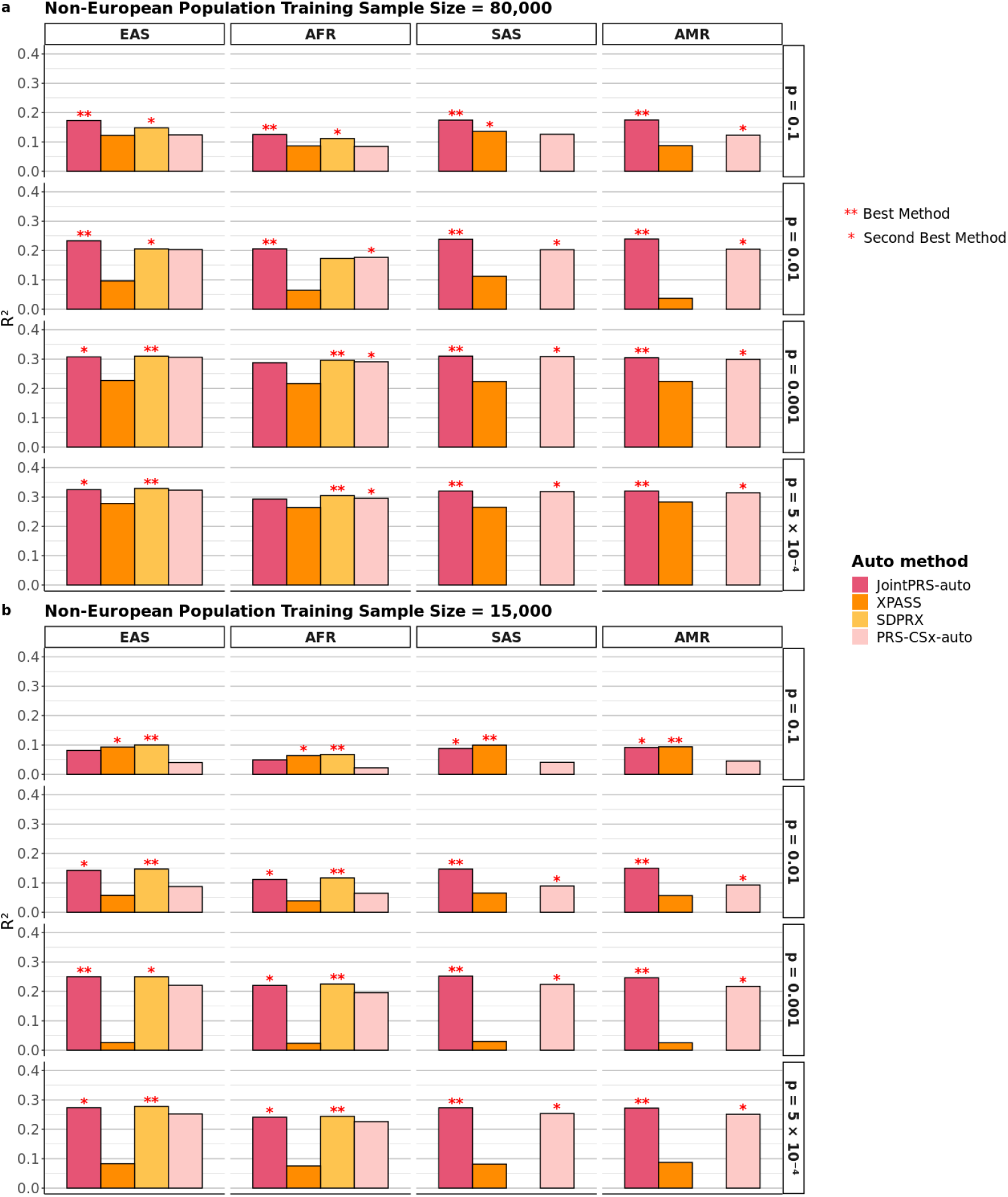
Simulation Results without a Tuning Dataset for Multi-population PRS Methods. The proportion of causal SNPs was set to *p* = 0.1, 0.01, 0.001, and 5 × 10^*−*4^. The training sample sizes for non-European populations were set to *N*_train_ = 80, 000 and 15, 000. Simulations in each scenario were repeated five times, and the mean values are presented in the bar plots for the four non-European populations (EAS, AFR, SAS, and AMR). Only four auto methods were considered here, with the best and second-best method denoted by two stars and one star, respectively, in the corresponding bar plots. Here, SDPRX cannot provide predictions for SAS and AMR as it does not provide the corresponding LD reference panels.

Specifically, XPASS exhibits the worst performance in most situations, indicating that an appropriate sparse distribution assumption is essential. Comparing JointPRS-auto with SDPRX involves a trade-off between modeling multiple populations and modeling exact genetic architecture. The significant improvement of JointPRS-auto over SDPRX with large training sample size (*N*_train_ = 80, 000) and large casual SNP proportions (*p* = 0.1, 0.001) implies that modeling multiple populations may be more helpful as non-European training sample sizes increase in highly polygenic traits. Conversely, genetic architecture modeling becomes more critical with small training sample sizes (*N*_train_ = 15, 000) or large effect SNPs (*p* = 0.001, 5 × 10^*−*4^). Furthermore, SDPRX can only be applied to EAS and AFR populations because the LD reference data are only available for these two populations in SDPRX. Lastly, as the causal SNP proportion decreases (*p* = 0.1, 0.01, 0.001, 5 × 10^*−*4^) with fixed genetic correlation (*ρ* = 0.8), the improvement of JointPRS-auto over PRS-CSx-auto decreases, suggesting that the genetic correlation structure is more helpful for highly polygenic traits.

We further investigated whether the improvement of JointPRS-auto over PRS-CSx-auto is related to the underlying cross-population genetic correlations. Five genetic correlation settings (*ρ* = 0, 0.2, 0.4, 0.6, and 0.8) were considered, with large causal SNP proportion (*p* = 0.1) and large non-European training sample size (*N*_train_ = 80, 000). Figure S9 demonstrates that the improvement of JointPRS-auto over PRS-CSx-auto increases as the genetic correlation increases.

When there is a tuning dataset available, the results (Figure 2b, Figure 4, and Figures S2 -S8; Table 2, and Tables S2 -S5) demonstrate that JointPRS has robust performance among seven methods (JointPRS, XPASS, SDPRX, PRS-CSx, MUSSEL, PROSPER, and BridgePRS) across non-European population training sample sizes (*N*_train_ = 80, 000 and 15, 000) with varying tuning sample sizes (*N*_tune_ = 500, 2, 000, 5, 000, and 10, 000), causal SNPs proportions (*p* = 0.1, 0.01, 0.001, and 5 × 10^*−*4^) when genetic correlation is 0.8 for non-European populations (EAS, AFR, SAS, AMR).

**Fig. 4.**
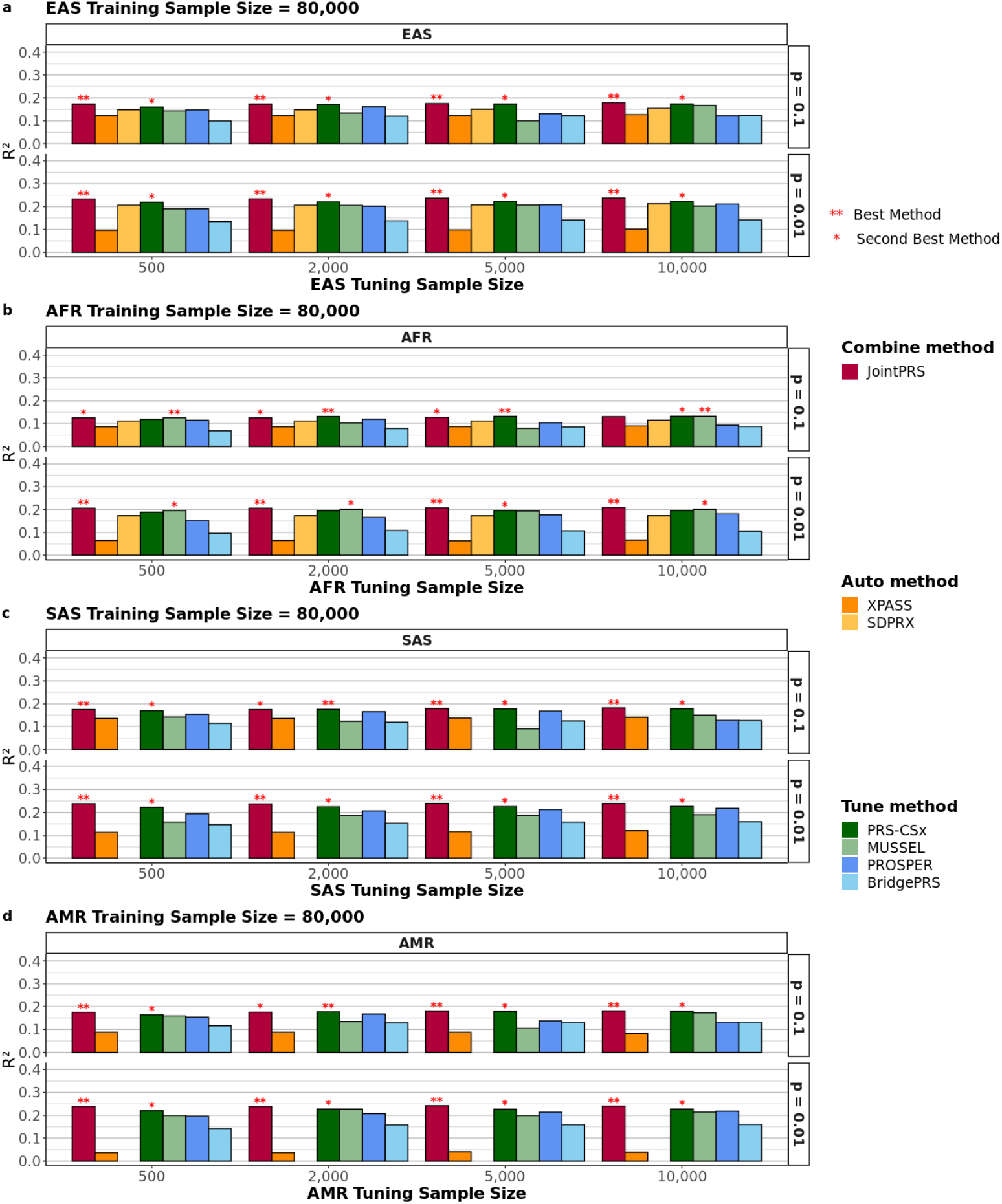
Simulation Results with Large Training Datasets Sample Sizes, Large Causal SNP Proportions, and Varying Tuning Datasets Sample Sizes for Multi-population PRS Methods. The proportion of causal SNPs was set to *p* = 0.1 and 0.01. The training sample sizes for non-European populations were set to *N*_train_ = 80, 000. The tuning sample sizes for the target population were set to *N*_tune_ = 500, 2, 000, 5, 000, and 10, 000. Simulations in each scenario were repeated five times, and the mean values are presented in the bar plots for the four non-European populations (EAS, AFR, SAS, and AMR). All methods were considered here, with the best and second-best method denoted by two stars and one star, respectively, in the corresponding bar plots. Here, SDPRX cannot provide predictions for SAS and AMR as it does not provide the corresponding LD reference panels.

Specifically, JointPRS achieves the top performance for SAS and AMR across all simulation settings when a tuning dataset is available. For EAS and AFR, JointPRS is more advantageous when the casual SNP proportion is large (*p* = 0.1 and 0.01), being the best or second-best method in 15 out of 32 simulation settings in the EAS population and 13 out of 32 simulation settings in the AFR population, indicating JointPRS performed well for highly polygenic traits. When the casual SNP proportion is small (*p* = 0.001 and 5 × 10^*−*4^), JointPRS remains advantageous for the EAS population, being the best or second-best method in 11 out of 32 simulation settings, while SDPRX and MUSSEL are more preferred in the AFR population.

Additionally, for different tuning dataset sizes (*N*_tune_ = 500, 2, 000, 5, 000 and 10, 000), Figures S5 - S8 demonstrate that JointPRS, SDPRX, PRS-CSx and BridgePRS have more stable performance than the other three methods (XPASS, MUSSEL, and PROSPER).

In conclusion, JointPRS enjoys robust performance across most simulation settings and populations, and performs better than all other methods when the non-European population training sample size is relatively large and the causal SNP proportion is large (highly polygenic traits).

### 2.3 Real Data Analysis

#### 2.3.1 Data Preparation

We collected GWAS summary statistics for 26 traits across five populations (EUR, EAS, AFR, SAS, and AMR) from various consortia. These traits were initially categorized by type (continuous or binary) and further subdivided by the number of available populations, resulting in four distinct groups.

1. **GLGC Traits**: This group includes four continuous traits—HDL-cholesterol (HDL), LDL-cholesterol (LDL), Total cholesterol (TC), and Triglycerides (logTG)—with GWAS summary statistics from five populations (EUR, EAS, AFR, SAS, and AMR).
2. **PAGE Traits:** This group comprises five continuous traits—Height, Body mass index (BMI), Systolic blood pressure (SBP), Diastolic blood pressure (DBP), and Platelet (PLT)—with GWAS summary statistics from three populations (EUR, EAS, and AFR).
3. **BBJ Traits:** This group includes 13 continuous traits—White blood cell (WBC), Neutrophil (ENU), Lymphocyte (LYM), Monocyte (MON), Eosinophil (EOS), Red blood cell (RBC), Hematocrit (HCT), Mean corpuscular hemoglobin (MCH), Mean corpuscular volume (MCV), Hemoglobin (HB), Alanine aminotransferase (ALT), Alkaline phosphatase (ALP), and *γ*-glutamyl transpeptidase (GGT)—with GWAS summary statistics available for two (EUR and EAS) populations.
4. **Binary Traits:** This group includes four binary traits—Type 2 diabetes (T2D), Breast cancer (BrC), Coronary artery disease (CAD), and Lung cancer (LuC)—with GWAS summary statistics from two or three populations (EUR, EAS, and possibly AFR).

The detailed data information for these traits is described in the following section.

##### Quality Control and Data Preparation

Quality control was conducted following the LDHub guidelines, using LDSC software to remove duplicate SNPs [38, 39]. To ensure fair comparisons for different PRS models, we restricted SNP lists for each population to those available in reference panels used across all evaluated methods. The 1000 Genomes Project served as the reference panel for all methods. Detailed GWAS summary statistics, including trait names, sample sizes and SNP numbers, are provided in Table 3 and Table S6.

**Table 3.**
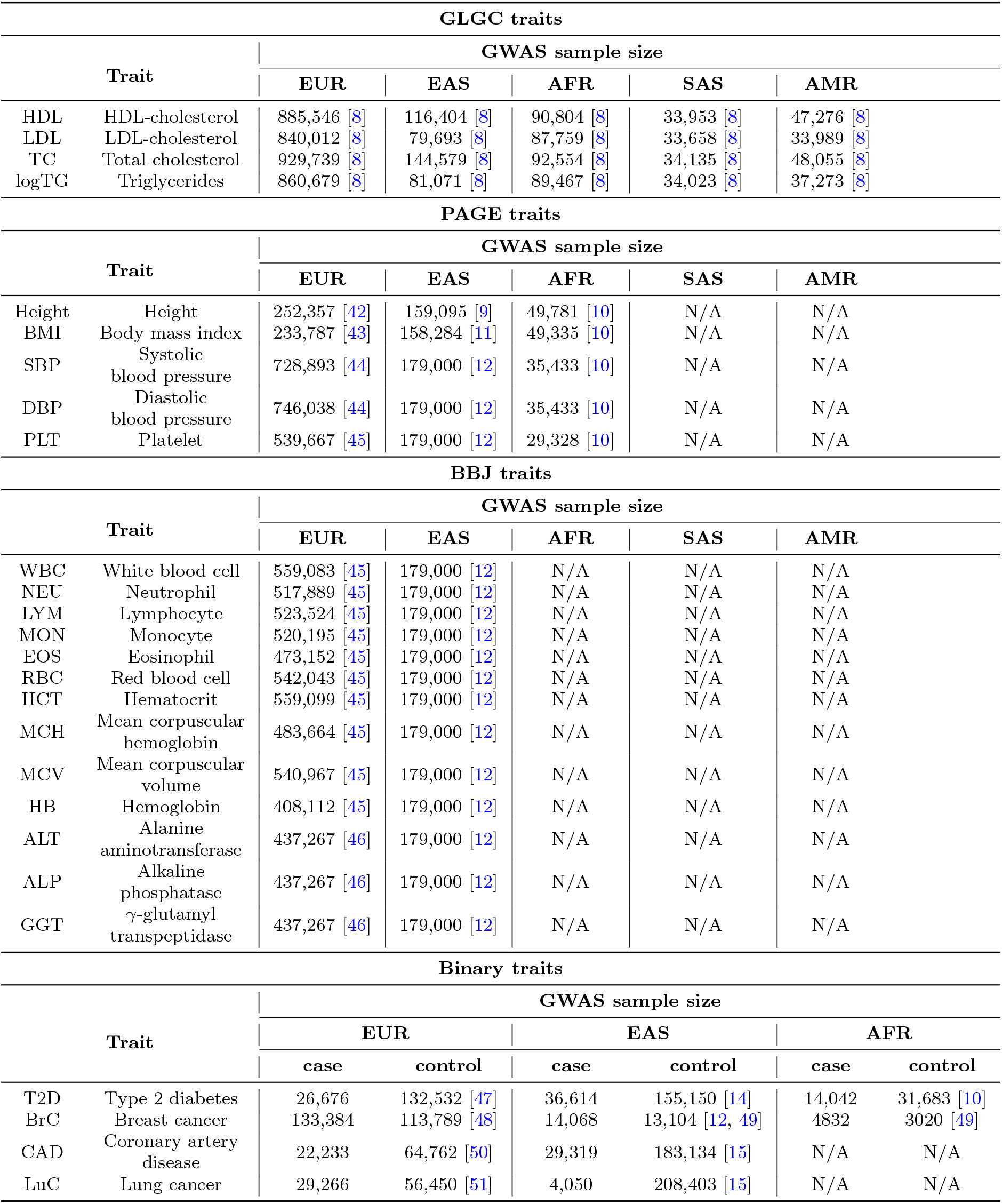
GWAS Summary Statistics Sample Size Information for 26 Traits across Five Populations.

We calculated the pairwise cross-population genetic correlation using popcorn [31] based on the GWAS summary statistics (Table S7). Our analysis reveals that the 26 traits exhibit high cross-population genetic correlations, mostly ranging from 0.7 to 0.99. However, the genetic correlations between EUR and AFR, and EAS and AFR in breast cancer, were notably lower. This discrepancy is likely attributable to the small sample size of the breast cancer AFR GWAS summary statistics.

For tuning and benchmarking multi-population methods, we utilized individual-level genetic data from the UKBB and AoU Program. Detailed sample size information is available in Tables S8 and S9, with data processing procedures including phenotype definition outlined in the Supplementary Notes (Individual-level Genetic Dataset Preparation section). Our primary goal is to assess prediction accuracy in non-European populations (EAS, AFR, SAS, and AMR). To ensure reliability, we have confirmed that there is no overlap between the training GWAS and the individual tuning and testing data from the UKBB or AoU for non-European populations. While we acknowledge the potential overlap between the EUR training GWAS and tuning and testing datasets, we do not consider this overlap to be a significant concern. Certain tune methods (MUSSEL, PROSPER, and BridgePRS) also require individual-level data for the EUR population; thus, we extracted data from 10,000 European individuals in the UKBB for these purposes.

##### Trait Analysis and Prediction Accuracy Evaluation

We analyzed 22 quantitative traits, evaluating the prediction accuracy of the derived PRS for each trait within each population. Prediction accuracy was assessed through the proportion of variance explained by the PRS, calculated as 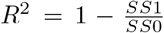, where *SS*0 and *SS*1 denote the sum of squares for residuals in the null and full models, respectively. The null model incorporated age, sex and the top 20 PCs as covariates, while the full model included the PRS and all covariates from the null model. For the four binary traits, we used a logistic model and considered the area under the curve (AUC) as the evaluation metric.

##### Method Comparison and Evaluation Scenarios

We compared the methods across all traits under three data scenarios:

1. **No tuning data:** We used the UKBB data to evaluate the performance of four auto methods: JointPRS-auto, XPASS, SDPRX, and PRS-CSx-auto.
2. **Tuning and testing data from the same cohort:** We performed 5-fold cross-validation in the UKBB data for seven methods: JointPRS, XPASS, SDPRX, PRS-CSx, MUSSEL, PROSPER, and BridgePRS.
3. **Tuning and testing data from different cohorts:** We conducted tuning in the UKBB data and evaluation in the AoU data for the same seven methods.

The percentage of relative change of method B over A is defined as (metric_*B*_ − metric_*A*_)*/*metric_*A*_ * 100%. This metric allowed us to assess the comparative performance of different PRS methods.

#### 2.3.2 Benchmarking Multi-population PRS Methods When There are No Tuning Data

We first evaluated the performance of four auto methods (JointPRS-auto, XPASS, SDPRX, and PRS-CSx-auto) across 22 quantitative traits and four binary traits in the UKBB, focusing on four non-European populations (EAS, AFR, SAS, and AMR) in the absence of tuning data. The results (Figure 5 and Figures S10 -S13, Tables S10 and S11) demonstrate that JointPRS-auto consistently outperforms the other three methods in terms of accuracy and robustness.

**Fig. 5.**
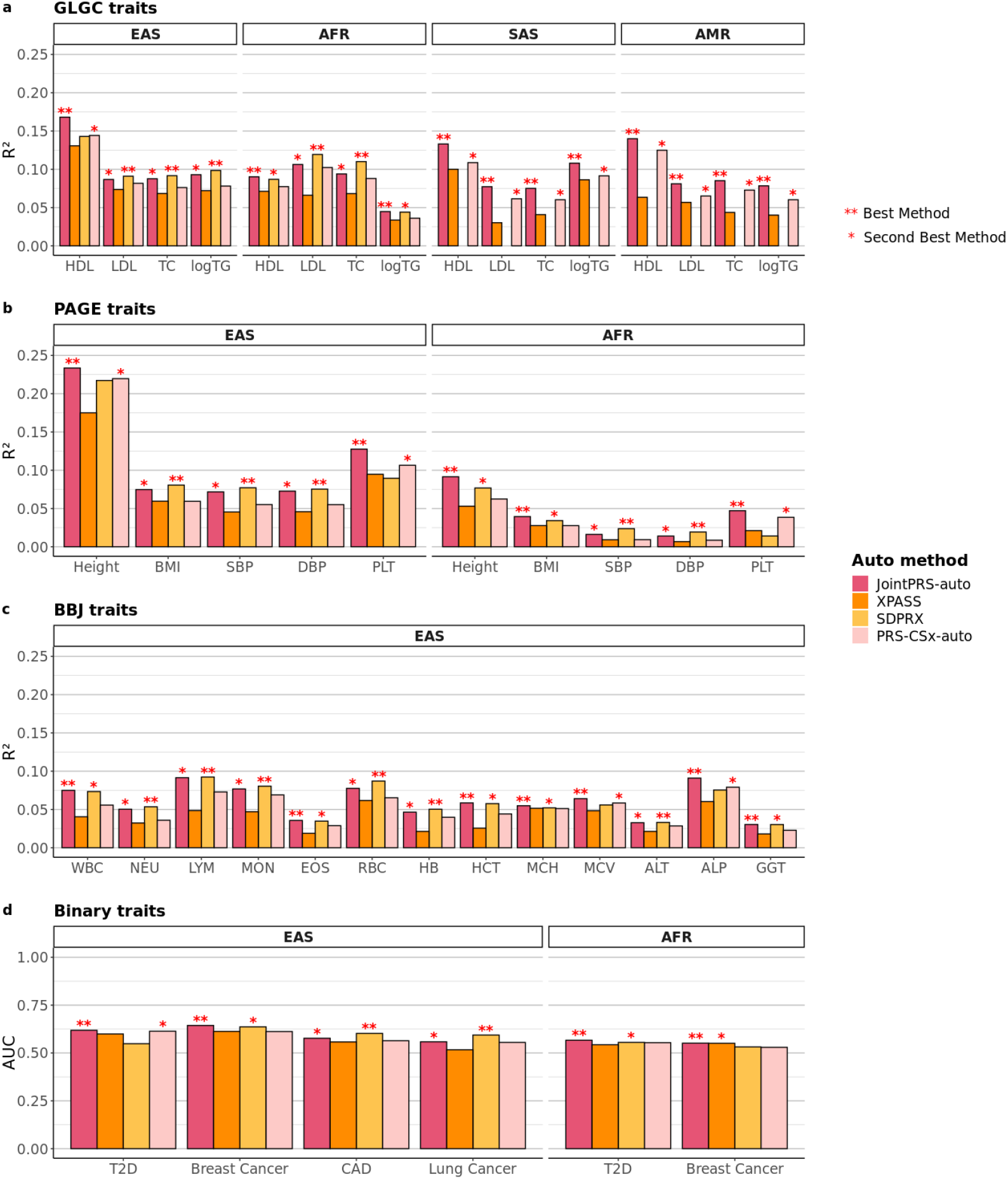
Prediction Accuracy of Multi-population PRS Methods across 26 Traits and Four Non-European Populations in UKBB When there is no Tuning Dataset. 26 traits from four categories were considered for the data scenarios when there is no tuning dataset. The *R*^2^ and AUC were used as evaluation metric for quantitative traits and binary traits and the value were presented in the bar plots for the four non-European populations (EAS, AFR, SAS, and AMR). Only four auto methods were considered here, with the best and second-best method denoted by two stars and one star, respectively, in the corresponding bar plots. Here, SDPRX cannot provide predictions for SAS and AMR as it does not provide the corresponding LD reference panels.

JointPRS-auto consistently outperforms XPASS in all four non-European populations across evaluated traits (Figure 5, Figures S10 and S11). The average improvement of JointPRS-auto over XPASS is 46.14% in the EAS population across 26 traits, 53.29% in the AFR population across 11 traits; 74.54% in the SAS population across four traits; 88.48% in the AMR population across four traits. This implies that the sparse distribution in JointPRS-auto is more appropriate.

Moreover, JointPRS-auto and SDPRX are the top performing methods in the EAS and AFR populations for many evaluated traits (Figure 5, Figures S10 and S12). However, JointPRS-auto shows more accurate and robust performance, with a 2.11% average improvement over SDPRX in the EAS population across 26 traits and a 17.86% average improvement in the AFR population across 11 traits. This indicates that JointPRS-auto achieves a balanced trade-off between modeling multiple populations and capturing complex genetic architectures, leading to more robustness across traits.

Notably, SDPRX’s application is currently limited to the EAS and AFR populations due to the availability of LD reference data provided by SDPRX.

JointPRS-auto demonstrates superior performance compared to PRS-CSx-auto across all four non-European populations for the evaluated traits (Figure 5, Figures S10 and S13). Specifically, the average improvement of JointPRS-auto over PRS-CSx-auto is 17.82% in the EAS population across 26 traits, 27.85% in the AFR population across 11 traits, 22.70% in the SAS population across four traits, and 20.94% in the AMR population across four traits. This consistent improvement indicates that the chromosome-wise cross-population genetic correlation structure utilized by JointPRS-auto enhances PRS prediction accuracy in non-European populations, particularly in the absence of tuning data.

Furthermore, we analyzed the pairwise chromosome-wise genetic correlations estimated by JointPRS-auto across 22 continuous traits (Figure S14). We observed moderate genetic correlations between the European population and the other four populations (EAS, AFR, SAS, AMR). However, contrary to our simulation results, we did not find a positive relationship between the mean chromosome-wise genetic correlation and the improvement of JointPRS-auto over PRS-CSx-auto in real data (Figure S15). This discrepancy can be attributed to various factors influencing PRS prediction performance in real-world scenarios, such as GWAS sample sizes and the distinct genetic architectures of different traits. Nonetheless, our findings suggest that JointPRS-auto consistently outperforms PRS-CSx-auto.

Moreover, we compared the performance of JointPRS-auto and PRS-CSx-auto with another continuous shrinkage prior PRS model, Xwing-auto [29], which accounts for local genetic correlations in the EAS population **(Methods)**. This analysis utilized GWAS summary statistics from the EUR and EAS populations for 22 continuous traits. The results (Figure S16) demonstrate that despite Xwing-auto incorporating more localized genetic information, JointPRS-auto consistently outperformed Xwing-auto, showing an average improvement of 19.41% across the 22 traits in the EAS population. In contrast, Xwing-auto did not perform consistently better than PRS-CSx-auto across traits, with an average improvement of only 1.53%, compared to the 21.32% improvement of JointPRS-auto over PRS-CSx-auto. These findings suggest that while incorporating local genetic correlations can be beneficial for PRS modeling, the challenge of accurately estimating these correlations limits its effectiveness. The chromosome-wise genetic correlation approach proposed by JointPRS offers a more efficient and robust method for improving multi-population PRS prediction.

In conclusion, JointPRS-auto exhibits the most robust and accurate performance across 22 quantitative traits and four binary traits evaluated in the UKBB for scenarios without tuning data, confirming the importance of appropriate sparsity distributions, incorporating chromosome-wise cross-population genetic correlations, and simultaneously modeling multiple populations for accurate non-European PRS predictions.

#### 2.3.3 Benchmarking Multi-population PRS Methods When Tuning and Testing Data Come From the Same Cohort

We evaluated the performance of seven methods—JointPRS, XPASS, SDPRX, PRS-CSx, MUSSEL, PROSPER, and BridgePRS—across 22 quantitative traits and four binary traits in four non-European populations (EAS, AFR, SAS, and AMR). In this scenario, both the tuning and testing data originated from the same cohort. The UKBB dataset was split into five folds for each populations, with each fold used for testing while the remaining folds for tuning, thus enabling a 5-fold cross-validation evaluation. The results (Figure 6, Figures S17-S23, Tables S12 and S13) illustrate that JointPRS outperforms the other methods in terms of accuracy and robustness.

**Fig. 6.**
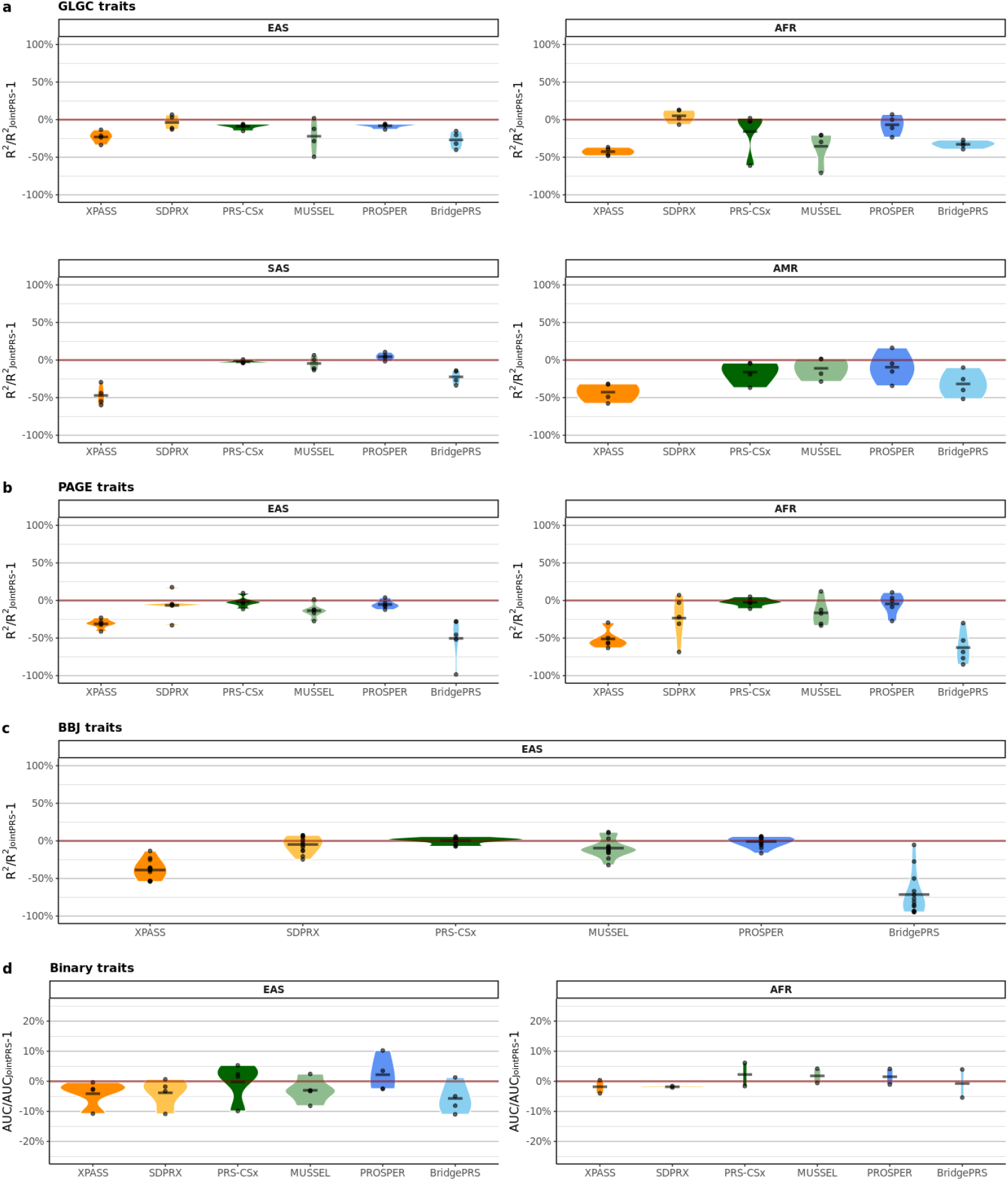
Relative Prediction Accuracy of Multi-population PRS Methods in Comparison to JointPRS across 26 Traits and Four Non-European Populations When the Tuning and Testing Data Come From the Same Cohort (UKBB). 26 traits from four categories were considered for the data scenarios when the tuning and testing data come from the same cohort (UKBB). The whole UKBB data were spilt into five folds, and we performed 5-fold cross-validation with each fold being treated as the testing data and the rest data begin treated as the tuning data. The relative change of the performance for six existing methods over that of JointPRS measured in mean of *R*^2^ and AUC across five folds were calculated for quantitative traits and binary traits. The results are presented in the violin plots for the four non-European populations (EAS, AFR, SAS, and AMR), with the mean value across traits presented as the black crossbar for each method in each trait type. Here, SDPRX cannot provide predictions for SAS and AMR as it does not provide the corresponding LD reference panels.

For GLGC traits, JointPRS and SDPRX are the top two performing methods in the EAS and AFR populations, whereas XPASS, PRS-CSx, MUSSEL, PROSPER, and BridgePRS had lower prediction accuracy. In the SAS population, JointPRS, PRS-CSx, MUSSEL, and PROSPER are the top performing methods, while SDPRX cannot provide predictions. In contrast, XPASS and BridgePRS exhibit lower prediction accuracy. In the AMR population, JointPRS consistently outperforms XPASS, PRS-CSx, MUSSEL, PROSPER, and BridgePRS, with SDPRX again providing no predictions.

Regarding the PAGE traits, JointPRS, SDPRX, PRS-CSx, and PROSPER are top performing methods for the EAS population, whereas XPASS, MUSSEL and BridgePRS show lower prediction accuracy. In the AFR population, JointPRS, PRS-CSx and PROSPER are the top performing methods, while XPASS, SDPRX, MUSSEL, and BridgePRS exhibit lower prediction accuracy.

For BBJ traits, JointPRS, SDPRX, PRS-CSx and PROSPER are the top performing methods, with XPASS, MUSSEL and BridgePRS showing lower prediction accuracy.

In the case of binary traits, all methods perform similarly, though XPASS, SDPRX, MUSSEL and BridgePRS show slightly worse performance in the EAS population.

Overall, JointPRS consistently outperformed XPASS across all populations, with average improvements of 80.58% in EAS, 89.98% in AFR, 99.98% in SAS, and 166.84% in AMR. JointPRS demonstrated better accuracy and robustness over SDPRX in the majority of evaluated traits, with average improvements of 9.18% in EAS and 29.10% in AFR populations. Compared to PRS-CSx, JointPRS showed average improvements of 2.78% in EAS, 25.02% in AFR, 2.57% in SAS, and 27.31% in AMR populations. When compared to MUSSEL, JointPRS had average improvements of 20.15% in EAS, 47.83% in AFR, 5.98% in SAS, and 84.91% in AMR. JointPRS outperformed PROSPER with average improvements of 6.30% in EAS, 7.82% in AFR, and 9.73% in AMR, and PROSPER failed to converge for the lung cancer trait in some cross-validation folds. Finally, JointPRS demonstrates consistent superiority over BridgePRS, with average improvements of 605.46% in EAS, 291.07% in AFR, 31.06% in SAS, and 60.84% in AMR.

To summarize, JointPRS consistently demonstrates better performance compared to XPASS and BridgePRS. Methods such as SDPRX, PRS-CSx, MUSSEL, and PROSPER can exhibit top performance similar to JointPRS for certain traits and populations but perform considerably worse than JointPRS in others. Notably, tune methods tend to be less accurate in GLGC traits, while auto methods tend to be less accurate in PAGE traits.

We also summarize the selection results of JointPRS versions, the prediction accuracy of the meta and tune versions, and the relative improvement achieved by the data-adaptive approach over these two versions (Figures S24-S28; Table S14). Joint-PRS selects different versions (tune or meta) depending on the trait, population, and tuning data, suggesting context-specific benefits of each version (Table S14). Notably, our data-adaptive approach often identifies the more accurate version between the meta and tune versions, especially when there is a substantial difference in their prediction accuracies (Figures S25-S28). Additionally, the positive relative improvement of the selected version over the meta and tune versions underscores the robustness of the data-adaptive approach, as evidenced by the presence of extreme positive and the well-bounded negative dots in the plot (Figure S24). These findings are consistent with our previous observations that while existing tune and auto methods can perform well for specific traits, JointPRS demonstrates robust performance across all traits due to its data-adaptive approach.

In conclusion, JointPRS exhibits the most robust and accurate performance across 22 quantitative traits and four binary traits evaluated in the UKBB when the tuning and testing data come from the same cohort. This superior performance is attributable to its data-adaptive approach, which effectively integrates the strengths of both meta-analysis and tuning strategies.

#### 2.3.4 Benchmarking Multi-population PRS Methods When Tuning and Testing Data Come From Different Cohorts

In this section, we study the performance of seven polygenic risk score (PRS) methods—JointPRS, XPASS, SDPRX, PRS-CSx, MUSSEL, PROSPER, and BridgePRS—across nine quantitative traits in two non-European populations (AFR and AMR). We consider the case where the tuning and testing data come from different cohorts: the UKBB dataset was used for tuning, while the AoU dataset was used for testing. The results (Figure 7, Figures S29-S35, Table S15) demonstrate that JointPRS outperforms the other methods in terms of both accuracy and robustness.

**Fig. 7.**
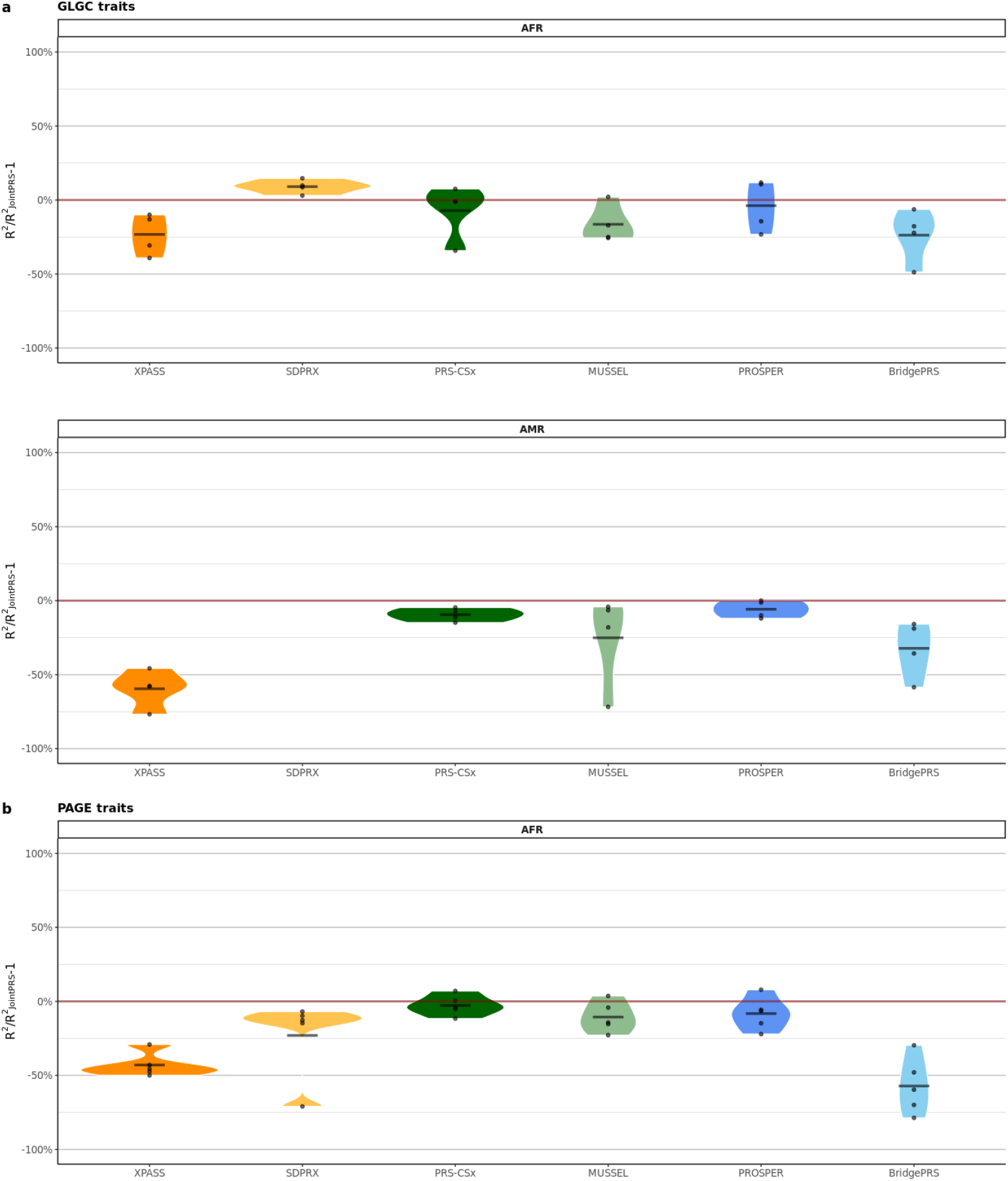
Relative Prediction Accuracy of Multi-population PRS Methods in Comparison to JointPRS across Nine Traits and Two Non-European Populations When the Tuning and Testing Data Come From Different Cohorts (UKBB and AoU). Nine traits from two categories were considered for the data scenarios when the tuning and testing data come from different cohorts (UKBB and AoU). The UKBB data were used as a tuning dataset, while the AoU data were used as a testing dataset. The relative change of the performance for six existing methods over that of JointPRS measured in *R*^2^ were calculated for quantitative traits. The results are presented in the violin plots for the two non-European populations (AFR and AMR), with the mean value across traits presented as the black crossbar for each method in each trait type. Here, SDPRX cannot provide predictions for SAS and AMR as it does not provide the corresponding LD reference panels.

For GLGC traits, JointPRS and SDPRX are the top two performing methods for the AFR population, whereas XPASS, PRS-CSx, MUSSEL, PROSPER, and BridgePRS have lower prediction accuracy. For the AMR population, JointPRS consistently outperforms XPASS, PRS-CSx, MUSSEL, PROSPER, and BridgePRS, with SDPRX failing to provide predictions. Regarding the PAGE traits, JointPRS and PRS-CSx are the top two performing methods, while XPASS, SDPRX, MUSSEL, PROSPER, and BridgePRS exhibit lower prediction accuracy.

Overall, JointPRS consistently outperformed XPASS across both AFR and AMR populations, with average improvements of 58.29% in AFR and 172.00% in AMR. JointPRS outperformed SDPRX with an average improvement of 29.14% in the AFR population. Against PRS-CSx, JointPRS showed average improvements of 7.03% in AFR and 10.55% in AMR. Compared to MUSSEL, JointPRS demonstrated average improvements of 16.85% in AFR and 71.72% in AMR. JointPRS also outperformed PROSPER in most traits, with average improvements of 8.71% in AFR and 6.46% in AMR. Finally, JointPRS consistently surpassed BridgePRS, with average improvements of 115.31% in AFR and 59.57% in AMR populations.

To summarize, XPASS and BridgePRS consistently perform worse than JointPRS. MUSSEL also perform worse than JointPRS in most traits. While SDPRX, PRS-CSx, and PROSPER exhibit top performances similar to JointPRS in some traits, JointPRS remains the only method with robust and accurate predictions across all traits and populations. Notably, tune methods tend to be less accurate for GLGC traits, whereas auto methods tend to be less accurate for PAGE traits, similar to the conclusions for the scenario where the tuning and testing data come from the same cohort.

It is also worth noting that prediction accuracy decreases when the tuning and testing data come from different cohorts. This scenario, often neglected in previous studies, should be emphasized to prevent overestimation of PRS prediction accuracy based on results where tuning and testing data are from the same cohort. However, the relative performance of different methods remains similar between these two scenarios. In conclusion, JointPRS exhibits robust and accurate performance across various traits and populations in all three data scenarios, suggesting it as a promising method for multi-population PRS prediction.

## 3 Discussion

In this manuscript, we have introduced JointPRS, an efficient approach for accurate multi-population PRS construction, applicable even when only GWAS summary statistics are available. JointPRS integrates a continuous shrinkage model, incorporates a genetic correlation structure, and simultaneously models multiple populations. When a tuning dataset is available, it employs a data-adaptive approach, effectively combining the advantages of both meta-analysis and tuning strategies. Notably, despite its additional modeling of genetic correlation structures and data-adaptive approaches compared to PRS-CSx, the computational time of JointPRS is similar to PRS-CSx (Table S16).

JointPRS was benchmarked against six state-of-the-art multi-population PRS methods: PRS-CSx [19], PROSPER [23], MUSSEL [22], SDPRX [20], XPASS [17], and BridgePRS [24]. This benchmarking was conducted across three real data scenarios: no tuning dataset (UKBB), tuning and testing data from the same cohort (UKBB), and tuning and testing data from different cohorts (UKBB and AoU), covering 22 quantitative traits and four binary traits in four non-European populations (EAS, AFR, SAS, and AMR).

Consistent with previous studies [21–23], our results show that no single method performs uniformly best across all traits and data scenarios. This highlights the critical importance of assessing the robust accuracy of each method.

In the absence of a tuning dataset, JointPRS-auto and SDPRX exhibited superior performance across various traits, with JointPRS-auto being more robust and accurate based on the average improvement over SDPRX. We further elucidated the advantages of the structural features within JointPRS-auto, including appropriate sparse distribution, cross-population genetic correlation, and multiple-population modeling. These benefits were validated through comparisons with XPASS, PRS-CSx-auto, and SDPRX. Additionally, comparisons of JointPRS-auto, PRS-CSx-auto, and Xwing-auto [29] show that the chromosome-wise genetic correlation structure in JointPRS-auto is advantageous to the no-correlation structure in PRS-CSx-auto and the local genetic correlation structure in Xwing-auto.

When a tuning dataset is available, JointPRS consistently demonstrates better or comparable performance across all traits, outperforming other methods that are less robust and accurate. This robust accuracy is attributed to the data-adaptive approach of JointPRS, which allows it to optimally select between meta-analysis and tuning strategies based on the available tuning data. Additionally, we observed that when the tuning and testing data come from different cohorts, the prediction accuracy of all PRS methods decreases compared to when they come from the same cohort. This finding implies that for different PRS application scenarios, reliance solely on prediction accuracy from the same cohort may be overly optimistic.

We also evaluated all seven methods across extensive simulation settings, including varying training sample sizes, different tuning dataset scenarios (no tuning data and tuning data with varying sample sizes), varying causal SNP proportions, and varying cross-population genetic correlations. Across all simulation settings, JointPRS exhibited robust and accurate performance, particularly excelling with large training sample sizes, high causal SNP proportions, and large genetic correlations. Additionally, JointPRS demonstrated robustness to varying tuning data sample sizes.

Despite its advantages, JointPRS has limitations. We have focused on Hapmap3 SNPs due to the LD reference panels used, aligning with practices in PRS-CSx, SDPRX, and BridgePRS [19, 20, 24]. Incorporating additional SNPs, as suggested by studies like CT-SLEB, MUSSEL, and PROSPER [21–23], could potentially enhance prediction accuracy. Moreover, a disparity in prediction accuracy between European and non-European populations persists. This gap cannot be bridged solely by sophisticated statistical models and requires larger GWAS sample sizes for non-European populations [6].

Additionally, further exploration of additional techniques or information in PRS modeling will also be valuable, such as the transfer learning techniques proposed by TL-Multi and TL-PRS [26, 28], cross-trait information considered by XPXP [18], and fine-mapping information incorporated in PolyPred [27]. However, these approaches may present additional challenges, and further investigations are required to evaluate their benefits.

In conclusion, we have proposed a robust and accurate method for incorporating GWAS summary statistics with possible tuning data. Our simulation and real data studies highlight crucial structures in PRS modeling and factors in real data that influence the accuracy and robustness of multi-population PRS methods. JointPRS stands out as a versatile and reliable method, suitable for diverse genetic prediction scenarios.

## 4 Method

Our proposed JointPRS framework comprises two key steps: [1] model development and estimation, and [2] data-adaptive approach implementation. These steps are elaborated in the following subsections, along with an additional section providing an overview of existing methods.

### 4.1 JointPRS Model Development and Estimation

In JointPRS model, we assume that the SNP effect size *β*_*j*_ for SNP *j* across *K* populations is modeled with a correlated Gaussian prior when SNP *j* is available for all *K* populations:

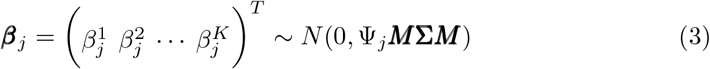

with

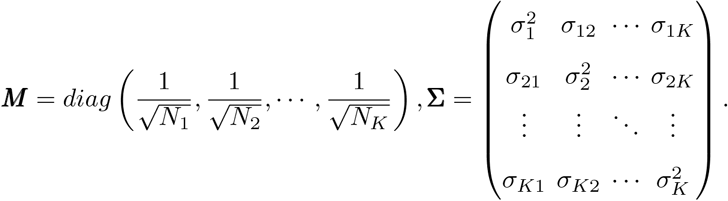

Here, we can further express the covariance matrix **Σ** as the transformation of the correlation matrix

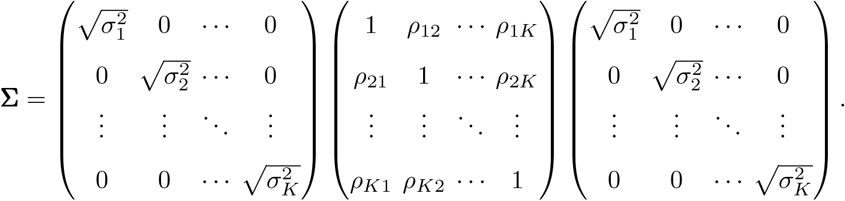

Here 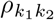 denotes the cross-population genetic correlation between populations *k*_1_ and *k*_2_. Here, *k*_1_, *k*_2_ ∈ {EUR, EAS, AFR, SAS, AMR}, *k*_1_ ≠ *k*_2_. In addition, Ψ_*j*_ is the sample-size normalized effect size for SNP *j* shared across *K* populations. This shared effect Ψ_*j*_ is modeled by a continuous shrinkage prior following the PRScsx model and assumed to follow the gamma distribution 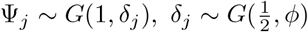[19]. Here the global shrinkage parameter *ϕ* is dependent on the JointPRS version we choose.

For SNP *j* available only in a subset of populations, this model is truncated to include only populations that contain SNP *j*.

Therefore, we can use the following index matrix ***T*** to encode the SNP missing patterns across the whole genome for different populations:

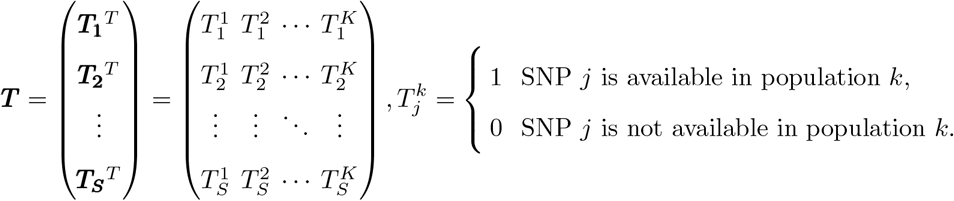

Each SNP *j* is assumed to follow a *sum*(***T***_***j***_)-dimension multivariate correlated Gaussian model, with the truncated covariance matrix keeping only rows and columns corresponding to 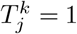 for *k* = 1, …, *K*.

The residual component ***ϵ***^*k*^ is assumed to follow a Gaussian distribution, denoted by 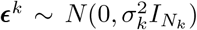. The variance component 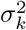 here is assumed to follow a non-informative Jeffreys prior with its density distribution expressed as 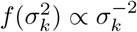. We note that the 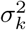 are also the diagonal elements of the covariance matrix, thus avoiding the convergence issue. Since different populations do not share samples, we can assume the independence of ***ϵ***^*k*^ across populations. Moreover, since the correlation matrix is symmetric, we only assume all upper triangle elements in the correlation matrix: *ρ*_12_, *ρ*_13_, · · ·, *ρ*_*K−*1*K*_ to follow a uniform distribution *Uniform*(0, 1) and estimate each correlation pair automatically from the GWAS summary statistics.

The following sections provide a detailed illustration of the JointPRS versions, JointPRS data preparation, and JointPRS parameter estimation procedures.

#### 4.1.1 JointPRS Versions

##### Auto Version

The auto version exclusively utilizes GWAS summary statistics and does not require any parameter tuning. Specifically, the input data consist of the original GWAS summary statistics, with the global shrinkage parameter *ϕ* assumed to follow a standard half-Cauchy prior,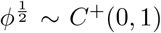. This version streamlines the process by eliminating the need for additional tuning data or parameter adjustments.

##### Meta Version

The meta version leverages meta-analysis to integrate the original GWAS summary statistics with tuning datasets, utilizing the METAL software [40]. This results in updated GWAS summary statistics as input data. Similar to the auto version, the meta version assumes the global shrinkage parameter *ϕ* follows a standard half-Cauchy prior,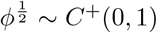. The meta version does not require parameter tuning while integrating the tuning data through GWAS summary statistics updates.

##### Tune Version

In contrast to the meta version, the tune version uses the original GWAS summary statistics as input data and chooses the global shrinkage parameter *ϕ* from a broader range: 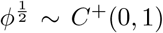 and the set *ϕ* ∈ {10^*−*6^, 10^*−*4^, 10^*−*2^, 1} using the tuning data. More specifically, this version performs a linear combination across populations for each candidate *ϕ*, selecting the optimal *ϕ* to derive the optimal weights. This approach ensures the model is finely tuned to achieve accurate performance for each population based on the tuning data.

#### 4.1.2 Data Preparation in JointPRS Model

We first need to obtain the GWAS summary statistics, which are the marginal least squares effect size estimates for *S* SNPS in *K* populations 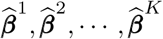.

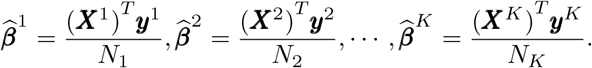

Here, ***y***^*k*^, ***X***^*k*^ and *N*_*k*_ denote the standardized phenotype vector, the column-standardized genetic matrix in population *k*, and the GWAS summary statistic sample size, respectively. Note that some SNP data might be missing in each population, and this issue will be carefully addressed in the subsequent algorithm.

We also need the LD matrix 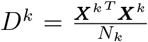 for each population *k*, utilizing either the data from the 1000 Genomes Project phase 3 [34] or the UKBB [32], as adopted from PRS-CSx [19]. Due to computational efficiency, the entire genome will be partitioned into *L*^*k*^ independent LD blocks based on the reference data for each population *k*. During each MCMC iteration, SNP effect sizes are updated sequentially for each population *k* within each LD block *l*_*k*_. We simplify the notation *l*^*k*^ to *l* during the updates, representing the block variable in the current updating population *k*.

In the current updating population *k*, 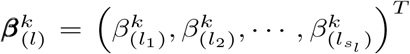 represents the effect size vectors for SNPs in block *l* of population *k*, with *s*_*l*_ indicating the number of SNPs in block *l* of population *k*. The marginal effect size estimates for SNPs in block *l* across *K* populations are denoted by 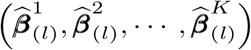. Additionally, 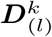 denotes the LD matrix for SNPs in block *l* of population *k*, whereas 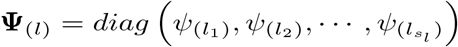 represents the shrinkage matrix for SNPs in block *l*, and the covariance matrix for SNPs in block *l* of population *k* are denoted by 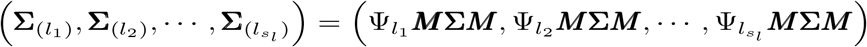.

#### 4.1.3 MCMC and MH Algorithm in JointPRS Model

We use 1000×*K* MCMC iterations with the first 500×*K* steps as burn-in as suggested by PRS-CSx [19]. For each iteration step, we update parameters by the following procedure:

1. Update 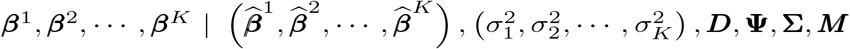: For the current updating population *k* in block *l*, we update the posterior effect size by the following:

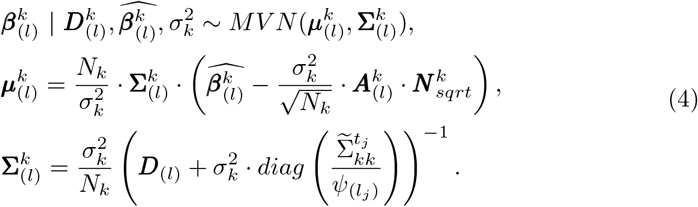

Here,

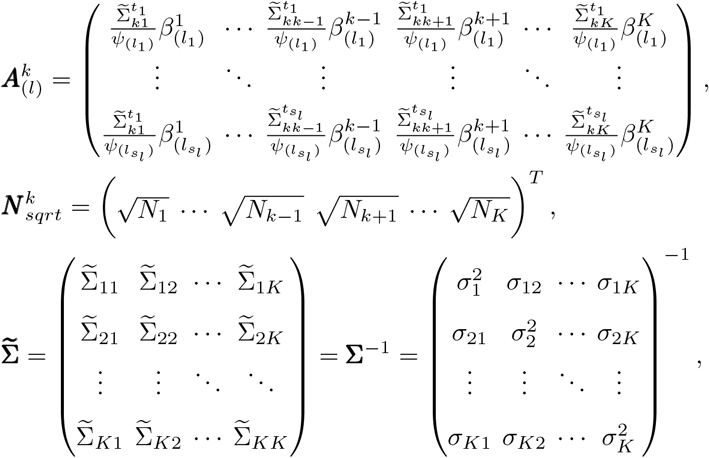

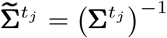 the inverse of the covariance matrix for non-missing populations in SNP *j*.
2. Update 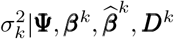: For the current updating population *k*, we update the variance by the following:

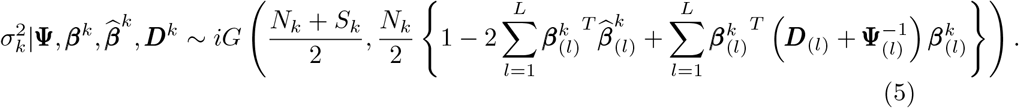

Here *iG*(*α, β*) is the inverse-gamma distribution with the probability density function

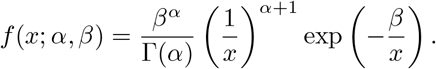
3. Update each pair of correlation 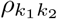 from the upper triangle under different constraint. Based on the joint model we propose, we can estimate the cross-population correlation 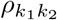 for two populations *k*_1_ and *k*_2_ (*k*_1_ ∈ {EUR, EAS, AFR, SAS, AMR}, *k*_2_ ∈ {EUR, EAS, AFR, SAS, AMR}) by assuming:

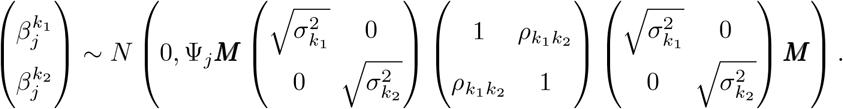

We note that when we update 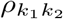, we only use SNPs shared by populations *k*_1_ and *k*_2_. And if we denote the posterior distribution of 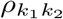 as 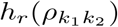, we have

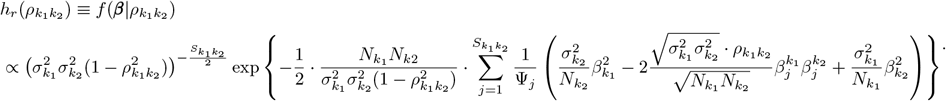

Here 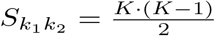 is a constant representing the total pairs of cross-population genetic correlation. Since there is no closed-form distribution to update the correlation 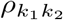, we use the following MH algorithm:

#### Algorithm 1

MH Algorithm for JointPRS

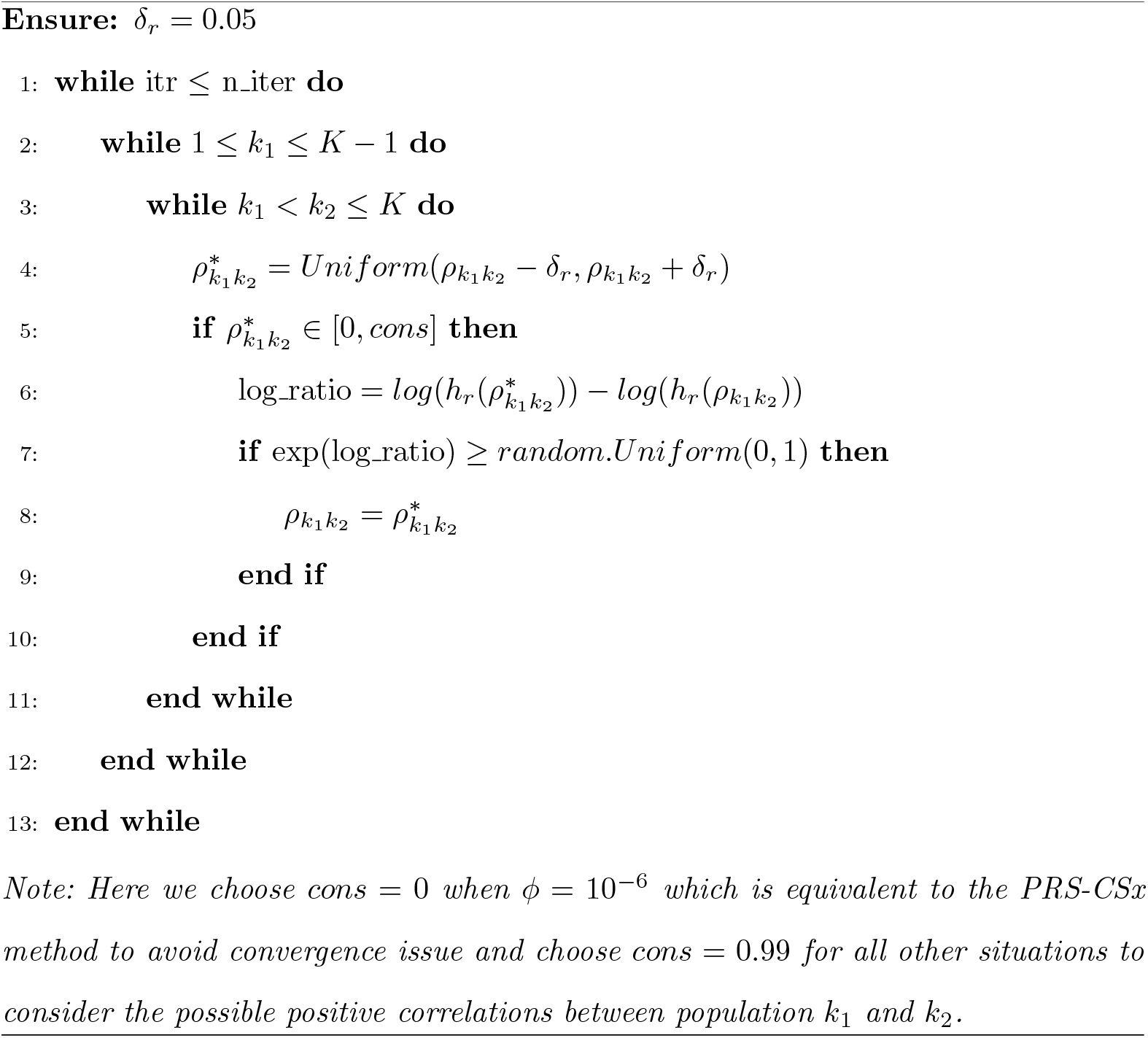 Then we can also update the corresponding covariance pair 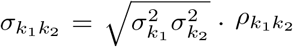.
4. Update Ψ_*j*_ | ***β***_*j*_, *σ*^2^, *δ*_*j*_: For each SNP *j*, we update the corresponding shrinkage parameter by the following:

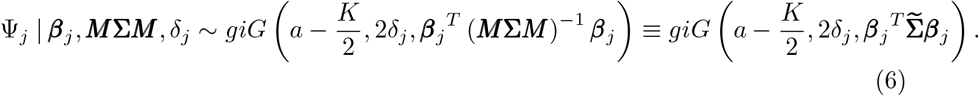

Here *giG*(*λ, ρ, χ*) is the three-parameter generalized inverse Gaussian distribution with the probability density function

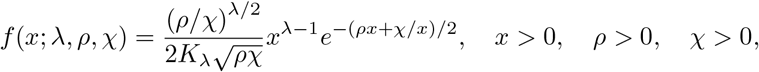

where *K*_*λ*_ is the modified Bessel function of the second kind.
5. Update *δ*_*j*_ | Ψ_*j*_: For each SNP *j*, we update the distribution parameter by the following:

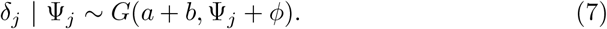

Here *G*(*c, d*) is the Gamma distribution with the probability density function

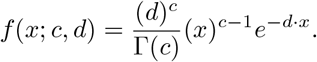

The detailed proof for the above updating procedure can be found in the Supplementary Notes (MCMC and MH algorithm in JointPRS Section).

### 4.2 JointPRS Data-Adaptive Approach Implementation

When a tuning dataset is available, JointPRS employs a data adaptive approach to select the PRS between two versions: the meta version and the tune version.

#### Meta Version

For the meta version in JointPRS, we utilized the SNP effect size estimates, denoted as 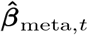, for the target population *t*. This estimate was applied to the tuning dataset to obtain the PRS estimation result:

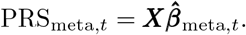

To address overfitting issues noted in Section 2.1, we use 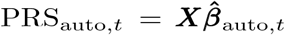 to approximate the performance of PRS_meta,*t*_. Here, 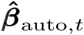 is the SNP effect estimate from the auto version for the target population *t*.

#### Tune Version

For the tune version, SNP effect size estimates,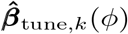, were obtained for each of the *K* populations (*k* = 1, 2, · · · *K*). These estimates were applied to the tuning dataset to generate *K* PRS estimation results for each *ϕ* ∈ {10^*−*6^, 10^*−*4^, 10^*−*2^, 1, *auto*} as follows:

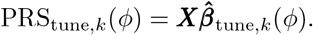

For convenience in model comparison, we set *ϕ* = auto and use PRS_tune,*k*_(auto) ≡ PRS_auto,*k*_ to approximate the tune version PRS for each population *k*.

#### Model Comparison

After establishing the two models and their approximation to avoid overfitting, we compare the performance of the two approximated versions on the tuning data using the F-test. Specifically, the model comparison between the tune version and the meta version approximately tests whether the full model is statistically significant compared to the submodel, as follows:

#### Regression Setting with Continuous Traits

**Submodel (meta version): *Y*** = PRS_auto,*t*_ · *α* + *ϵ*, given some *t* ∈ [1, *K*],

**Full model (tune version):** 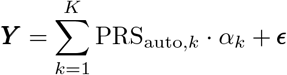.

We use residual sum of squares (RSS) from standard linear regression of the above two models (*RSS*_auto,*t*_ and *RSS*_auto,1:*K*_, respectively) to evaluate the reduction in RSS:

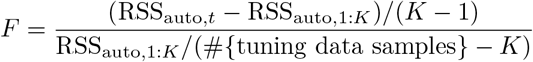

The right-tail p-value of this F-test statistic is assessed with *F* (*K* − 1, #{ tuning data samples} − *K*), using 0.05 as the p-value threshold. The full model is accepted only when the p-value is less than 0.05.

#### Regression Setting with Binary Traits

**Submodel (meta version): *Y*** ∼ Bernoulli(*p*), 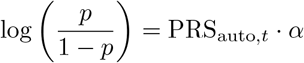, given some *t* ∈ [1, *K*],

**Full model (tune version): *Y*** ∼ Bernoulli(*p*),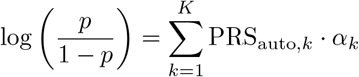.

Maximum likelihood estimates (MLE) are obtained for the above two logistic regression models. We then calculate the *χ*^2^-test statistic:

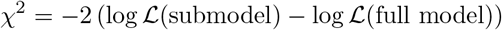

The right-tail p-value of this *χ*^2^-test statistic is assessed with *χ*^2^(*K* − 1), using 0.05 as the p-value threshold. The full model is accepted only when the p-value is less than 0.05. Here,logℒ(submodel) represents the log-likelihood of the submodel, and logℒ(full model) represents the log-likelihood of the full model.

### 4.3 Overview of Existing Methods

#### XPASS

The XPASS method [17] jointly integrates GWAS summary statistics and LD structures from two populations through a bivariate Gaussian distribution. The genetic correlation structure is incorporated into the model to facilitate the information transference from the auxiliary to the target population. The following model illustrate its idea for the prior on the effect size of each SNP *j* for population 1 and population 2.

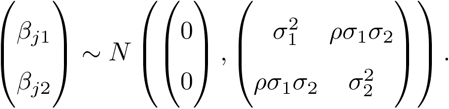

This method is further augmented by considering population-specific effects using selected SNPs based on the P+T procedure, treating them as fixed effects during the estimation. Although this method does not need tuning parameters, when a tuning dataset is available, a linear combination of scores from both populations will be performed to derive the final score for the target population in our analysis.

#### SDPRX

The SDPRX method [20] establishes a hierarchical Bayesian framework, jointly modeling GWAS summary statistics and LD structures from two populations. This framework comprises four components to characterize the genetic architecture, identifying SNPs as no effect, being population-specific, or being shared across two populations. The essential component of this method is the shared component, which uses a mixture of bivariate Gaussian distributions, coupled with a precalculated genetic correlation to approximate the true shared structure. The following equation represent the prior on the effect sizes for each SNP *j* for population 1 and population 2.

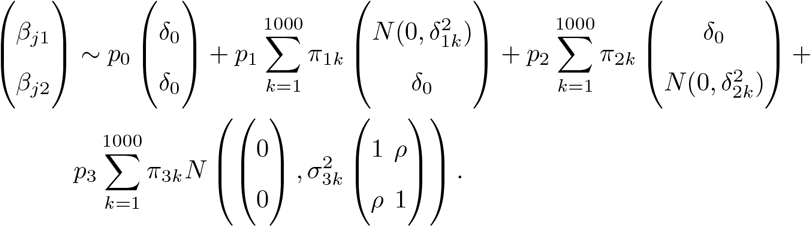

Although this method does not need tuning parameters, in the presence of a tuning dataset, a linear combination of scores from both populations will be performed to obtain the final score for the target population in our analysis.

#### PRS-CSx(-auto)

The PRS-CSx model [19], an extended Bayesian model of the PRS-CS framework [41], integrates GWAS summary statistics and LD structures from multiple populations by utilizing a shared continuous shrinkage prior for each SNP *j* in population *k* as the following equation.

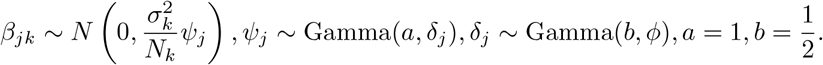

In scenarios when a tuning dataset is unavailable, the model leverages a full Bayesian approach to estimate the global shrinkage parameter *ϕ*^0.5^ ∼ *C*^+^(0, 1). However, when a tuning dataset is available, PRS-CSx evaluates a predefined set of global shrinkage parameters *ϕ* ∈ {10^*−*6^, 10^*−*4^, 10^*−*2^, 1, *auto*}. For each parameter in the set, the obtained scores for each population will be linearly combined to obtain the final score for each population. This integration relies on the tuning dataset, and the shrinkage parameter that leads to the best performed combined scores will then be selected based on the prediction accuracy of the tuning dataset.

#### MUSSEL

The MUSSEL method [22] jointly models GWAS summary statistics and LD structures from multiple populations, utilizing the following multivariate spikeand-slab prior with an incorporated genetic correlation structure for the effect size of each SNP *j* in each block (*J*).

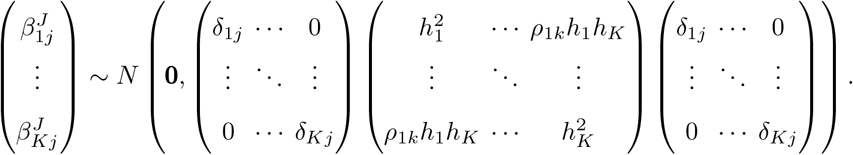

This method requires a tuning dataset due to the presence of two sets of tuning parameters within the model: the causal SNP proportion and heritability in each population 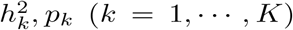 and the between-population correlation *ρ*_*ij*_ (*i*≠*j, i* = 1, · · ·, *K, j* = 1, · · ·, *K*). Additionally, a super learning step, which selects linear methods from Lasso, Ridge, Elastic net, and Linear regression, is introduced to further integrate the scores from various populations and tuning parameters.

#### PROSPER

The PROSPER method [23] integrates GWAS summary statistics and LD structures from multiple populations, leveraging a linear regression with a combination of Lasso and ridge penalties for estimation in order to consider the spraity of genetic effect sizes and the similarity across populations. The objective function to optimize the effect size vectors of SNP *i* in *K* populations can be represented by the following equation

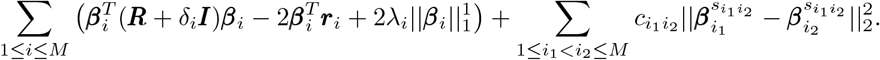

This method requires a tuning dataset to determine the tuning parameters associated with these penalties 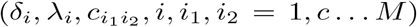. A further ensemble step (also called super learning, which selects linear methods from Lasso, Ridge, Elastic net, and Linear regression) is implemented to combine PRS scores generated across different penalty parameters and populations.

#### BridgePRS

The BridgePRS method [24] integrates GWAS summary statistics from two populations to consider shared and population-specific SNP effects for the target population. In the first stage, it models the SNP effect sizes for each population under the following zero-centered Gaussian prior

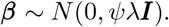

In the second stage, it utilizes the auxiliary population (population 1) to determine the prior of the target population under the following Gaussian model

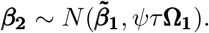

And they finally linearly combine the PRS from the target population with the above joint PRS based on the ridge regression fit.

#### XWing(-auto)

XWing method [29] integrates GWAS summary statistics and LD information from multiple populations, incorporating portable genetic effects identified through quantifying local genetic correlation between populations with continuous shrinkage prior to construct cross-population PRS. The algorithm consists of three main steps: Firstly, Xwing detects local genetic correlations between a target population and an auxiliary population using a scan statistic approach. Secondly, Xwing incorporates the identified correlated regions as an annotation into for the SNP effect sizes modeling:

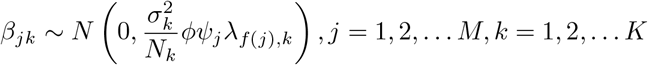

Here, *λ*_*f*(*j*),*k*_ is the annotation-dependent shrinkage parameter that allows differential regularization based on whether the SNP is annotated or not.

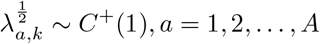

The global shrinkage parameter *ϕ* was set to *ϕ* ∈ {10^*−*6^, 10^*−*4^, 10^*−*2^, 1, *auto*} Population-specific PRS models are generated for each target-auxiliary population pairing. Finally, X-Wing combines all generated PRS models via an innovative repeated learning strategy to estimate optimal linear combination weights.

## Code Availability

### JointPRS Method

JointPRS, https://github.com/LeqiXu/JointPRS

### Existing Methods

XPASS, https://github.com/YangLabHKUST/XPASS

SDPRX, https://github.com/eldronzhou/SDPRX

PRS-CSx, https://github.com/getian107/PRScsx

MUSSEL, https://github.com/Jin93/MUSSEL

PROSPER, https://github.com/Jingning-Zhang/PROSPER

BridgePRS, https://github.com/clivehoggart/BridgePRS

## Data Availability

BBJ summary statistics, http://jenger.riken.jp/en/result, https://humandbs.biosciencedbc.jp/en/hum0197-v3-220

BCAC summary statistics, http://bcac.ccge.medschl.cam.ac.uk/bcacdata/, https://www.ebi.ac.uk/gwas/publications/29059683

BCX summary statistics, http://www.mhi-humangenetics.org/en/resources/

CARDIoGRAM summary statistics, https://www.ebi.ac.uk/gwas/publications/21378990

TRICL-ILCCO and LC3 summary statistics, https://www.ebi.ac.uk/gwas/publications/28604730

DIAGRAM summary statistics, https://www.ebi.ac.uk/gwas/publications/28566273

GIANT summary statistics, https://portals.broadinstitute.org/collaboration/giant/index.php/GIANT_consortium_data_files

GLGC summary statistics, https://csg.sph.umich.edu/willer/public/glgc-lipids2021/

ICBP summary statistics, https://www.ebi.ac.uk/gwas/publications/30224653

PAGE summary statistics, https://www.ebi.ac.uk/gwas/publications/31217584

UKBB summary statistics, https://www.ebi.ac.uk/gwas/publications/33972514

## Acknowledgments

This research was supported in part by NIH grant R01 HG012735 and NSF grant DMS 2310836. We thank Jiaqi Hu, Dr. Chi Zhang, and Dr. Yingxin Lin for their invaluable discussions. We are also grateful to Dr. Tian Ge, Dr. Jingning Zhang, Dr. Jin Jin, Jiacheng Miao, and Dr. Hanmin Guo for sharing their codes, LD reference panels, and for their suggestions on implementing various methods.

Our research utilized the UK Biobank resource under approved data request (refs: 29900) and the All of Us resource. We gratefully acknowledge UK Biobank and All of Us participants for their contributions, without whom this research would not have been possible. We also thank the National Institutes of Health’s All of Us Research Program for making available the participant data examined in this study.

## References

[1] Khera, A. V. et al. Genome-wide polygenic scores for common diseases identify individuals with risk equivalent to monogenic mutations. Nature Genetics 50, 1219–1224 (2018).

[2] Seibert, T. M. et al. Polygenic hazard score to guide screening for aggressive prostate cancer: development and validation in large scale cohorts. BMJ 360 (2018).

[3] Abraham, G., Rutten-Jacobs, L. & Inouye, M. Risk prediction using polygenic risk scores for prevention of stroke and other cardiovascular diseases. Stroke 52, 2983–2991 (2021).

[4] Dehestani, M., Liu, H. & Gasser, T. Polygenic risk scores contribute to personalized medicine of parkinson’s disease. Journal of Personalized Medicine 11, 1030 (2021).

[5] Tam, V. et al. Benefits and limitations of genome-wide association studies. Nature Reviews Genetics 20, 467–484 (2019).

[6] Martin, A. R. et al. Clinical use of current polygenic risk scores may exacerbate health disparities. Nature Genetics 51, 584–591 (2019).

[7] Lewis, C. M. & Vassos, E. Polygenic risk scores: from research tools to clinical instruments. Genome Medicine 12, 1–11 (2020).

[8] Graham, S. E. et al. The power of genetic diversity in genome-wide association studies of lipids. Nature 600, 675–679 (2021).

[9] Akiyama, M. et al. Characterizing rare and low-frequency height-associated variants in the japanese population. Nature Communications 10, 4393 (2019).

[10] Wojcik, G. L. et al. Genetic analyses of diverse populations improves discovery for complex traits. Nature 570, 514–518 (2019).

[11] Akiyama, M. et al. Genome-wide association study identifies 112 new loci for body mass index in the japanese population. Nature Genetics 49, 1458–1467 (2017).

[12] Sakaue, S. et al. A cross-population atlas of genetic associations for 220 human phenotypes. Nature Genetics 53, 1415–1424 (2021).

[13] Chen, M.-H. et al. Trans-ethnic and ancestry-specific blood-cell genetics in 746,667 individuals from 5 global populations. Cell 182, 1198–1213 (2020).

[14] Suzuki, K. et al. Identification of 28 new susceptibility loci for type 2 diabetes in the japanese population. Nature Genetics 51, 379–386 (2019).

[15] Ishigaki, K. et al. Large-scale genome-wide association study in a japanese population identifies novel susceptibility loci across different diseases. Nature Genetics 52, 669–679 (2020).

[16] Jia, G. et al. Genome-wide association analyses of breast cancer in women of african ancestry identify new susceptibility loci and improve risk prediction. Nature genetics 1–8 (2024).

[17] Cai, M. et al. A unified framework for cross-population trait prediction by leveraging the genetic correlation of polygenic traits. The American Journal of Human Genetics 108, 632–655 (2021).

[18] Xiao, J. et al. Xpxp: improving polygenic prediction by cross-population and cross-phenotype analysis. Bioinformatics 38, 1947–1955 (2022).

[19] Ruan, Y. et al. Improving polygenic prediction in ancestrally diverse populations. Nature Genetics 54, 573–580 (2022).

[20] Zhou, G., Chen, T. & Zhao, H. Sdprx: A statistical method for cross-population prediction of complex traits. The American Journal of Human Genetics 110, 13–22 (2023).

[21] Zhang, H. et al. A new method for multiancestry polygenic prediction improves performance across diverse populations. Nature genetics 55, 1757–1768 (2023).

[22] Jin, J. et al. Mussel: Enhanced bayesian polygenic risk prediction leveraging information across multiple ancestry groups. Cell Genomics 4 (2024).

[23] Zhang, J. et al. An ensemble penalized regression method for multi-ancestry polygenic risk prediction. Nature Communications 15, 3238 (2024).

[24] Hoggart, C. J. et al. Bridgeprs leverages shared genetic effects across ancestries to increase polygenic risk score portability. Nature Genetics 56, 180–186 (2024).

[25] Amariuta, T. et al. Improving the trans-ancestry portability of polygenic risk scores by prioritizing variants in predicted cell-type-specific regulatory elements. Nature Genetics 52, 1346–1354 (2020).

[26] Tian, P. et al. Multiethnic polygenic risk prediction in diverse populations through transfer learning. Frontiers in Genetics 13, 906965 (2022).

[27] Weissbrod, O. et al. Leveraging fine-mapping and multipopulation training data to improve cross-population polygenic risk scores. Nature Genetics 54, 450–458 (2022).

[28] Zhao, Z., Fritsche, L. G., Smith, J. A., Mukherjee, B. & Lee, S. The construction of cross-population polygenic risk scores using transfer learning. The American Journal of Human Genetics 109, 1998–2008 (2022).

[29] Miao, J. et al. Quantifying portable genetic effects and improving cross-ancestry genetic prediction with gwas summary statistics. Nature Communications 14, 832 (2023).

[30] Mostafavi, H. et al. Variable prediction accuracy of polygenic scores within an ancestry group. elife 9, e48376 (2020).

[31] Brown, B. C., Ye, C. J., Price, A. L. & Zaitlen, N. Transethnic genetic-correlation estimates from summary statistics. The American Journal of Human Genetics 99, 76–88 (2016).

[32] Sudlow, C. et al. Uk biobank: an open access resource for identifying the causes of a wide range of complex diseases of middle and old age. PLoS medicine 12, e1001779 (2015).

[33] of Us Research Program Investigators, A. The “all of us” research program. New England Journal of Medicine 381, 668–676 (2019).

[34] Consortium,. G. P. et al. A global reference for human genetic variation. Nature 526, 68 (2015).

[35] Consortium, I. H. et al. Integrating common and rare genetic variation in diverse human populations. Nature 467, 52 (2010).

[36] Yang, J., Lee, S. H., Goddard, M. E. & Visscher, P. M. Gcta: a tool for genomewide complex trait analysis. The American Journal of Human Genetics 88, 76–82 (2011).

[37] Chang, C. C. et al. Second-generation plink: rising to the challenge of larger and richer datasets. Gigascience 4, s13742–015 (2015).

[38] Zheng, J. et al. Ld hub: a centralized database and web interface to perform ld score regression that maximizes the potential of summary level gwas data for snp heritability and genetic correlation analysis. Bioinformatics 33, 272–279 (2017).

[39] Bulik-Sullivan, B. K. et al. Ld score regression distinguishes confounding from polygenicity in genome-wide association studies. Nature Genetics 47, 291–295 (2015).

[40] Willer, C. J., Li, Y. & Abecasis, G. R. Metal: fast and efficient meta-analysis of genomewide association scans. Bioinformatics 26, 2190–2191 (2010).

[41] Ge, T., Chen, C.-Y., Ni, Y., Feng, Y.-C. A. & Smoller, J. W. Polygenic prediction via bayesian regression and continuous shrinkage priors. Nature Communications 10, 1776 (2019).

[42] Wood, A. R. et al. Defining the role of common variation in the genomic and biological architecture of adult human height. Nature Genetics 46, 1173–1186 (2014).

[43] Locke, A. E. et al. Genetic studies of body mass index yield new insights for obesity biology. Nature 518, 197–206 (2015).

[44] Evangelou, E. et al. Genetic analysis of over 1 million people identifies 535 new loci associated with blood pressure traits. Nature Genetics 50, 1412–1425 (2018).

[45] Vuckovic, D. et al. The polygenic and monogenic basis of blood traits and diseases. Cell 182, 1214–1231 (2020).

[46] Pazoki, R. et al. Genetic analysis in european ancestry individuals identifies 517 loci associated with liver enzymes. Nature Communications 12, 2579 (2021).

[47] Scott, R. A. et al. An expanded genome-wide association study of type 2 diabetes in europeans. Diabetes 66, 2888–2902 (2017).

[48] Zhang, H. et al. Genome-wide association study identifies 32 novel breast cancer susceptibility loci from overall and subtype-specific analyses. Nature genetics 52, 572–581 (2020).

[49] Michailidou, K. et al. Association analysis identifies 65 new breast cancer risk loci. Nature 551, 92–94 (2017).

[50] Schunkert, H. et al. Large-scale association analysis identifies 13 new susceptibility loci for coronary artery disease. Nature Genetics 43, 333–338 (2011).

[51] McKay, J. D. et al. Large-scale association analysis identifies new lung cancer susceptibility loci and heterogeneity in genetic susceptibility across histological subtypes. Nature Genetics 49, 1126–1132 (2017).

