## Supplement 1 for "JointPRS: A Data-Adaptive Framework for Multi-Population Genetic Risk Prediction Incorporating Genetic Correlation"

### Supplementary Notes

#### MCMC and MH Algorithm in JointPRS

In this section, we illustrate the estimation procedure involving the Markov Chain Monte Carlo (MCMC) and Metropolis-Hastings (MH) algorithms within the JointPRS model. We use $1000 \times K$ MCMC iterations with the first $500 \times K$ steps as burn-in as suggested by PRS-CSx [1]. For each iteration step, we update parameters by the following procedure:

1. Update $\boldsymbol{\beta}^1, \boldsymbol{\beta}^2, \cdots, \boldsymbol{\beta}^K \mid \left(\widehat{\boldsymbol{\beta}}^1, \widehat{\boldsymbol{\beta}}^2, \cdots, \widehat{\boldsymbol{\beta}}^K\right), (\sigma_1^2, \sigma_2^2, \cdots, \sigma_K^2), \boldsymbol{D}, \boldsymbol{\Psi}, \boldsymbol{\Sigma}, \boldsymbol{M}$:

   For the current updating population $k$ in block $l$, we update the posterior effect size by the following:

$$\begin{aligned}
&\boldsymbol{\beta}_{(l)}^k \mid \boldsymbol{D}_{(l)}^k, \widehat{\boldsymbol{\beta}_{(l)}}^k, \sigma_k^2 \sim MVN(\boldsymbol{\mu}_{(l)}^k, \boldsymbol{\Sigma}_{(l)}^k), \\
&\boldsymbol{\mu}_{(l)}^k = \frac{N_k}{\sigma_k^2} \cdot \boldsymbol{\Sigma}_{(l)}^k \cdot \left(\widehat{\boldsymbol{\beta}_{(l)}}^k - \frac{\sigma_k^2}{\sqrt{N_k}} \cdot \boldsymbol{A}_{(l)}^k \cdot \boldsymbol{N}_{sqrt}^k\right), \\
&\boldsymbol{\Sigma}_{(l)}^k = \frac{\sigma_k^2}{N_k} \left(\boldsymbol{D}_{(l)} + \sigma_k^2 \cdot diag\left(\frac{\widetilde{\Sigma}_{kk}^{t_j}}{\psi_{(l_j)}}\right)\right)^{-1}.
\end{aligned} \tag{1}$$

   Here,

$$\boldsymbol{A}_{(l)}^k = \begin{pmatrix} \frac{\widetilde{\Sigma}_{k1}^{t_1}}{\psi_{(l_1)}}\beta_{(l_1)}^1 & \cdots & \frac{\widetilde{\Sigma}_{kk-1}^{t_1}}{\psi_{(l_1)}}\beta_{(l_1)}^{k-1} & \frac{\widetilde{\Sigma}_{kk+1}^{t_1}}{\psi_{(l_1)}}\beta_{(l_1)}^{k+1} & \cdots & \frac{\widetilde{\Sigma}_{kK}^{t_1}}{\psi_{(l_1)}}\beta_{(l_1)}^{K} \\ \vdots & \ddots & \vdots & \vdots & \ddots & \vdots \\ \frac{\widetilde{\Sigma}_{k1}^{t_1}}{\psi_{(l_{s_l})}}\beta_{(l_{s_l})}^1 & \cdots & \frac{\widetilde{\Sigma}_{kk-1}^{t_{s_l}}}{\psi_{(l_{s_l})}}\beta_{(l_{s_l})}^{k-1} & \frac{\widetilde{\Sigma}_{kk+1}^{t_{s_l}}}{\psi_{(l_{s_l})}}\beta_{(l_{s_l})}^{k+1} & \cdots & \frac{\widetilde{\Sigma}_{kK}^{t_{s_l}}}{\psi_{(l_{s_l})}}\beta_{(l_{s_l})}^{K} \end{pmatrix},$$

$$\boldsymbol{N}_{sqrt}^k = \begin{pmatrix} \sqrt{N_1} & \cdots & \sqrt{N_{k-1}} & \sqrt{N_{k+1}} & \cdots & \sqrt{N_K} \end{pmatrix}^T,$$

$$\widetilde{\boldsymbol{\Sigma}} = \begin{pmatrix} \widetilde{\Sigma}_{11} & \widetilde{\Sigma}_{12} & \cdots & \widetilde{\Sigma}_{1K} \\ \widetilde{\Sigma}_{21} & \widetilde{\Sigma}_{22} & \cdots & \widetilde{\Sigma}_{2K} \\ \vdots & \vdots & \ddots & \ddots \\ \widetilde{\Sigma}_{K1} & \widetilde{\Sigma}_{K2} & \cdots & \widetilde{\Sigma}_{KK} \end{pmatrix} = \boldsymbol{\Sigma}^{-1} = \begin{pmatrix} \sigma_1^2 & \sigma_{12} & \cdots & \sigma_{1K} \\ \sigma_{21} & \sigma_2^2 & \cdots & \sigma_{2K} \\ \vdots & \vdots & \ddots & \vdots \\ \sigma_{K1} & \sigma_{K2} & \cdots & \sigma_K^2 \end{pmatrix}^{-1},$$

   $\widetilde{\boldsymbol{\Sigma}}^{t_j} = \left(\boldsymbol{\Sigma}^{t_j}\right)^{-1}$ the inverse of the covariance matrix for non-missing populations in SNP $j$.

   *Proof.*

$$
\begin{aligned}
& f\left(\boldsymbol{\beta} \mid \boldsymbol{y}, \boldsymbol{X},\left(\sigma_1^2, \sigma_2^2, \cdots, \sigma_K^2\right), \boldsymbol{D}, \boldsymbol{\Psi}, \boldsymbol{\Sigma}, \boldsymbol{M}\right) \\
& \propto f\left(\boldsymbol{y} \mid \boldsymbol{\beta}, \boldsymbol{X},\left(\sigma_1^2, \sigma_2^2, \cdots, \sigma_K^2\right)\right) \cdot f(\boldsymbol{\beta} \mid \left(\boldsymbol{\Sigma}_1, \boldsymbol{\Sigma}_2, \cdots, \boldsymbol{\Sigma}_S\right)) \\
& \propto \exp \left(-\frac{1}{2} \sum_{k=1}^K \frac{\left\|\boldsymbol{y}^k-\boldsymbol{X}^k \boldsymbol{\beta}^k\right\|_2^2}{\sigma_k^2}\right) \cdot \exp \left(-\frac{1}{2} \sum_{j=1}^S \boldsymbol{\beta}_j^T \boldsymbol{\Sigma}_j^{-1} \boldsymbol{\beta}_j\right) \\
& \propto \exp \left(-\frac{1}{2} \sum_{k=1}^K \frac{\left\|\boldsymbol{y}^k-\sum_{l=1}^L \boldsymbol{X}_{(l)}^k \boldsymbol{\beta}_{(l)}^k\right\|_2^2}{\sigma_k^2}\right) \cdot \exp \left(-\frac{1}{2} \sum_{l=1}^L \sum_{j=1}^{s_l} \boldsymbol{\beta}_{(l_j)}^T \boldsymbol{\Sigma}_{(l_j)}^{-1} \boldsymbol{\beta}_{(l_j)}\right) \\
& \propto \exp \left(-\frac{1}{2} \sum_{l=1}^L\left(\sum_{k=1}^K\left(\boldsymbol{\beta}_{(l)}^k{}^T \frac{\boldsymbol{X}_{(l)}^k{}^T \boldsymbol{X}_{(l)}^k}{\sigma_k^2} \boldsymbol{\beta}_{(l)}^k-2 \boldsymbol{\beta}_{(l)}^k{}^T \frac{\boldsymbol{X}_{(l)}^k{}^T \boldsymbol{y}^k}{\sigma_k^2}\right)+\sum_{j=1}^{s_l} \boldsymbol{\beta}_{(l_j)}^T \boldsymbol{\Sigma}_{(l_j)}^{-1} \boldsymbol{\beta}_{(l_j)}\right)\right) .
\end{aligned}
$$

For each block $l,\ l=1,2,\cdots,L$ :

$$
\begin{aligned}
& f\left(\boldsymbol{\beta}_{(l)} \mid \boldsymbol{y}, \boldsymbol{X}_{(l)},\left(\sigma_1^2, \sigma_2^2, \cdots, \sigma_k^2\right),\left(\boldsymbol{\Sigma}_{(l_1)}, \boldsymbol{\Sigma}_{(l_2)}, \cdots, \boldsymbol{\Sigma}_{(l_{s_l})}\right)\right) \\
& \quad \propto \exp \left(-\frac{1}{2}\left(\sum_{k=1}^K\left(\boldsymbol{\beta}_{(l)}^k{}^T \frac{\boldsymbol{X}_{(l)}^k{}^T \boldsymbol{X}_{(l)}^k}{\sigma_k^2} \boldsymbol{\beta}_{(l)}^k-2 \boldsymbol{\beta}_{(l)}^k{}^T \frac{\boldsymbol{X}_{(l)}^k{}^T \boldsymbol{y}^k}{\sigma_k^2}\right)+\sum_{j=1}^{s_l} \boldsymbol{\beta}_{(l_j)}^T \boldsymbol{\Sigma}_{(l_j)}^{-1} \boldsymbol{\beta}_{(l_j)}\right)\right) .
\end{aligned}
$$

For population $k,\ k=1,2,\cdots,K$, we fix other populations value. So

$$
\begin{aligned}
& f\left(\boldsymbol{\beta}_{(l)}^k | \boldsymbol{y}^k, \boldsymbol{X}_{(l)}^k, \sigma_k^2,\left(\boldsymbol{\Sigma}_{(l_1)}, \boldsymbol{\Sigma}_{(l_2)}, \cdots, \boldsymbol{\Sigma}_{(l_{s_l})}\right), \boldsymbol{\beta}_{(l)}^{-k}\right) \\
& \propto \exp \left(-\frac{1}{2}\left(\boldsymbol{\beta}_{(l)}^k{}^T \frac{\boldsymbol{X}_{(l)}^k{}^T \boldsymbol{X}_{(l)}^k}{\sigma_k^2} \boldsymbol{\beta}_{(l)}^k-2 \boldsymbol{\beta}_{(l)}^k{}^T \frac{\boldsymbol{X}_{(l)}^k{}^T \boldsymbol{y}^k}{\sigma_k^2}+\sum_{j=1}^{s_l} \boldsymbol{\beta}_{(l_j)}^T \boldsymbol{\Sigma}_{(l_j)}^{-1} \boldsymbol{\beta}_{(l_j)}\right)\right) \\
& \propto \exp \left(-\frac{1}{2}\left(\boldsymbol{\beta}_{(l)}^k{}^T \frac{\boldsymbol{X}_{(l)}^k{}^T \boldsymbol{X}_{(l)}^k}{\sigma_k^2} \boldsymbol{\beta}_{(l)}^k-2 \boldsymbol{\beta}_{(l)}^k{}^T \frac{\boldsymbol{X}_{(l)}^k{}^T \boldsymbol{y}^k}{\sigma_k^2}+\sum_{j=1}^{s_l} \frac{1}{\psi_{(l_j)}} \cdot \boldsymbol{\beta}_{(l_j)}^T \boldsymbol{M}^{-1} \widetilde{\boldsymbol{\Sigma}} \boldsymbol{M}^{-1} \boldsymbol{\beta}_{(l_j)}\right)\right) .
\end{aligned}
$$

Here, different SNPs have different missing patterns, denoted as $t_j$ for SNP $j$. And for notation simplicity, we modify $\widetilde{\boldsymbol{\Sigma}}$ to $\widetilde{\boldsymbol{\Sigma}}^{t_j}$ to remove populations without SNP $j$, while maintaining other notations unchanged. However, we note that all notations involving SNP $j$ must omit the elements corresponding to the population missing SNP $j$. Then we have

$$\sum_{j=1}^{s_l} \frac{1}{\psi_{(l_j)}} \cdot \boldsymbol{\beta}_{(l_j)}^T \boldsymbol{M}^{-1} \widetilde{\boldsymbol{\Sigma}}^{t_j} \boldsymbol{M}^{-1} \boldsymbol{\beta}_{(l_j)} \propto \sum_{j=1}^{s_l} \frac{1}{\psi_{(l_j)}} \cdot \left( \widetilde{\Sigma}_{kk}^{t_j} \cdot \beta_{(l_j)}^{k}{}^2 \cdot N_k + 2\beta_{(l_j)}^k \cdot \sqrt{N_k} \sum_{k_1 \neq k} \widetilde{\Sigma}_{kk_1}^{t_j} \beta_{(l_j)}^{k_1} \sqrt{N_{k_1}} \right)$$

$$= \sum_{j=1}^{s_l} \beta_{(l_j)}^{k}{}^2 \cdot \frac{\widetilde{\Sigma}_{kk}^{t_j}}{\psi_{(l_j)}} \cdot N_k + 2 \cdot \sqrt{N_k} \cdot \beta_{(l_j)}^k \left( \frac{\widetilde{\Sigma}_{k1}^{t_j}}{\psi_{(l_j)}} \beta_{(l_j)}^1 \cdots \frac{\widetilde{\Sigma}_{kk-1}^{t_j}}{\psi_{(l_j)}} \beta_{(l_j)}^{k-1} \quad \frac{\widetilde{\Sigma}_{kk+1}^{t_j}}{\psi_{(l_j)}} \beta_{(l_j)}^{k+1} \cdots \frac{\widetilde{\Sigma}_{kK}^{t_j}}{\psi_{(l_j)}} \beta_{(l_j)}^{K} \right) \begin{pmatrix} \sqrt{N_1} \\ \vdots \\ \sqrt{N_{k-1}} \\ \sqrt{N_{k+2}} \\ \vdots \\ \sqrt{N_K} \end{pmatrix}$$

$$\equiv N_k \cdot \boldsymbol{\beta}_{(l)}^{k}{}^T \left( diag \left( \frac{\widetilde{\Sigma}_{kk}^{t_j}}{\psi_{(l_j)}} \right) \right) \boldsymbol{\beta}_{(l)}^k + 2 \cdot \sqrt{N_k} \cdot \boldsymbol{\beta}_{(l)}^{k}{}^T \boldsymbol{A}_{(l)}^k \boldsymbol{N}_{sqrt}^k.$$

Then we have

$$f\left(\boldsymbol{\beta}_{(l)}^k | \boldsymbol{y}^k, \boldsymbol{X}_{(l)}^k, \sigma_k^2, \left(\boldsymbol{\Sigma}_{(l_1)}, \boldsymbol{\Sigma}_{(l_2)}, \cdots, \boldsymbol{\Sigma}_{(l_{s_l})}\right), \boldsymbol{\beta}_{(l)}^{-k}\right)$$

$$\propto \exp\left( -\frac{1}{2} \left( \boldsymbol{\beta}_{(l)}^{k}{}^T \frac{\boldsymbol{X}_{(l)}^{k}{}^T \boldsymbol{X}_{(l)}^k}{\sigma_k^2} \boldsymbol{\beta}_{(l)}^k - 2\boldsymbol{\beta}_{(l)}^{k}{}^T \frac{\boldsymbol{X}_{(l)}^{k}{}^T \boldsymbol{y}^k}{\sigma_k^2} + N_k \cdot \boldsymbol{\beta}_{(l)}^{k}{}^T \left( diag \left( \frac{\widetilde{\Sigma}_{kk}^{t_j}}{\psi_{(l_j)}} \right) \right) \boldsymbol{\beta}_{(l)}^k + 2 \cdot \sqrt{N_k} \cdot \boldsymbol{\beta}_{(l)}^{k}{}^T \boldsymbol{A}_{(l)}^k \boldsymbol{N}_{sqrt}^k \right) \right)$$

$$\propto \exp\left( -\frac{N_k}{2\sigma_k^2} \left( \boldsymbol{\beta}_{(l)}^{k}{}^T \frac{\boldsymbol{X}_{(l)}^{k}{}^T \boldsymbol{X}_{(l)}^k}{N_k} \boldsymbol{\beta}_{(l)}^k - 2\boldsymbol{\beta}_{(l)}^{k}{}^T \frac{\boldsymbol{X}_{(l)}^{k}{}^T \boldsymbol{y}^k}{N_k} + \sigma_k^2 \cdot \boldsymbol{\beta}_{(l)}^{k}{}^T \left( diag \left( \frac{\widetilde{\Sigma}_{kk}^{t_j}}{\psi_{(l_j)}} \right) \right) \boldsymbol{\beta}_{(l)}^k + 2 \cdot \frac{\sigma_k^2}{\sqrt{N_k}} \cdot \boldsymbol{\beta}_{(l)}^{k}{}^T \boldsymbol{A}_{(l)}^k \boldsymbol{N}_{sqrt}^k \right) \right)$$

$$\propto \exp\left( -\frac{N_k}{2\sigma_k^2} \left( \boldsymbol{\beta}_{(l)}^{k}{}^T \boldsymbol{D}_{(l)} \boldsymbol{\beta}_{(l)}^k - 2\boldsymbol{\beta}_{(l)}^{k}{}^T \widehat{\boldsymbol{\beta}_{(l)}}^k + \sigma_k^2 \cdot \boldsymbol{\beta}_{(l)}^{k}{}^T \left( diag \left( \frac{\widetilde{\Sigma}_{kk}^{t_j}}{\psi_{(l_j)}} \right) \right) \boldsymbol{\beta}_{(l)}^k + 2 \cdot \frac{\sigma_k^2}{\sqrt{N_k}} \cdot \boldsymbol{\beta}_{(l)}^{k}{}^T \boldsymbol{A}_{(l)}^k \boldsymbol{N}_{sqrt}^k \right) \right)$$

$$\propto \exp\left( -\frac{N_k}{2\sigma_k^2} \left( \boldsymbol{\beta}_{(l)}^{k}{}^T \left( \boldsymbol{D}_{(l)} + \sigma_k^2 \cdot diag \left( \frac{\widetilde{\Sigma}_{kk}^{t_j}}{\psi_{(l_j)}} \right) \right) \boldsymbol{\beta}_{(l)}^k - 2\boldsymbol{\beta}_{(l)}^{k}{}^T \left( \widehat{\boldsymbol{\beta}_{(l)}}^k - \frac{\sigma_k^2}{\sqrt{N_k}} \boldsymbol{A}_{(l)}^k \boldsymbol{N}_{sqrt}^k \right) \right) \right).$$

Therefore, we have

$$\begin{aligned} \boldsymbol{\beta}_{(l)}^k \mid \boldsymbol{D}_{(l)}^k, \widehat{\boldsymbol{\beta}_{(l)}}^k, \sigma_k^2 &\sim MVN(\boldsymbol{\mu}_{(l)}^k, \boldsymbol{\Sigma}_{(l)}^k), \\ \boldsymbol{\mu}_{(l)}^k &= \frac{N_k}{\sigma_k^2} \cdot \boldsymbol{\Sigma}_{(l)}^k \cdot \left( \widehat{\boldsymbol{\beta}_{(l)}}^k - \frac{\sigma_k^2}{\sqrt{N_k}} \cdot \boldsymbol{A}_{(l)}^k \cdot \boldsymbol{N}_{sqrt}^k \right), \\ \boldsymbol{\Sigma}_{(l)}^k &= \frac{\sigma_k^2}{N_k} \left( \boldsymbol{D}_{(l)} + \sigma_k^2 \cdot diag \left( \frac{\widetilde{\Sigma}_{kk}^{t_j}}{\psi_{(l_j)}} \right) \right)^{-1}. \end{aligned}$$

□

2. Update $\sigma_k^2 | \boldsymbol{\Psi}, \boldsymbol{\beta}^k, \widehat{\boldsymbol{\beta}}^k, \boldsymbol{D}^k$ :
   For the current updating population $k$, we update the variance by the following:

$$\sigma_k^2 | \boldsymbol{\Psi}, \boldsymbol{\beta}^k, \widehat{\boldsymbol{\beta}}^k, \boldsymbol{D}^k \sim iG\left( \frac{N_k + S_k}{2}, \frac{N_k}{2} \left\{ 1 - 2\sum_{l=1}^{L} \boldsymbol{\beta}_{(l)}^{k}{}^T \widehat{\boldsymbol{\beta}}_{(l)}^k + \sum_{l=1}^{L} \boldsymbol{\beta}_{(l)}^{k}{}^T \left( \boldsymbol{D}_{(l)} + \boldsymbol{\Psi}_{(l)}^{-1} \right) \boldsymbol{\beta}_{(l)}^k \right\} \right). \quad (2)$$

Here $iG(\alpha, \beta)$ is the inverse-gamma distribution with the probability density function

$$f(x; \alpha, \beta) = \frac{\beta^\alpha}{\Gamma(\alpha)} \left(\frac{1}{x}\right)^{\alpha+1} \exp\left(-\frac{\beta}{x}\right).$$

*Proof.* Based on the joint model we propose, we can estimate the diagonal elements of the covariance matrix by assuming

$$\beta_j^k \sim N\left(0, \Psi_j \frac{\sigma_k^2}{N_k}\right), \quad f(\sigma_k^2) \sim \frac{1}{\sigma_k^2}.$$

for each population $k$. We note that when we update $\sigma_k^2$, we only use SNPs available in population $k$. Then we have

$$f(\boldsymbol{y}^k|\sigma_k^2) \propto \left(\sigma_k^2\right)^{-\frac{N_k}{2}} \cdot \exp\left\{-\frac{1}{2} \cdot \frac{(\boldsymbol{y}^k - \boldsymbol{X}^k\boldsymbol{\beta}^k)^T(\boldsymbol{y}^k - \boldsymbol{X}^k\boldsymbol{\beta}^k)}{\sigma_k^2}\right\}$$

$$\propto \left(\frac{1}{\sigma_k^2}\right)^{\frac{N_k}{2}} \cdot \exp\left\{-\frac{N_k}{2} \cdot \frac{1 - 2\sum_{l=1}^{L} \boldsymbol{\beta}_{(l)}^{k\ T} \widehat{\boldsymbol{\beta}_{(l)}}^{k} + \sum_{l=1}^{L} \boldsymbol{\beta}_{(l)}^{k\ T} \boldsymbol{D}_{(l)} \boldsymbol{\beta}_{(l)}^{k}}{\sigma_k^2}\right\}$$

$$f(\boldsymbol{\beta}^k|\sigma_k^2) \propto \left(\sigma_k^2\right)^{-\frac{S_k}{2}} \exp\left\{-\frac{N_k}{2} \frac{\sum_{l=1}^{L} \boldsymbol{\beta}_{(l)}^{k\ T} \boldsymbol{\Psi}_{(l)}^{-1} \boldsymbol{\beta}_{(l)}^{k}}{\sigma_k^2}\right\}$$

$$f(\sigma_k^2|\boldsymbol{\beta}^k, \widehat{\boldsymbol{\beta}}^k, \boldsymbol{D}^k) \propto f(\sigma_k^2) \cdot f(\boldsymbol{y}^k|\sigma_k^2) \cdot f(\boldsymbol{\beta}^k|\sigma_k^2)$$

$$= \left(\frac{1}{\sigma_k^2}\right)^{1+\frac{N_k+S_k}{2}} \cdot \exp\left\{-\frac{N_k}{2} \cdot \frac{1 - 2\sum_{l=1}^{L} \boldsymbol{\beta}_{(l)}^{k\ T} \widehat{\boldsymbol{\beta}}_{(l)}^{k} + \sum_{l=1}^{L} \boldsymbol{\beta}_{(l)}^{k\ T} \left(\boldsymbol{D}_{(l)} + \boldsymbol{\Psi}_{(l)}^{-1}\right) \boldsymbol{\beta}_{(l)}^{k}}{\sigma_k^2}\right\}$$

Therefore, we have

$$\sigma_k^2|\boldsymbol{\Psi}, \boldsymbol{\beta}^k, \widehat{\boldsymbol{\beta}}^k, \boldsymbol{D}^k \sim iG\left(\frac{N_k + S_k}{2}, \frac{N_k}{2}\left\{1 - 2\sum_{l=1}^{L} \boldsymbol{\beta}_{(l)}^{k\ T} \widehat{\boldsymbol{\beta}}_{(l)}^{k} + \sum_{l=1}^{L} \boldsymbol{\beta}_{(l)}^{k\ T} \left(\boldsymbol{D}_{(l)} + \boldsymbol{\Psi}_{(l)}^{-1}\right) \boldsymbol{\beta}_{(l)}^{k}\right\}\right).$$

□

3. Update each pair of correlation $\rho_{k_1k_2}$ from the upper triangle under different constraint. Based on the joint model we propose, we can estimate the covariance for two populations $k_1$ and $k_2$ by assuming:

$$\begin{pmatrix} \beta_j^{k_1} \\ \beta_j^{k_2} \end{pmatrix} \sim N\left(0, \Psi_j \boldsymbol{M} \begin{pmatrix} \sqrt{\sigma_{k_1}^2} & 0 \\ 0 & \sqrt{\sigma_{k_2}^2} \end{pmatrix} \begin{pmatrix} 1 & \rho_{k_1k_2} \\ \rho_{k_1k_2} & 1 \end{pmatrix} \begin{pmatrix} \sqrt{\sigma_{k_1}^2} & 0 \\ 0 & \sqrt{\sigma_{k_2}^2} \end{pmatrix} \boldsymbol{M}\right).$$

We note that when we update $\rho_{k_1k_2}$, we only use SNPs shared by populations $k_1$ and $k_2$. And if we denote the posterior distribution of $\rho_{k_1k_2}$ as $h_r(\rho_{k_1k_2})$, we have

$$h_r(\rho_{k_1k_2}) \equiv f(\boldsymbol{\beta}|\rho_{k_1k_2})$$
$$\propto \left(\sigma_{k_1}^2\sigma_{k_2}^2(1-\rho_{k_1k_2}^2)\right)^{-\frac{S_{k_1k_2}}{2}} \exp\left\{-\frac{1}{2}\cdot\frac{N_{k_1}N_{k2}}{\sigma_{k_1}^2\sigma_{k_2}^2(1-\rho_{k_1k_2}^2)}\cdot\sum_{j=1}^{S_{k_1k_2}}\frac{1}{\Psi_j}\left(\frac{\sigma_{k_2}^2}{N_{k_2}}\beta_{k_1}^2 - 2\frac{\sqrt{\sigma_{k_1}^2\sigma_{k_2}^2}\cdot\rho_{k_1k_2}}{\sqrt{N_{k_1}N_{k_2}}}\beta_j^{k_1}\beta_j^{k_2} + \frac{\sigma_{k_1}^2}{N_{k_1}}\beta_{k_2}^2\right)\right\}.$$

Since there is no closed-form distribution to update the correlation $\rho_{k_1k_2}$, we use the following MH algorithm:

**Algorithm 1** MH Algorithm for JointPRS

**Ensure:** $\delta_r = 0.05$
**while** itr $\leq$ n_iter **do**
  **while** $1 \leq k_1 \leq K-1$ **do**
    **while** $k_1 < k_2 \leq K$ **do**
      $\rho_{k_1k_2}^* = Uniform(\rho_{k_1k_2} - \delta_r, \rho_{k_1k_2} + \delta_r)$
      **if** $\rho_{k_1k_2}^* \in [0, cons]$ **then**
        log_ratio $= log(h_r(\rho_{k_1k_2}^*)) - log(h_r(\rho_{k_1k_2}))$
        **if** exp(log_ratio) $\geq random.Uniform(0,1)$ **then**
          $\rho_{k_1k_2} = \rho_{k_1k_2}^*$
        **end if**
      **end if**
    **end while**
  **end while**
**end while**

*Note: Here we choose* $cons = 0$ *when* $\phi = 10^{-6}$ *which is equivalent to the PRS-CSx method to avoid convergence issue and choose* $cons = 0.99$ *for all other situations to consider the possible positive correlations between population* $k_1$ *and* $k_2$.

Then we can also update the corresponding covariance pair $\sigma_{k_1k_2} = \sqrt{\sigma_{k_1}^2\sigma_{k_2}^2}\cdot\rho_{k_1k_2}$.

4. Update $\Psi_j \mid \boldsymbol{\beta}_j, \sigma^2, \delta_j$:
For each SNP $j$, we update the corresponding shrinkage parameter by the following:

$$\Psi_j \mid \boldsymbol{\beta}_j, \boldsymbol{M\Sigma M}, \delta_j \sim giG\left(a - \frac{K}{2}, 2\delta_j, \boldsymbol{\beta}_j^T(\boldsymbol{M\Sigma M})^{-1}\boldsymbol{\beta}_j\right) \equiv giG\left(a - \frac{K}{2}, 2\delta_j, \boldsymbol{\beta}_j^T\widetilde{\boldsymbol{\Sigma}}\boldsymbol{\beta}_j\right). \quad (3)$$

Here $giG(\lambda, \rho, \chi)$ is the three-parameter generalized inverse Gaussian distribution with the probability density function

$$f(x;\lambda,\rho,\chi) = \frac{(\rho/\chi)^{\lambda/2}}{2K_\lambda\sqrt{\rho\chi}}x^{\lambda-1}e^{-(\rho x+\chi/x)/2}, \quad x>0, \quad \rho>0, \quad \chi>0,$$

where $K_\lambda$ is the modified Bessel function of the second kind.

*Proof.*

$$
\begin{aligned}
f\left(\Psi_j \mid \boldsymbol{\beta}_j, \boldsymbol{M\Sigma M}, \delta_j\right) &\propto f\left(\boldsymbol{\beta}_j \mid \boldsymbol{M\Sigma M}, \delta_j, \Psi_j\right) \cdot f\left(\Psi_j \mid \delta_j\right) \\
&\propto \left(\Psi_j\right)^{-\frac{K}{2}} \exp\left(-\frac{1}{2}\frac{\boldsymbol{\beta}_j{}^T\left(\boldsymbol{M\Sigma M}\right)^{-1}\boldsymbol{\beta}_j}{\Psi_j}\right) \cdot \left(\Psi_j\right)^{a-1} \exp\left(-\delta_j\Psi_j\right) \\
&\propto \left(\Psi_j\right)^{a-\frac{K}{2}-1} \exp\left(-\frac{1}{2}\left(2\delta_j\Psi_j + \frac{\boldsymbol{\beta}_j{}^T\left(\boldsymbol{M\Sigma M}\right)^{-1}\boldsymbol{\beta}_j}{\Psi_j}\right)\right)
\end{aligned}
$$

Therefore we have

$$
\Psi_j \mid \boldsymbol{\beta}_j, \boldsymbol{M\Sigma M}, \delta_j \sim giG\left(a-\frac{K}{2}, 2\delta_j, \boldsymbol{\beta}_j{}^T\left(\boldsymbol{M\Sigma M}\right)^{-1}\boldsymbol{\beta}_j\right) \equiv giG\left(a-\frac{K}{2}, 2\delta_j, \boldsymbol{\beta}_j{}^T\widetilde{\boldsymbol{\Sigma}}\boldsymbol{\beta}_j\right).
$$

□

5. Update $\delta_j \mid \Psi_j$:
   For each SNP $j$, we update the distribution parameter by the following:

$$
\delta_j \mid \Psi_j \sim G(a+b, \Psi_j+\phi). \tag{4}
$$

Here $G(c,d)$ is the Gamma distribution with the probability density function

$$
f(x;c,d) = \frac{(d)^c}{\Gamma(c)}(x)^{c-1}e^{-d\cdot x}.
$$

*Proof.*

$$
\begin{aligned}
f(\delta_j \mid \Psi_j) &\propto f(\Psi_j \mid \delta_j) f(\delta_j) \\
&\propto \frac{(\delta_j)^a}{\Gamma(a)}(\Psi_j)^{a-1}e^{-\delta_j\Psi_j} \cdot \frac{\phi^b}{\Gamma(b)}(\delta_j)^{b-1}e^{-\phi\delta_j} \\
&\propto (\delta_j)^{a+b-1}e^{-(\Psi_j+\phi)\delta_j}.
\end{aligned}
$$

Therefore, we have

$$
\delta_j \mid \Psi_j \sim G(a+b, \Psi_j+\phi).
$$

□

### Individual-level Genetic Dataset Preparation

#### UK Biobank Dataset (UKBB)

For the UKBB dataset [2], we classified individuals into five super-populations from the 1000 Genomes Project using the procedure outlined in SDPRX [3]: European (EUR), East Asian (EAS), African (AFR), South Asian (SAS), and Admixed American (AMR). The population counts are as follows: 311,601 EUR, 2,091 EAS, 6,829 AFR, 7,857 SAS, and 636 AMR.

We obtained 22 quantitative phenotypes from the UKBB subjects using their corresponding data fields, as detailed in Table S8. For systolic and diastolic blood pressures (SBP and DBP), we integrated both manual and automated readings, adhering to the recommendations of the relevant GWAS study [4]. In the analysis of LDL-cholesterol (LDL), Total cholesterol (TC), SBP, and DBP, we excluded individuals without medication information (data fields 6177 and 6153) and adjusted the remaining data according to guidelines from pertinent GWAS literature [4, 5].

For four binary traits, we determined cases and controls based on ICD-9, ICD-10, operation codes, and self-reported disease codes. For breast cancer, we considered only female participants. For all binary traits, the effective sample size was calculated using the formula $\frac{4*N_{\text{case}}*N_{\text{control}}}{(N_{\text{case}}+N_{\text{control}})}$, where $N_{\text{case}}$ and $N_{\text{control}}$ represent the number of cases and controls, respectively.

#### All of Us Dataset (AoU)

For the AoU dataset [6], individuals were categorized into six super-populations based on reference samples from the Human Genome Diversity Project and the 1000 Genomes Project [7]: European (EUR), East Asian (EAS), African/African American (AFR), South Asian (SAS), Admixed American/Latino (AMR), and Middle Eastern (MID). The subject counts for each population are as follows: 133,581 EUR, 5,706 EAS, 56,913 AFR, 45,035 AMR, 942 MID, and 3,217 SAS. This study focuses on predicting nine quantitative phenotypes within the AFR and AMR populations of the AoU dataset.

We extracted nine quantitative phenotypes from the AoU subjects using their corresponding concept IDs, as detailed in Table S9. Notably, Height and BMI are measured only once per individual, thus the observed values will be used directly. For the remaining seven traits, each individual has multiple measurements. To ensure robustness, we will use the median value after removing outliers.

### Supplementary Figures and Tables

**Figure S1: Pipeline for Existing Methods.** Existing methods can be divided into auto methods and tune methods, each category has its own pipeline depending on the availability of a tuning dataset.

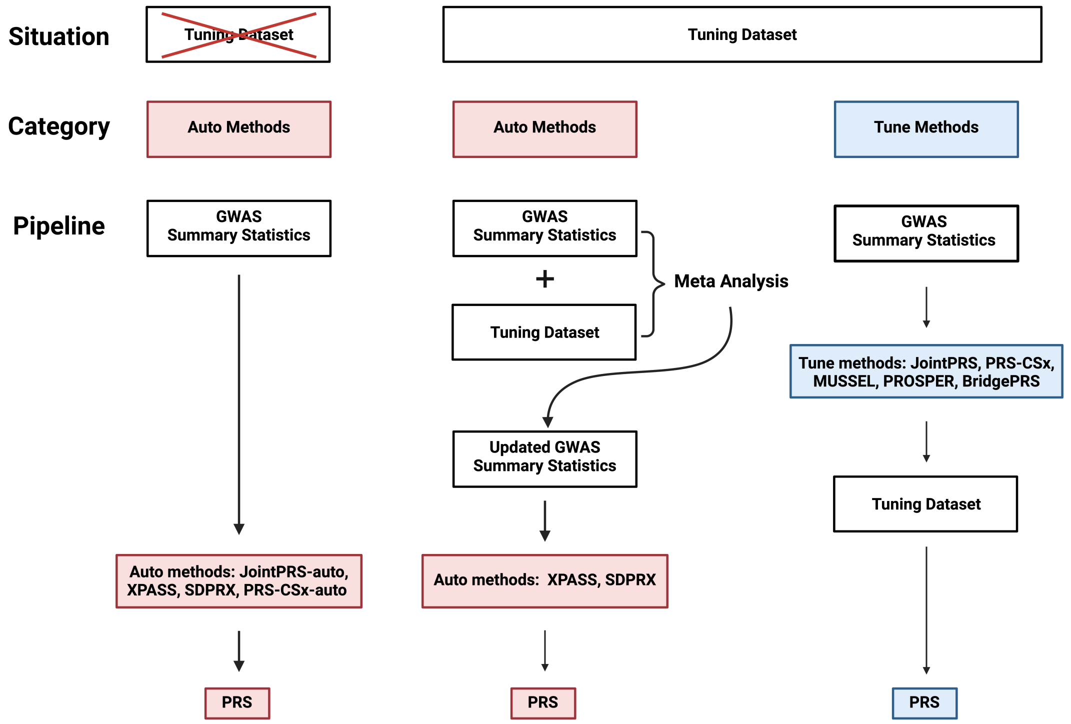

**Figure S2: Simulation Results with Large Training Datasets Sample Sizes, Small Causal SNP Proportions, and Varying Tuning Datasets Sample Sizes for Multi-population PRS Methods.** The proportion of causal SNPs was set to $p = 0.001, 5 \times 10^{-4}$. The training dataset sample size for non-European populations were set to $N_{\text{train}} = 80,000$. The tuning dataset sample size for the target population were set to $N_{\text{tune}} = 500, 2,000, 5,000, 10,000$. Simulations in each scenario were repeated five times, and the mean value were presented in the bar plots for the four non-European populations (EAS, AFR, SAS, and AMR). All methods were considered here, with the best and second-best method denoted by two stars and one star, respectively, in the corresponding bar plots. Here, SDPRX cannot provide predictions for SAS and AMR as it does not provide the corresponding LD reference panels.

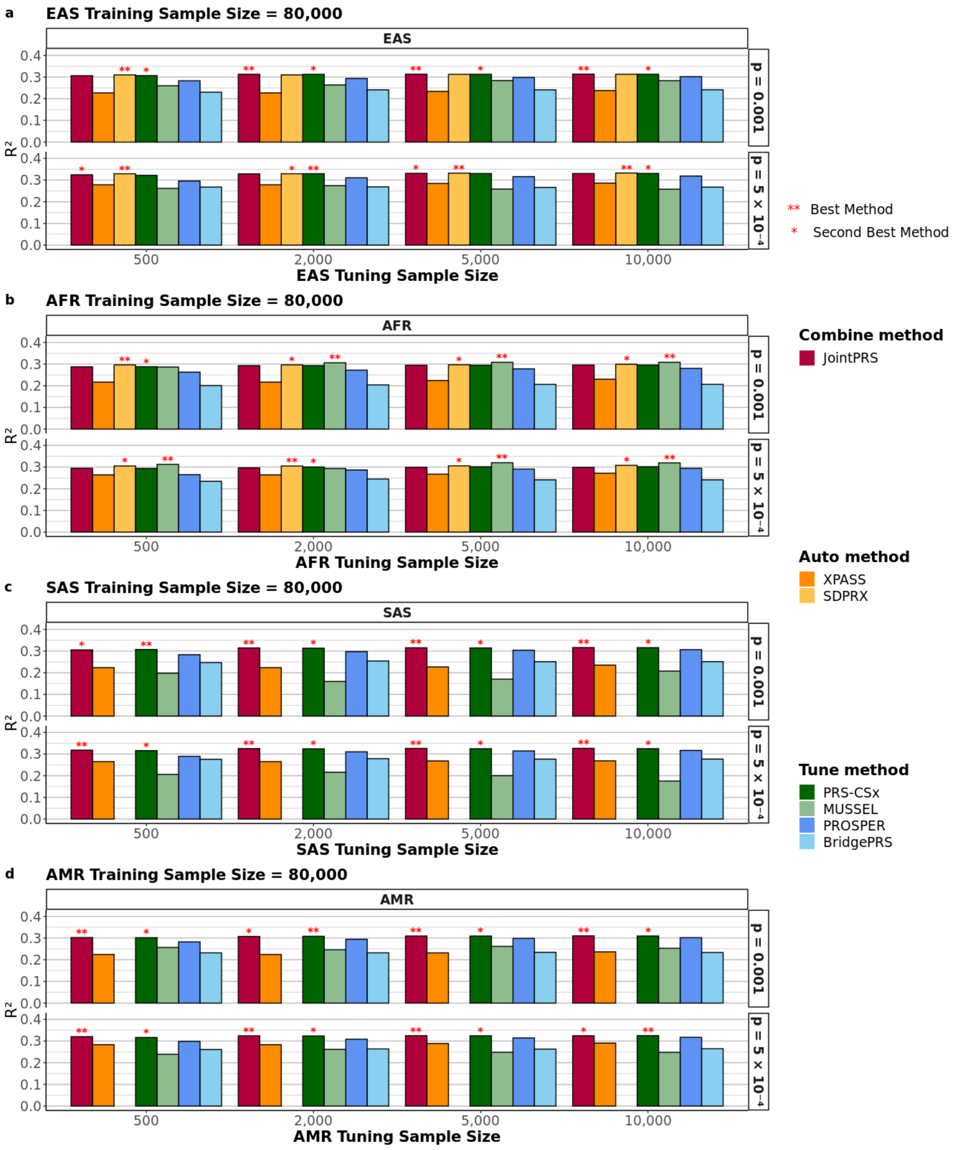

**Figure S3: Simulation Results with Small Training Datasets Sample Sizes, Large Causal SNP Proportions, and Varying Tuning Datasets Sample Sizes for Multi-population PRS Methods.** The proportion of causal SNPs was set to $p = 0.1, 0.01$. The training dataset sample size for non-European populations were set to $N_{\text{train}} = 15,000$. The tuning dataset sample size for the target population were set to $N_{\text{tune}} = 500, 2,000, 5,000, 10,000$. Simulations in each scenario were repeated five times, and the mean value were presented in the bar plots for the four non-European populations (EAS, AFR, SAS, and AMR). All methods were considered here, with the best and second-best method denoted by two stars and one star, respectively, in the corresponding bar plots. Here, SDPRX cannot provide predictions for SAS and AMR as it does not provide the corresponding LD reference panels.

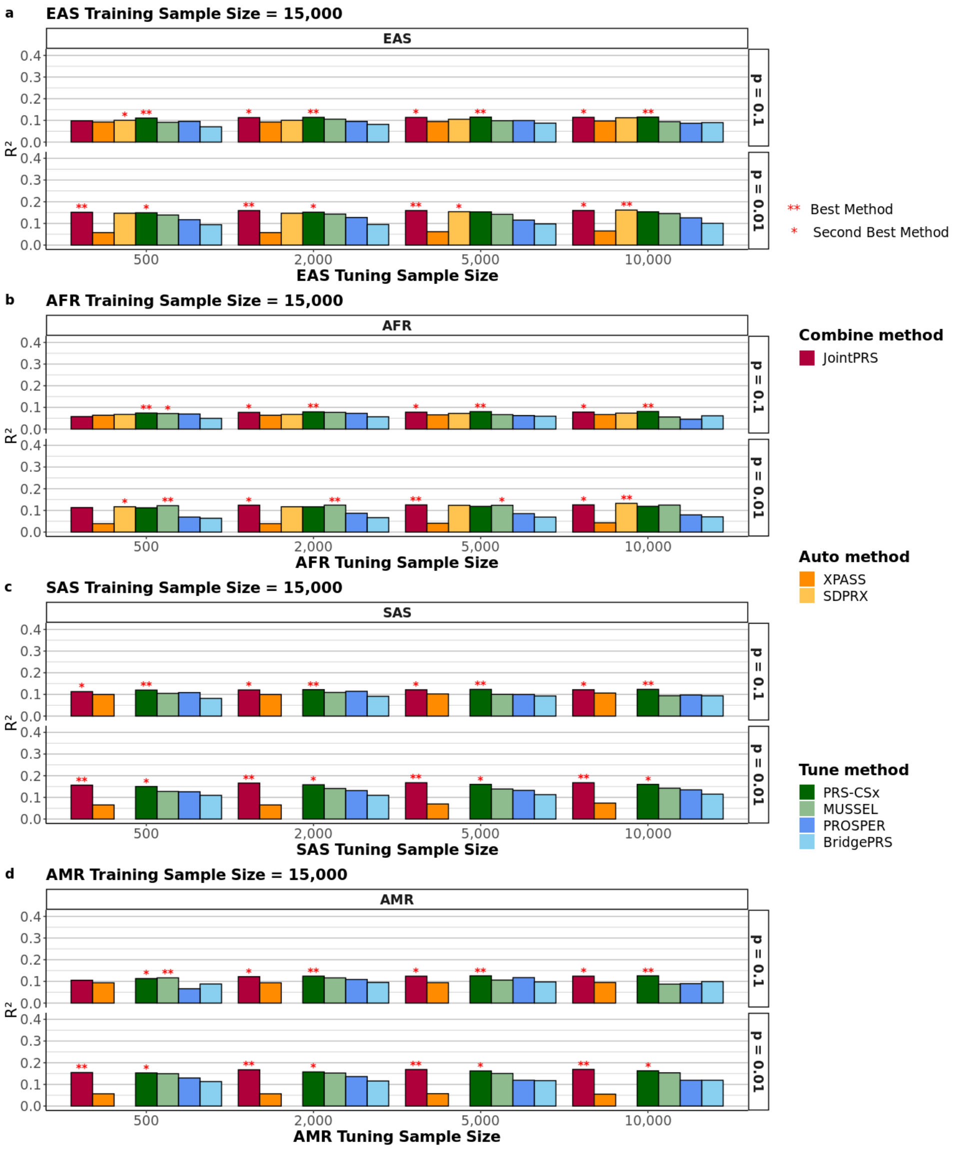

**Figure S4: Simulation Results with Small Training Datasets Sample Sizes, Small Causal SNP Proportions, and Varying Tuning Datasets Sample Sizes for Multi-population PRS Methods.** The proportion of causal SNPs was set to $p = 0.001, 5 \times 10^{-4}$. The training dataset sample size for non-European populations were set to $N_{\text{train}} = 15,000$. The tuning dataset sample size for the target population were set to $N_{\text{tune}} = 500, 2,000, 5,000, 10,000$. Simulations in each scenario were repeated five times, and the mean value were presented in the bar plots for the four non-European populations (EAS, AFR, SAS, and AMR). All methods were considered here, with the best and second-best method denoted by two stars and one star, respectively, in the corresponding bar plots. Here, SDPRX cannot provide predictions for SAS and AMR as it does not provide the corresponding LD reference panels.

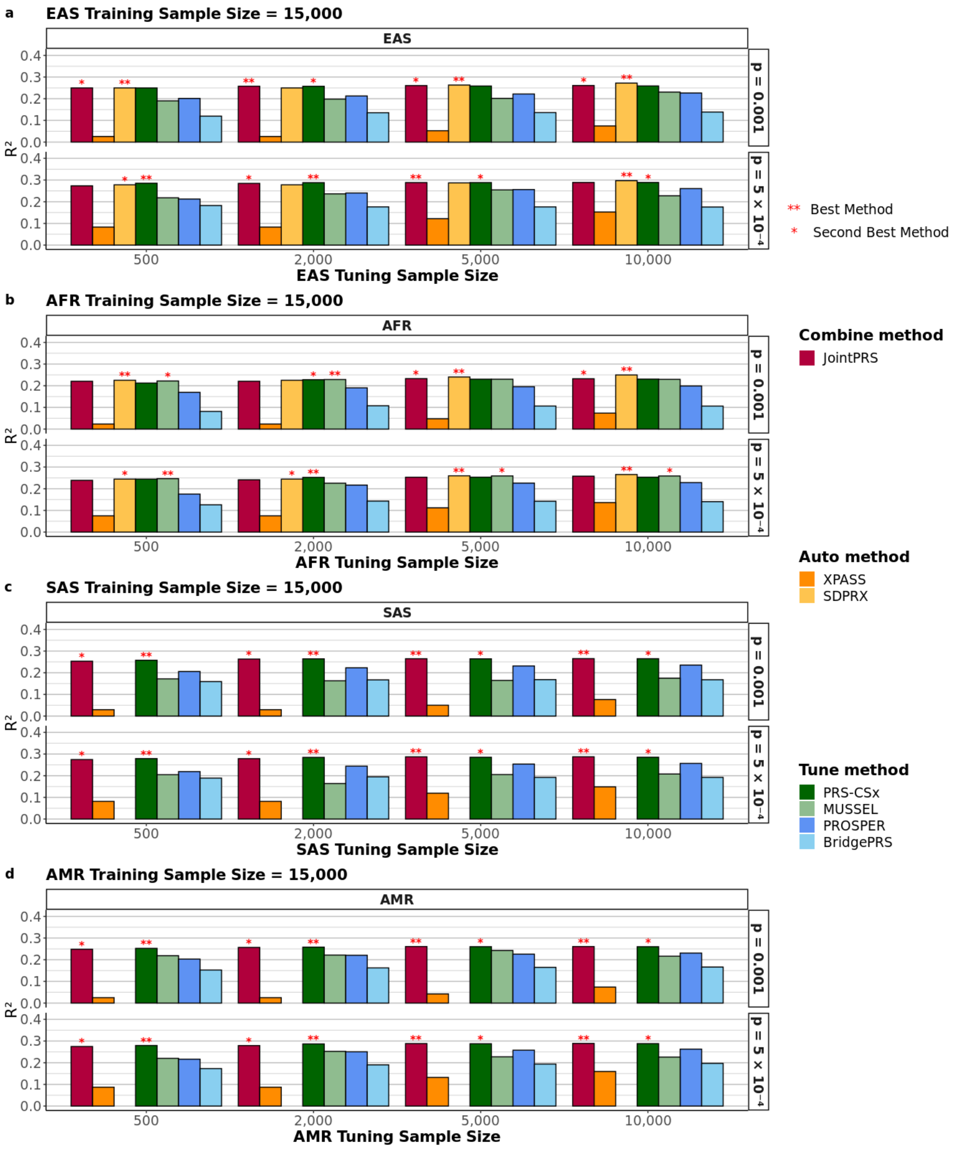

**Figure S5: EAS Population Simulation Results with Different Causal SNP Proportions and Varying Tuning Datasets Sample Sizes for Multi-population PRS Methods.** The proportion of causal SNPs was set to $p = 0.1, 0.01, 0.001, 5 \times 10^{-4}$. The training dataset sample size for non-European populations were set to $N_{\text{train}} = 80,000, 15,000$. The tuning dataset sample size for the target population were set to $N_{\text{tune}} = 500, 2,000, 5,000, 10,000$. Simulations in each scenario were repeated five times, and the mean value were presented in the bar plots for the EAS population.

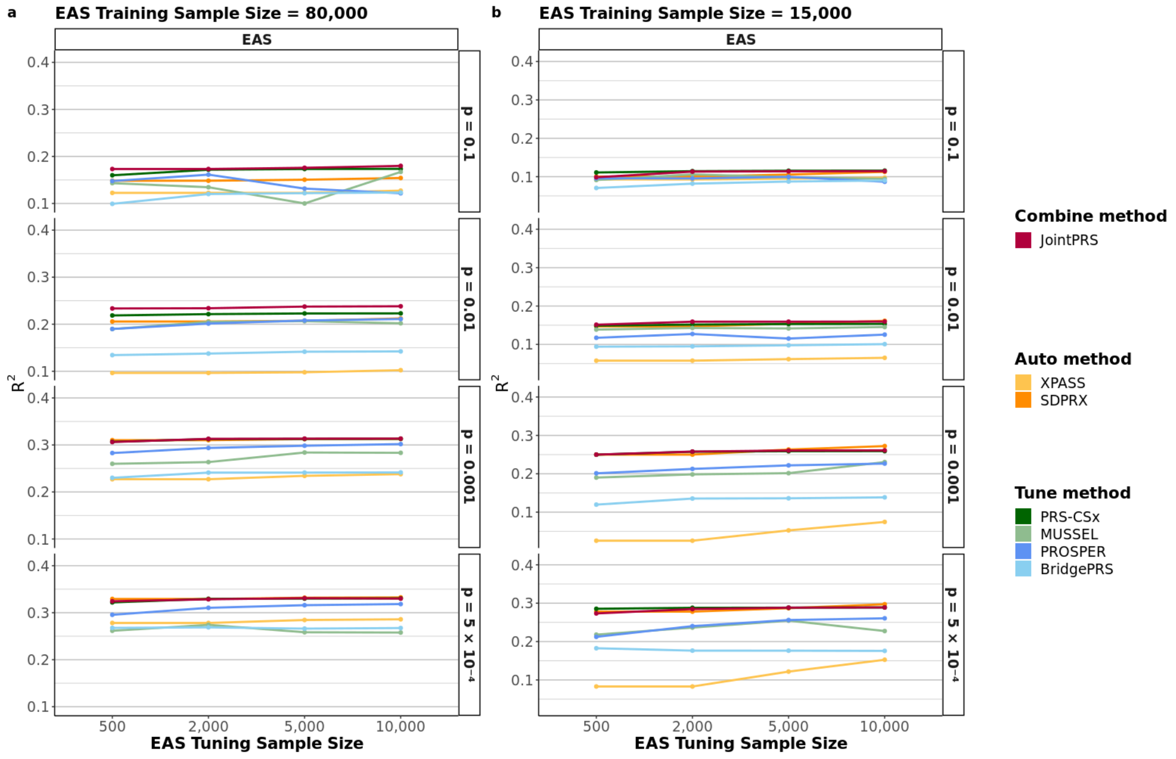

**Figure S6: AFR Population Simulation Results with Different Causal SNP Proportions and Varying Tuning Datasets Sample Sizes for Multi-population PRS Methods.** The proportion of causal SNPs was set to $p = 0.1, 0.01, 0.001, 5 \times 10^{-4}$. The training dataset sample size for non-European populations were set to $N_{\text{train}} = 80,000, 15,000$. The tuning dataset sample size for the target population were set to $N_{\text{tune}} = 500, 2,000, 5,000, 10,000$. Simulations in each scenario were repeated five times, and the mean value were presented in the bar plots for the AFR population.

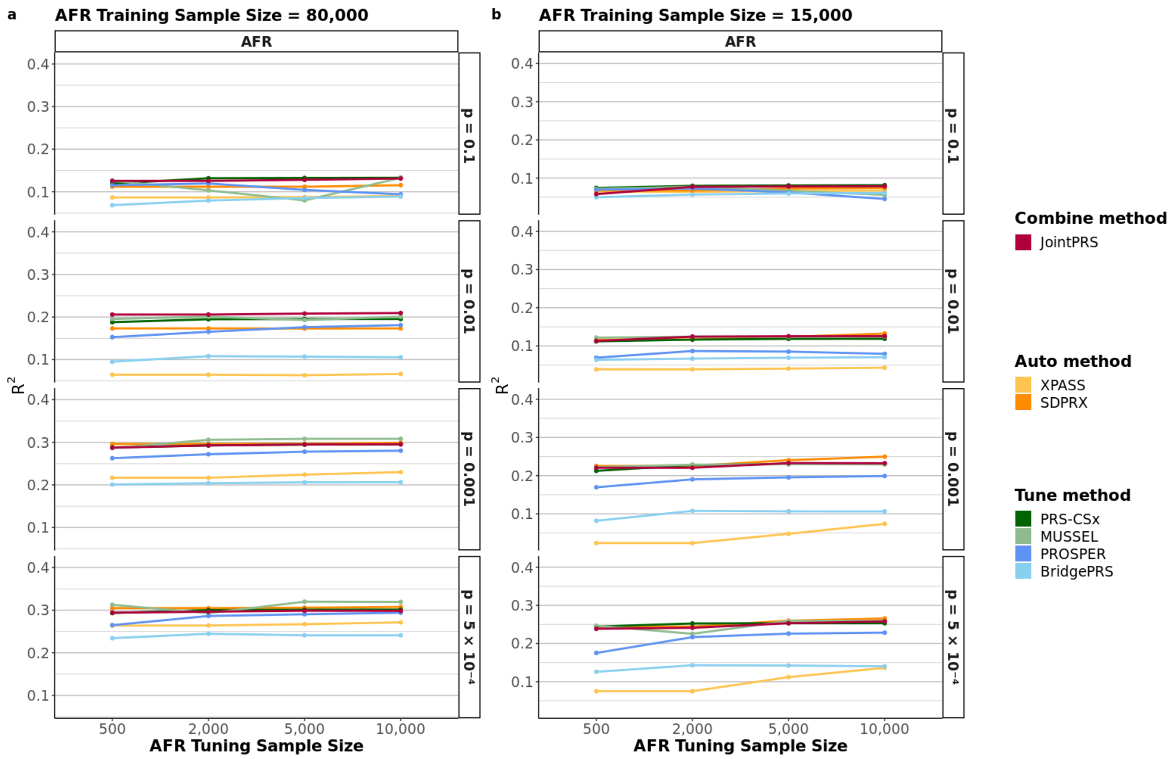

**Figure S7: SAS Population Simulation Results with Different Causal SNP Proportions and Varying Tuning Datasets Sample Sizes for Multi-population PRS Methods.** The proportion of causal SNPs was set to $p = 0.1, 0.01, 0.001, 5 \times 10^{-4}$. The training dataset sample size for non-European populations were set to $N_{\text{train}} = 80,000, 15,000$. The tuning dataset sample size for the target population were set to $N_{\text{tune}} = 500, 2,000, 5,000, 10,000$. Simulations in each scenario were repeated five times, and the mean value were presented in the bar plots for the SAS population. Here, SDPRX cannot provide predictions for SAS as it does not provide the corresponding LD reference panels.

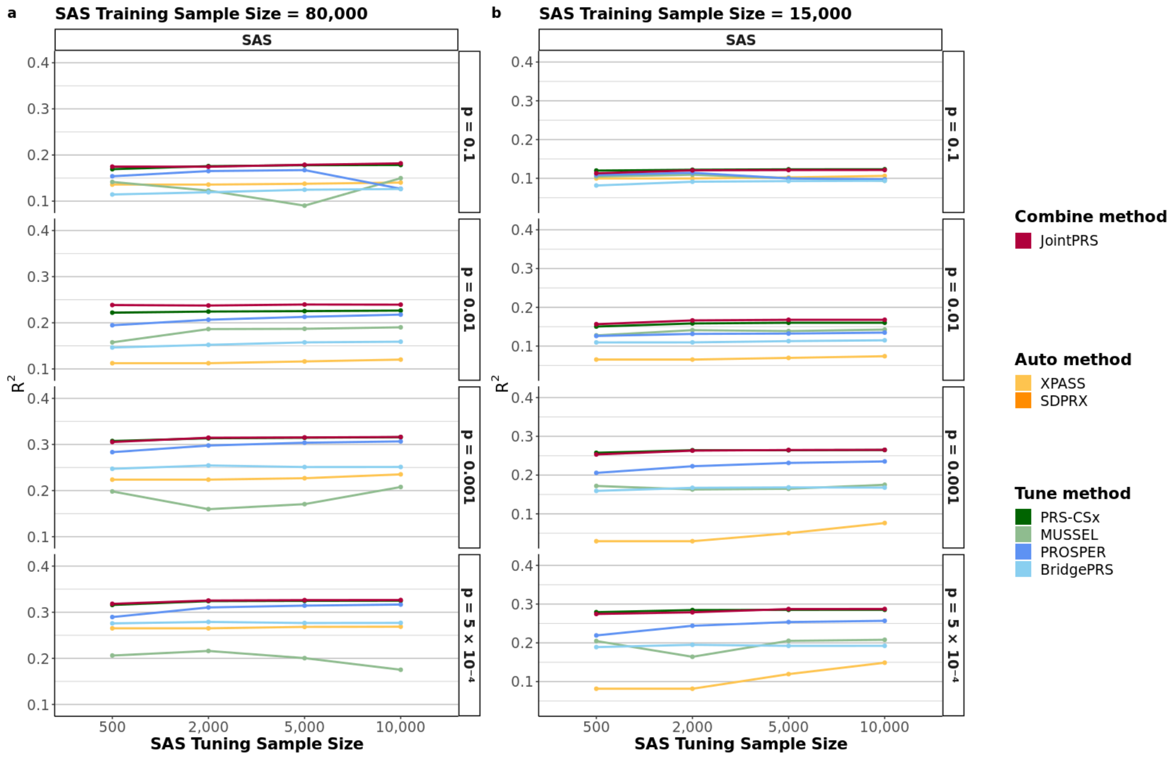

**Figure S8: AMR Population Simulation Results with Different Causal SNP Proportions and Varying Tuning Datasets Sample Sizes for Multi-population PRS Methods.** The proportion of causal SNPs was set to $p = 0.1, 0.01, 0.001, 5 \times 10^{-4}$. The training dataset sample size for non-European populations were set to $N_{\text{train}} = 80,000, 15,000$. The tuning dataset sample size for the target population were set to $N_{\text{tune}} = 500, 2,000, 5,000, 10,000$. Simulations in each scenario were repeated five times, and the mean value were presented in the bar plots for the AMR population. Here, SDPRX cannot provide predictions for AMR as it does not provide the corresponding LD reference panels.

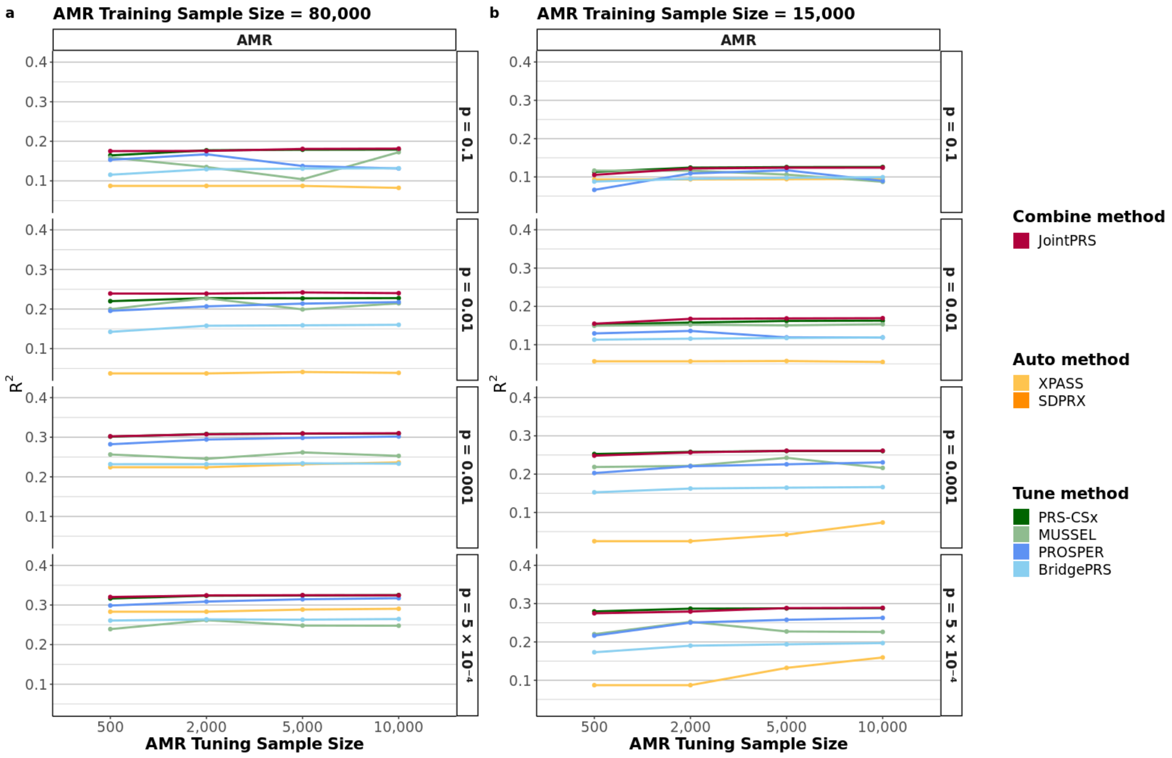

**Figure S9: Simulation Results with Varying Cross-population Genetic Correlation Settings for JointPRS-auto and PRS-CSx-auto in No Tuning Dataset Setting.** The cross-population genetic correlations were set to $\rho = 0, 0.2, 0.4, 0.6, 0.8$. The causal SNP proportion was set to $p = 0.1$. The training dataset sample size for non-European populations were set to $N_{\text{train}} = 80,000$. Simulations in each scenario were repeated five times, and the mean value were presented in the bar plots for four non-European populations (EAS, AFR, SAS, AMR).

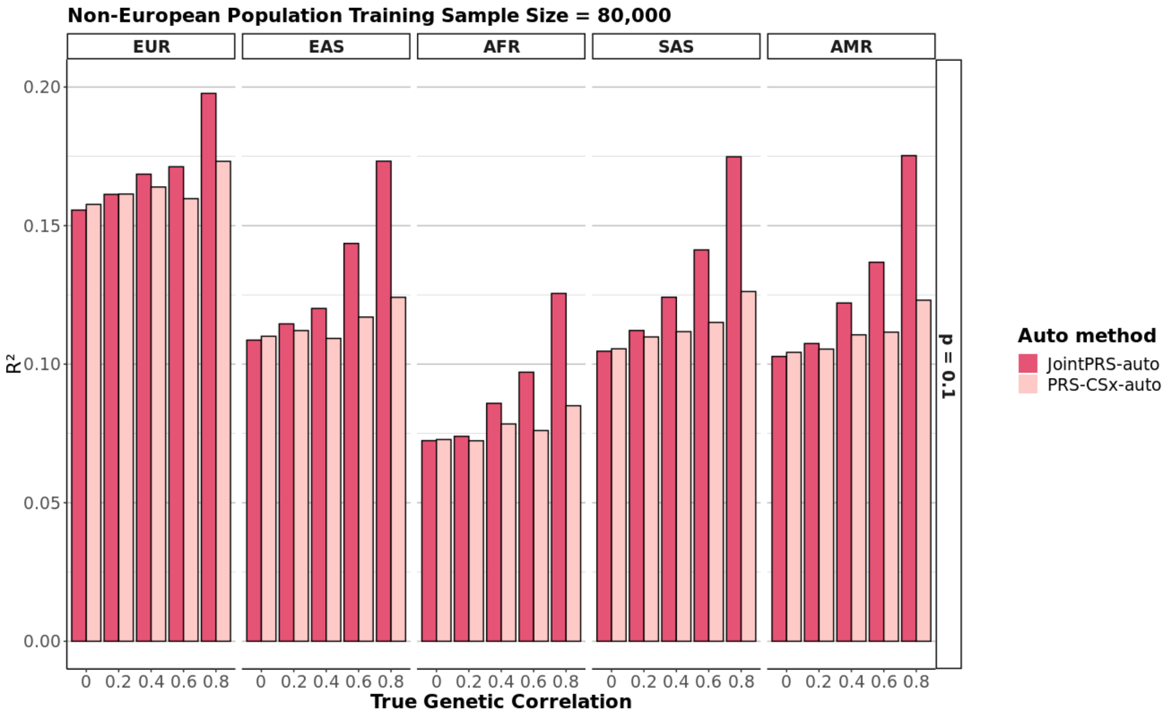

**Figure S10: Relative Prediction Accuracy of Multi-population PRS Methods in Comparison to JointPRS-auto across 26 Traits and Four Non-European Populations in UKBB When there is no Tuning Dataset.** 26 traits from four categories we predefined were considered for the data scenarios when there is no tuning dataset. The relative change of the performance for three existing auto methods over that of JointPRS measured in $R^2$ and AUC were calculated for quantitative traits and binary traits. The results were presented in the violin plots for the four non-European populations (EAS, AFR, SAS, and AMR), with the mean value across traits presented as the black crossbar for each method in each trait type. Here, SDPRX cannot provide predictions for SAS and AMR as it does not provide the corresponding LD reference panels.

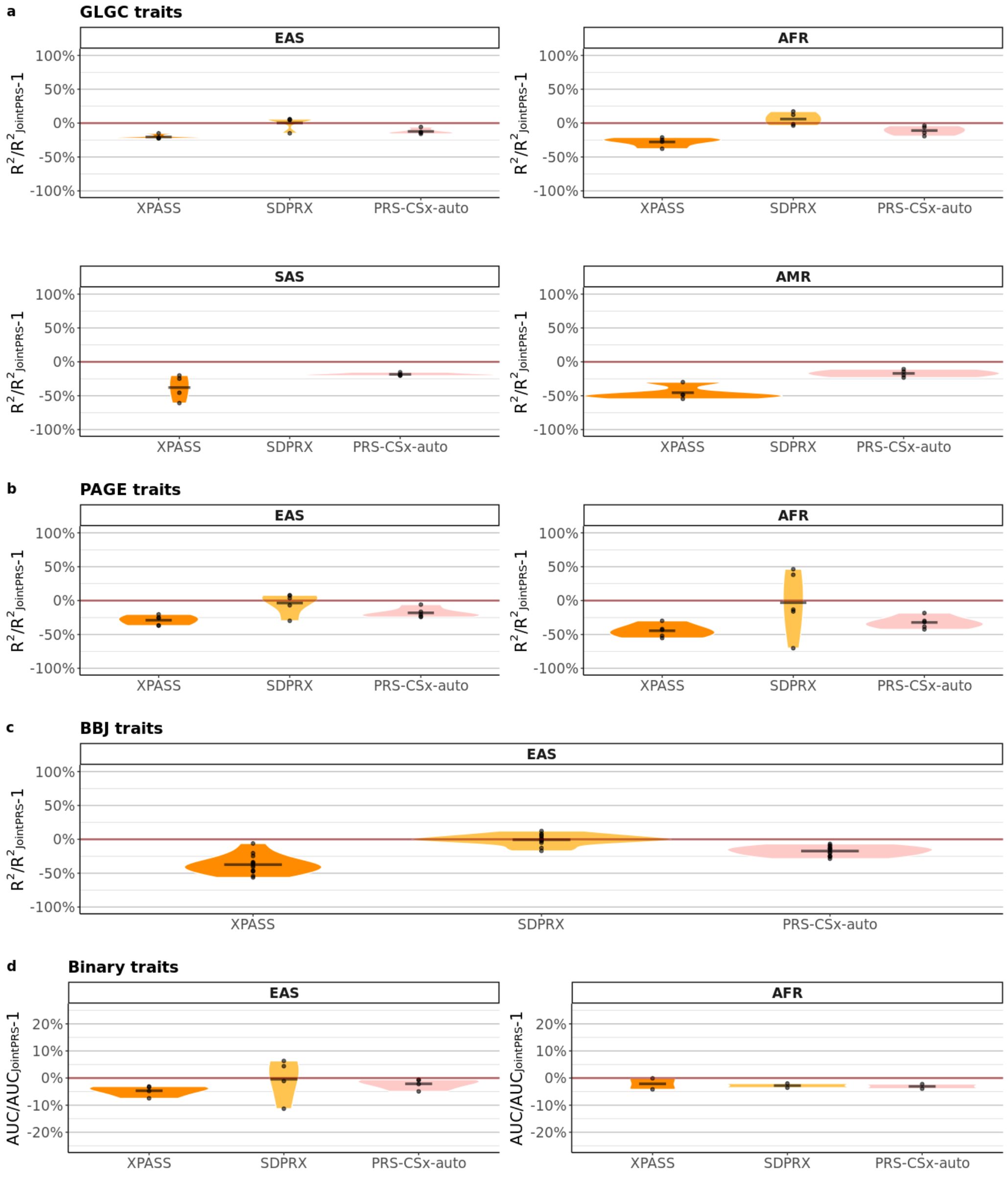

**Figure S11: Relative Improvement of JointPRS-auto over XPASS across 26 Traits and Four Non-European Populations in UKBB When there is no Tuning Dataset.** 26 traits from four populations (EAS, AFR, SAS, and AMR) were considered for the data scenarios when there is no tuning dataset. The relative improvement of the performance for JointPRS-auto over that of XPASS measured in $R^2$ and AUC were calculated for quantitative traits and binary traits. The results were presented in the bar with the pink color represent positive improvement and blue color representing negative value.

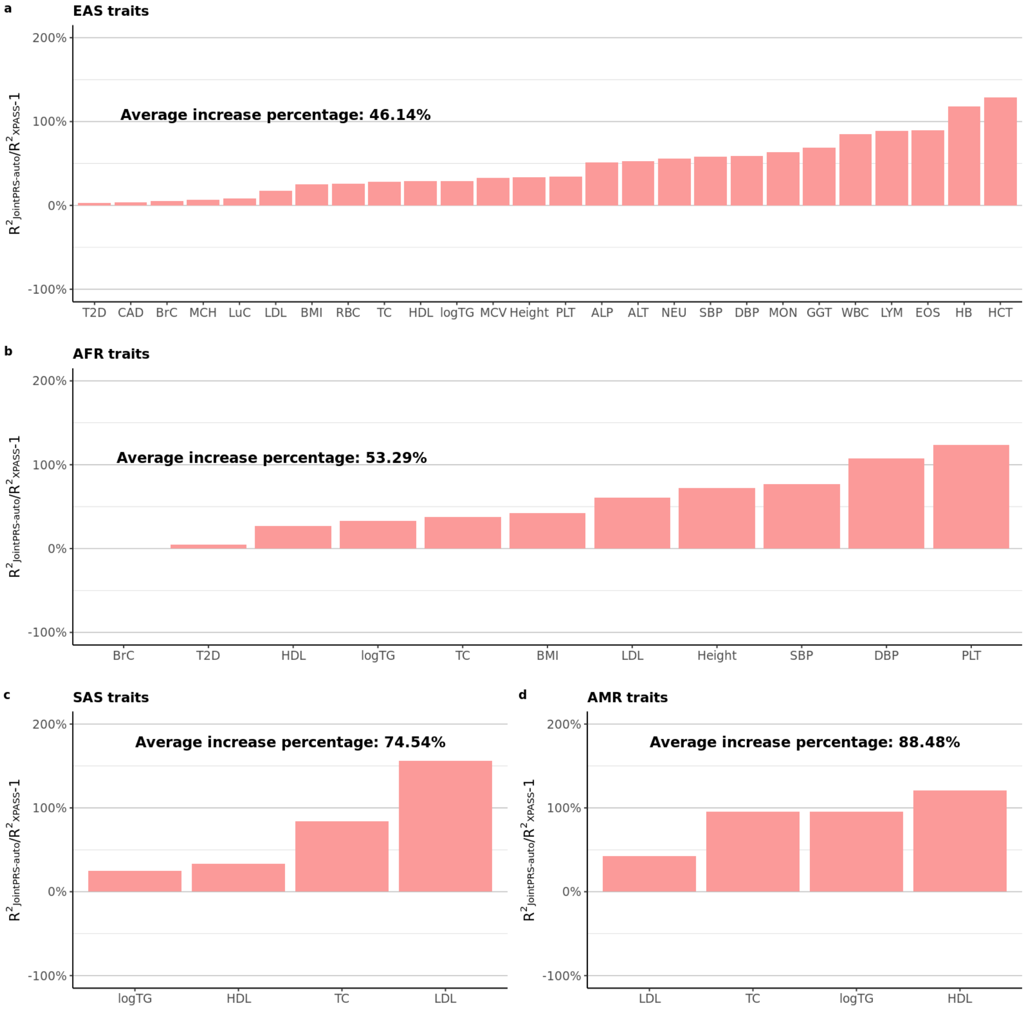

**Figure S12: Relative Improvement of JointPRS-auto over SDPRX across 26 Traits and Two Non-European Populations in UKBB When there is no Tuning Dataset.** 26 traits from two populations (AFR and AMR) were considered for the data scenarios when there is no tuning dataset. The relative improvement of the performance for JointPRS-auto over that of SDPRX measured in $R^2$ and AUC were calculated for quantitative traits and binary traits. The results were presented in the bar with the pink color represent positive improvement and blue color representing negative value. Here, SDPRX cannot provide predictions for SAS and AMR as it does not provide the corresponding LD reference panels.

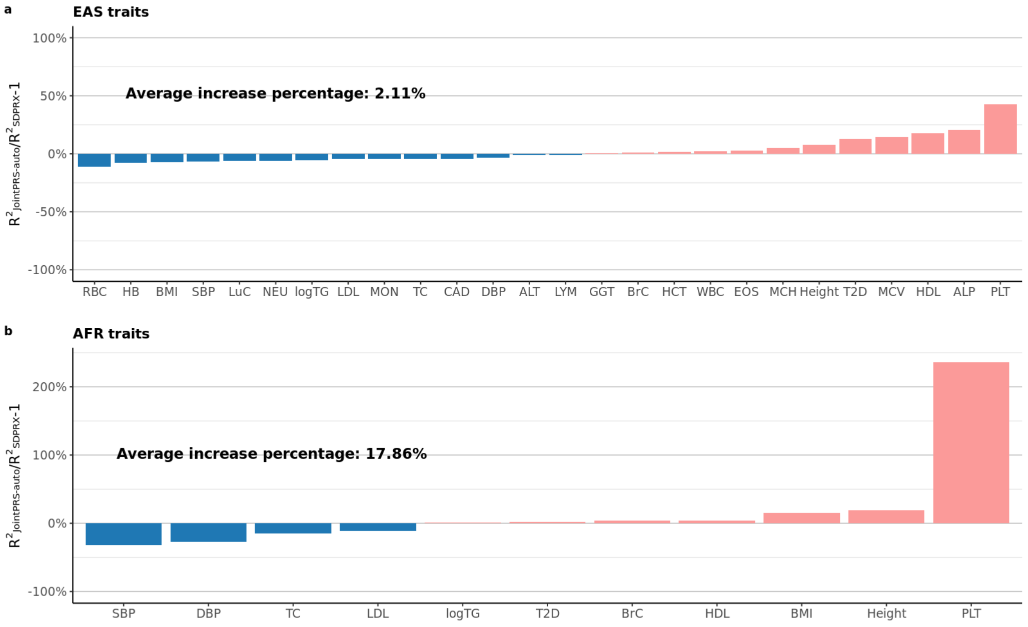

**Figure S13: Relative Improvement of JointPRS-auto over PRS-CSx-auto across 26 Traits and Four Non-European Populations in UKBB When there is no Tuning Dataset.** 26 traits from four populations (EAS, AFR, SAS, and AMR) were considered for the data scenarios when there is no tuning dataset. The relative improvement of the performance for JointPRS-auto over that of PRS-CSx-auto measured in $R^2$ and AUC were calculated for quantitative traits and binary traits. The results were presented in the bar with the pink color represent positive improvement and blue color representing negative value.

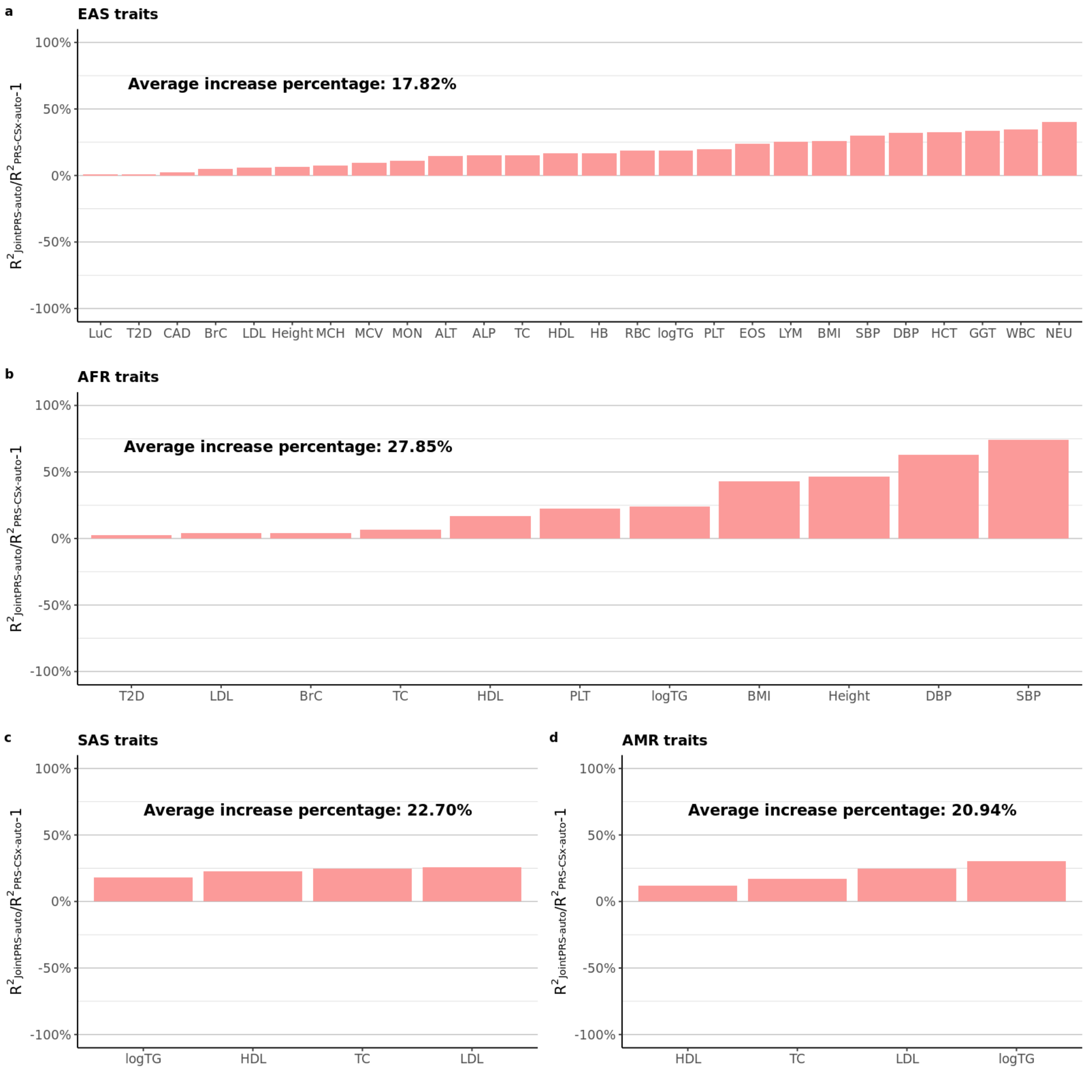

**Figure S14: Chromosome-wise Genetic Correlation Estimated by JointPRS-auto across 22 Traits between European Population and Four non-European Populations in UKBB When there is no Tuning Dataset.** Chromosome-wise genetic correlation for 22 traits between European population and four non-European populations (EAS, AFR, SAS, AMR) were considered for the data scenarios when there is no tuning dataset. The estimated correlation for 22 chromosomes using JointPRS-auto were presented in the boxplot with different color representing different trait groups.

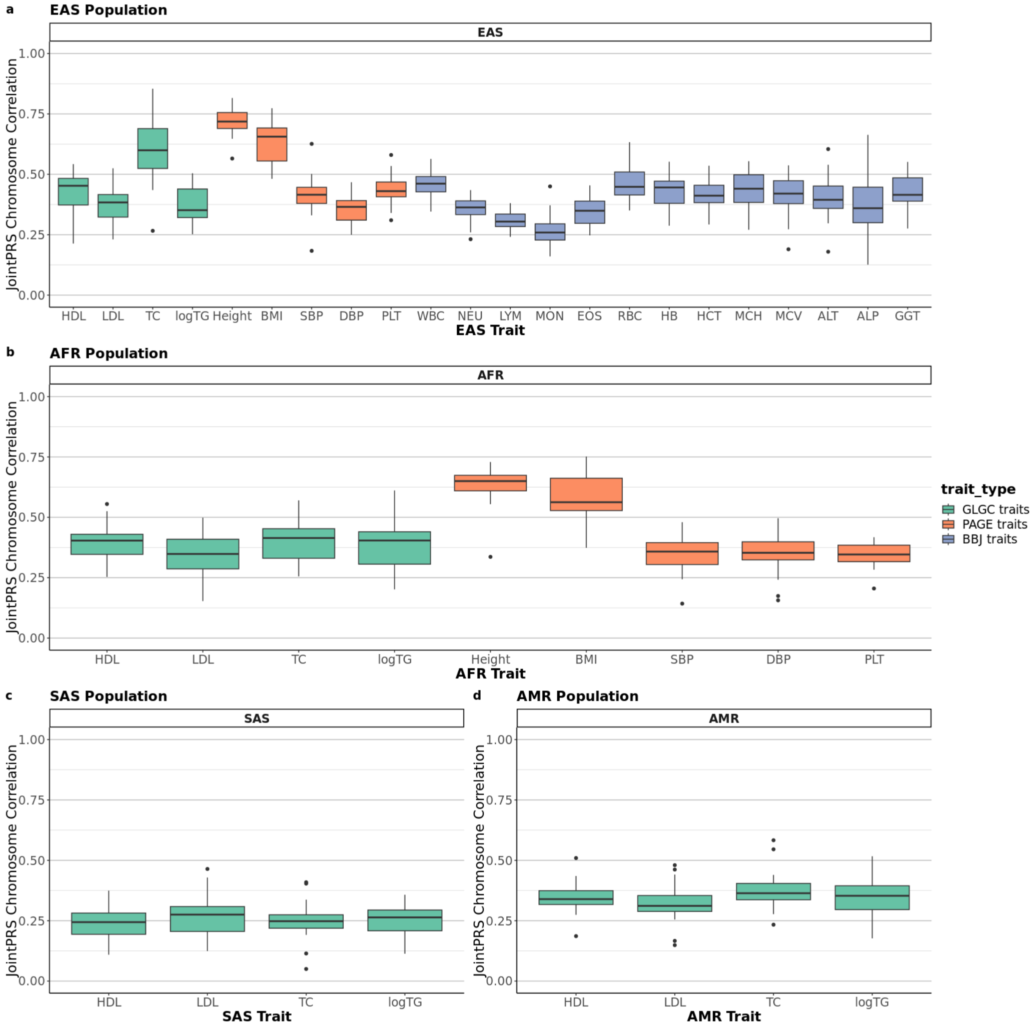

**Figure S15: Relationship between Mean of Chromosome-wise Genetic Correlation Estimated by JointPRS-auto and Percentage Improvement of JointPRS-auto over PRS-CSx-auto across 22 Traits in Four non-European Populations in UKBB When there is no Tuning Dataset.** 22 traits from four non-European populations (EAS, AFR, SAS, AMR) were considered for the data scenarios when there is no tuning dataset. The mean of estimated correlation across 22 chromosomes using JointPRS-auto and the percentages improvement of prediction accuracy measures in $R^2$ for JointPRS-auto over PRS-CSx-auto were presented in the scatter plot with different color representing different trait groups.

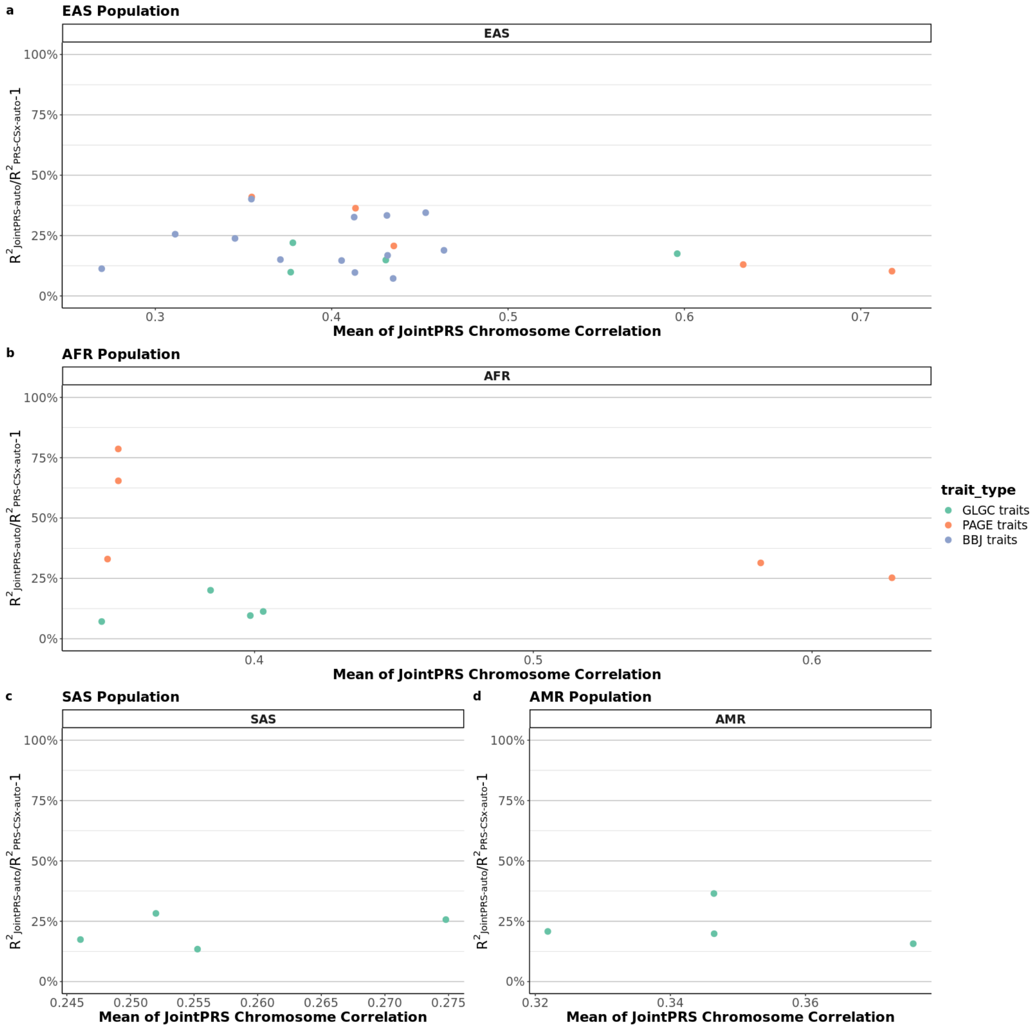

**Figure S16: Comparing JointPRS-auto, Xwing-auto, and PRS-CSx-auto based on Relative Improvement across 22 Traits in EAS Populations in UKBB When there is no Tuning Dataset.** 22 traits from EAS populations were considered for the data scenarios when there is no tuning dataset. The relative improvement of the performance for JointPRS-auto over that of PRS-CSx-auto, the relative improvement of the performance for JointPRS-auto over that of Xwing-auto, and the relative improvement of the performance for Xwing-auto over that of PRS-CSx-auto measured in $R^2$ were calculated for quantitative traits. The results were presented in the bar with the pink color representing positive improvement and blue color representing negative value.

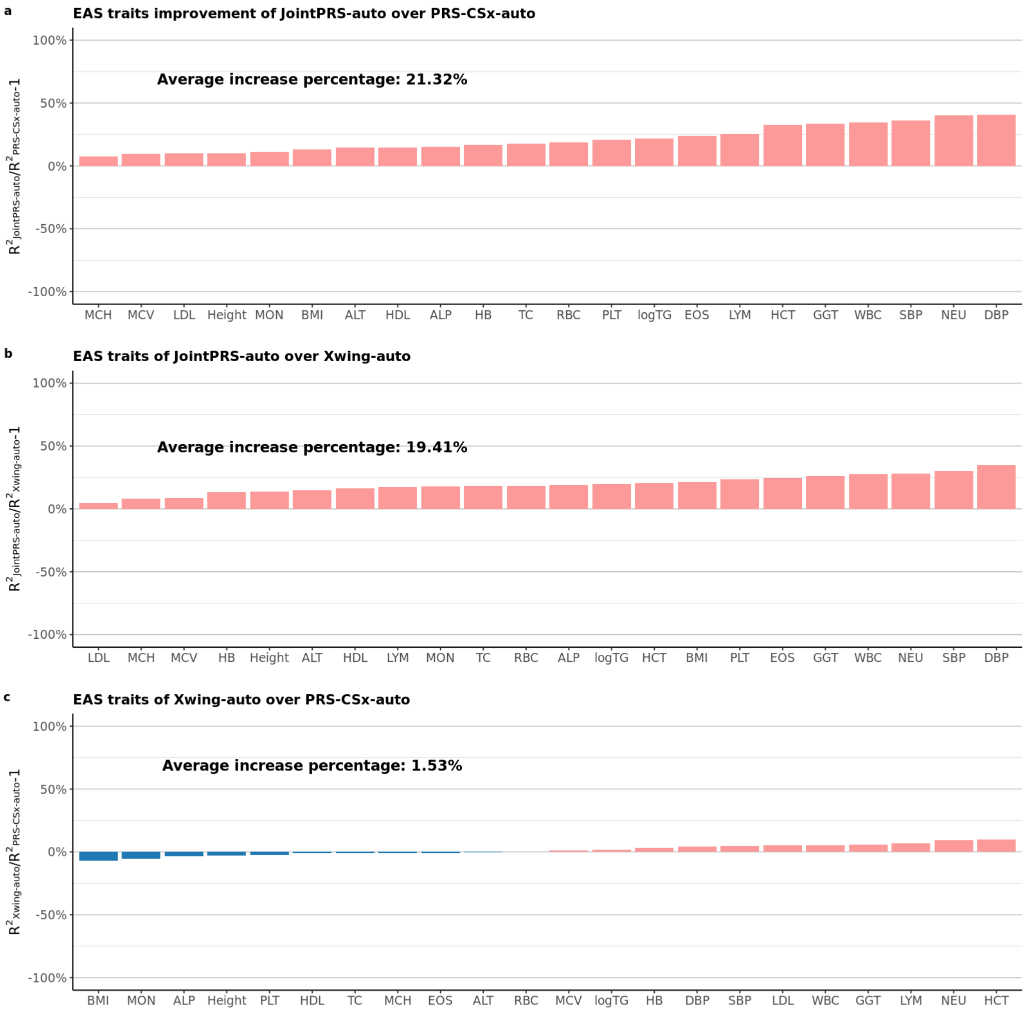

**Figure S17: Prediction Accuracy of Multi-population PRS Methods across 26 Traits and Four Non-European Populations When the Tuning and Testing Data Come From the Same Cohort (UKBB).** 26 traits from four categories we predefined were considered for the data scenarios when the tuning and testing data come from the same cohort (UKBB). The whole UKBB data was spilt into five folds, and we performed 5-fold cross-validation with each fold being treated as the testing data and the rest data being treated as the tuning data. The mean of $R^2$ and AUC across five folds were used as evaluation metric for quantitative traits and binary traits and the value were presented in the bar plots for the four non-European populations (EAS, AFR, SAS, and AMR). All seven methods were considered here, with the best and second-best method denoted by two stars and one star, respectively, in the corresponding bar plots. Here, SDPRX cannot provide predictions for SAS and AMR as it does not provide the corresponding LD reference panels.

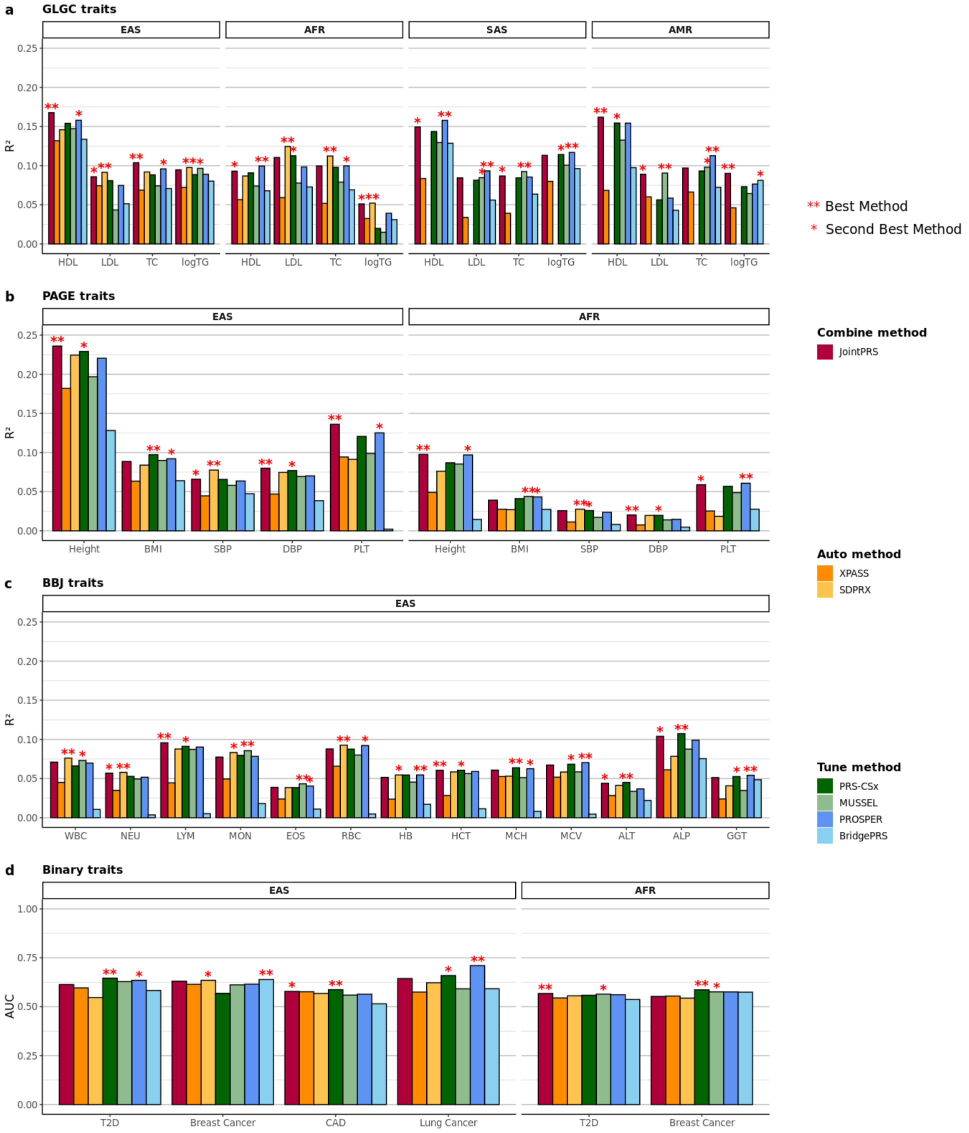

**Figure S18: Relative Improvement of JointPRS over XPASS across 26 Traits and Four Non-European Populations When the Tuning and Testing Data Come From the Same Cohort (UKBB).** 26 traits from four populations were considered for the data scenarios when the tuning and testing data come from the same cohort. The whole UKBB data was spilt into five folds, and we performed 5-fold cross-validation with each fold being treated as the testing data and the rest data being treated as the tuning data. The relative improvement of the performance for JointPRS over that of XPASS measured in mean of $R^2$ and AUC across five folds were calculated for quantitative traits and binary traits. The results were presented in the bar with the pink color represent positive improvement and blue color representing negative value.

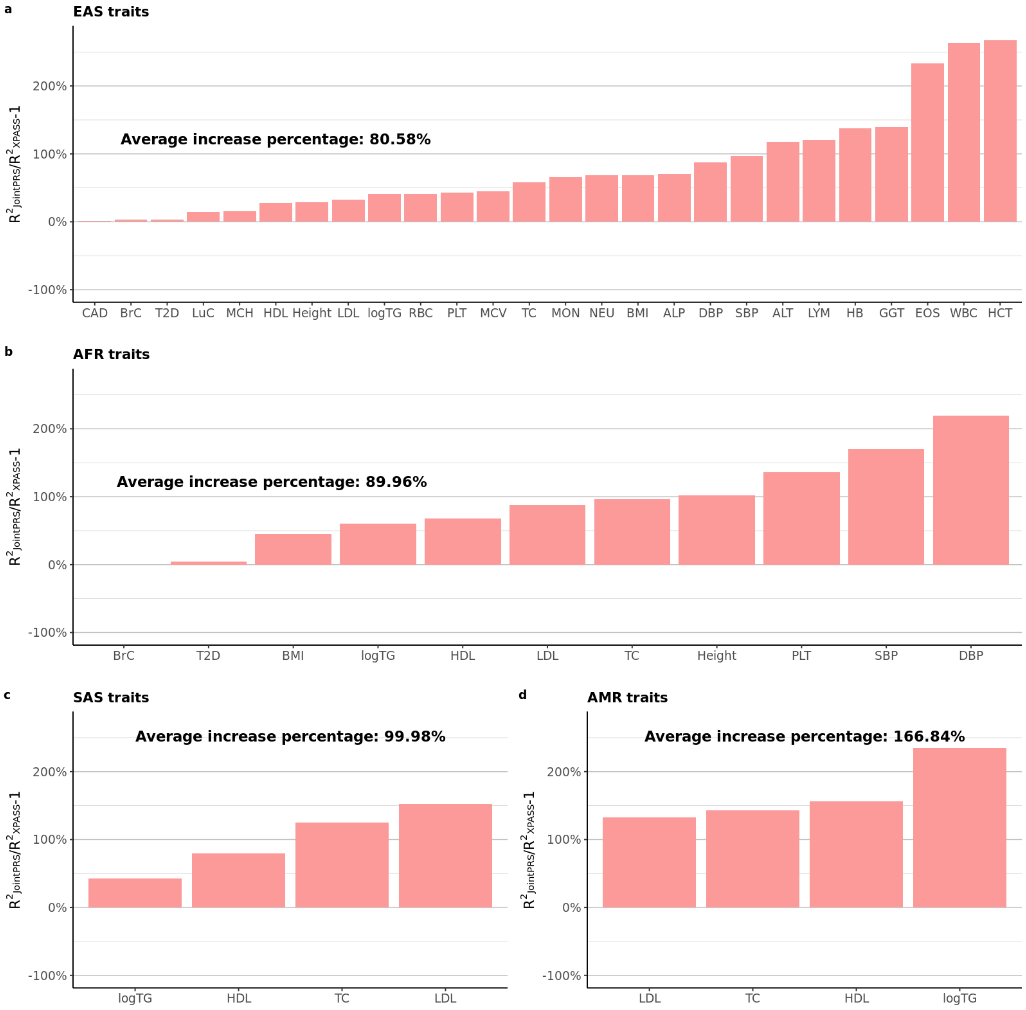

**Figure S19: Relative Improvement of JointPRS over SDPRX across 26 Traits and Two Non-European Populations When the Tuning and Testing Data Come From the Same Cohort (UKBB).** 26 traits from two populations (AFR and AMR) were considered for the data scenarios when the tuning and testing data come from the same cohort. The whole UKBB data was spilt into five folds, and we performed 5-fold cross-validation with each fold being treated as the testing data and the rest data being treated as the tuning data. The relative improvement of the performance for JointPRS over that of SDPRX measured in mean of $R^2$ and AUC across five folds were calculated for quantitative traits and binary traits. The results were presented in the bar with the pink color represent positive improvement and blue color representing negative value. Here, SDPRX cannot provide predictions for SAS and AMR as it does not provide the corresponding LD reference panels.

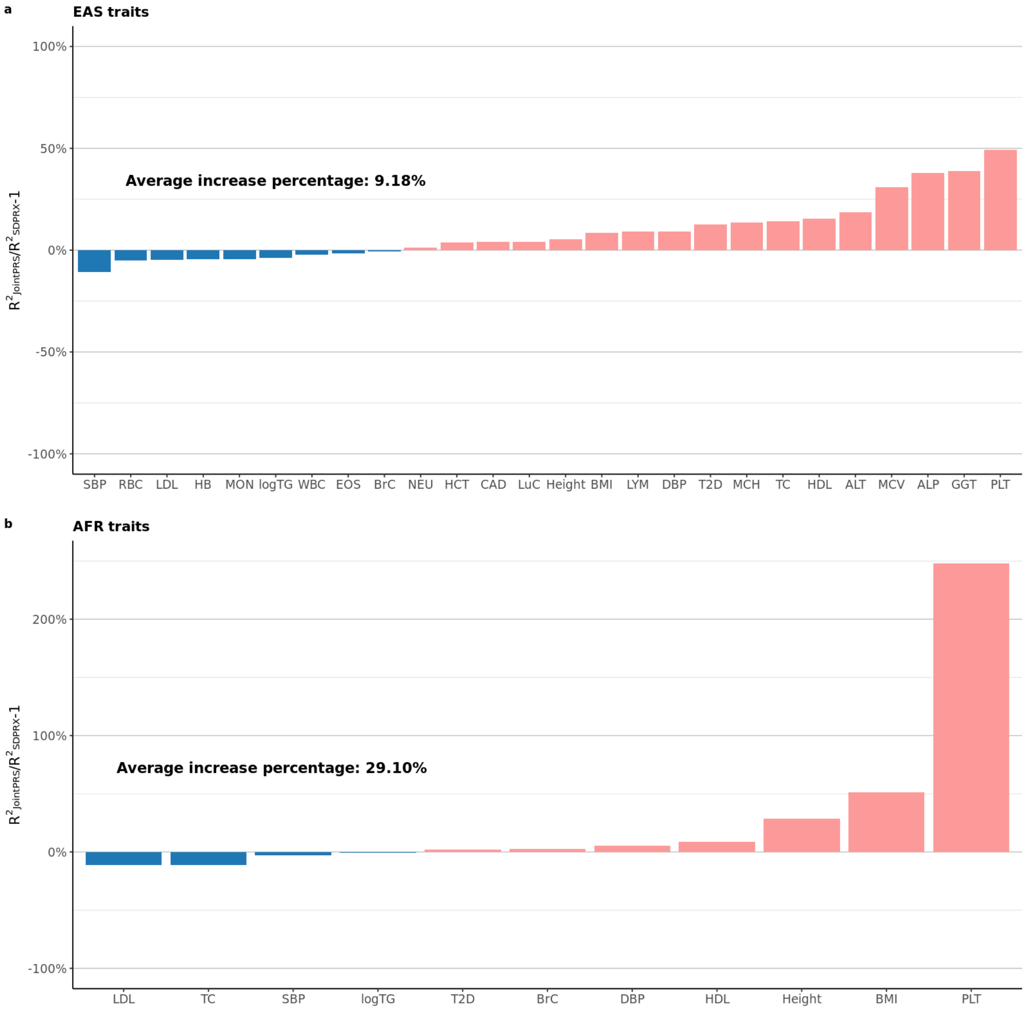

**Figure S20: Relative Improvement of JointPRS over PRS-CSx across 26 Traits and Four Non-European Populations When the Tuning and Testing Data Come From the Same Cohort (UKBB).** 26 traits from four populations were considered for the data scenarios when the tuning and testing data come from the same cohort. The whole UKBB data was spilt into five folds, and we performed 5-fold cross-validation with each fold being treated as the testing data and the rest data being treated as the tuning data. The relative improvement of the performance for JointPRS over that of PRS-CSx measured in mean of $R^2$ and AUC across five folds were calculated for quantitative traits and binary traits. The results were presented in the bar with the pink color represent positive improvement and blue color representing negative value.

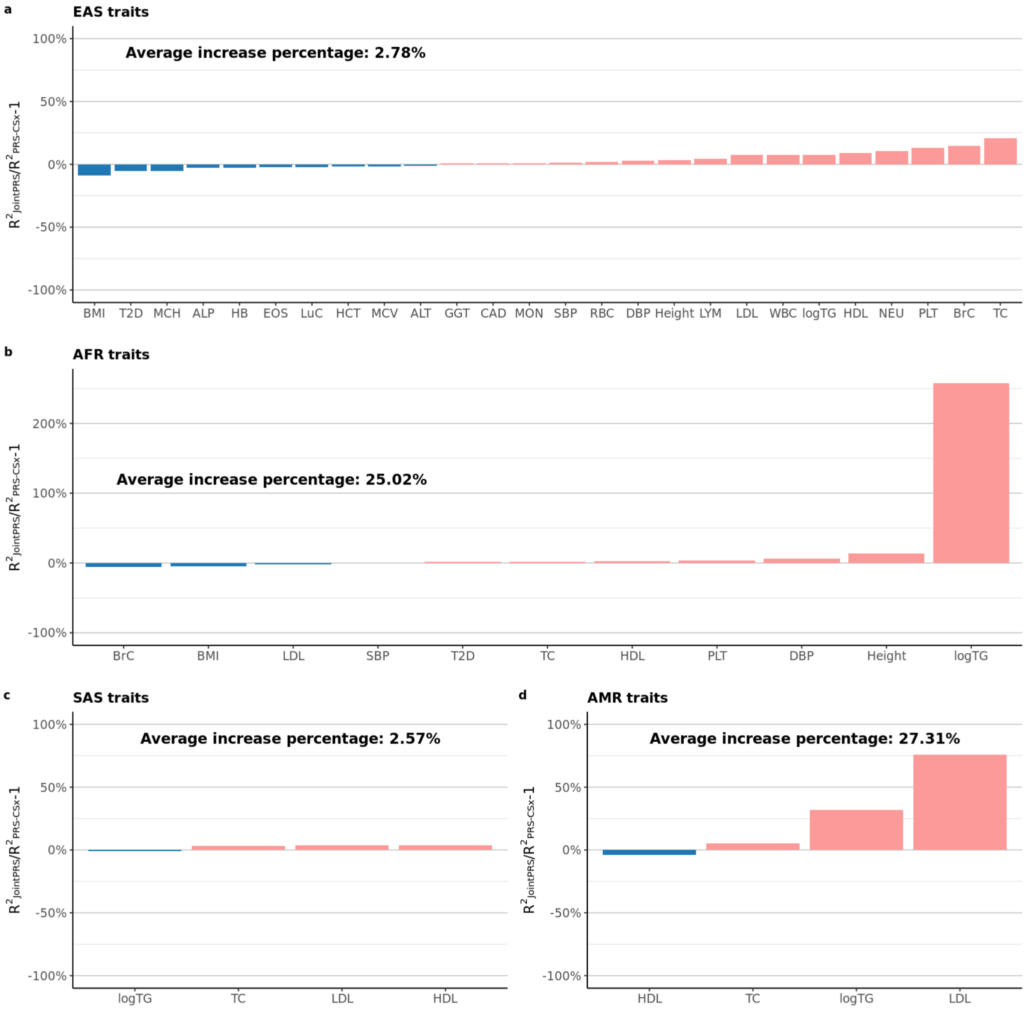

**Figure S21: Relative Improvement of JointPRS over MUSSEL across 26 Traits and Four Non-European Populations When the Tuning and Testing Data Come From the Same Cohort (UKBB).** 26 traits from four populations were considered for the data scenarios when the tuning and testing data come from the same cohort. The whole UKBB data was spilt into five folds, and we performed 5-fold cross-validation with each fold being treated as the testing data and the rest data being treated as the tuning data. The relative improvement of the performance for JointPRS over that of MUSSEL measured in mean of $R^2$ and AUC across five folds were calculated for quantitative traits and binary traits. The results were presented in the bar with the pink color represent positive improvement and blue color representing negative value.

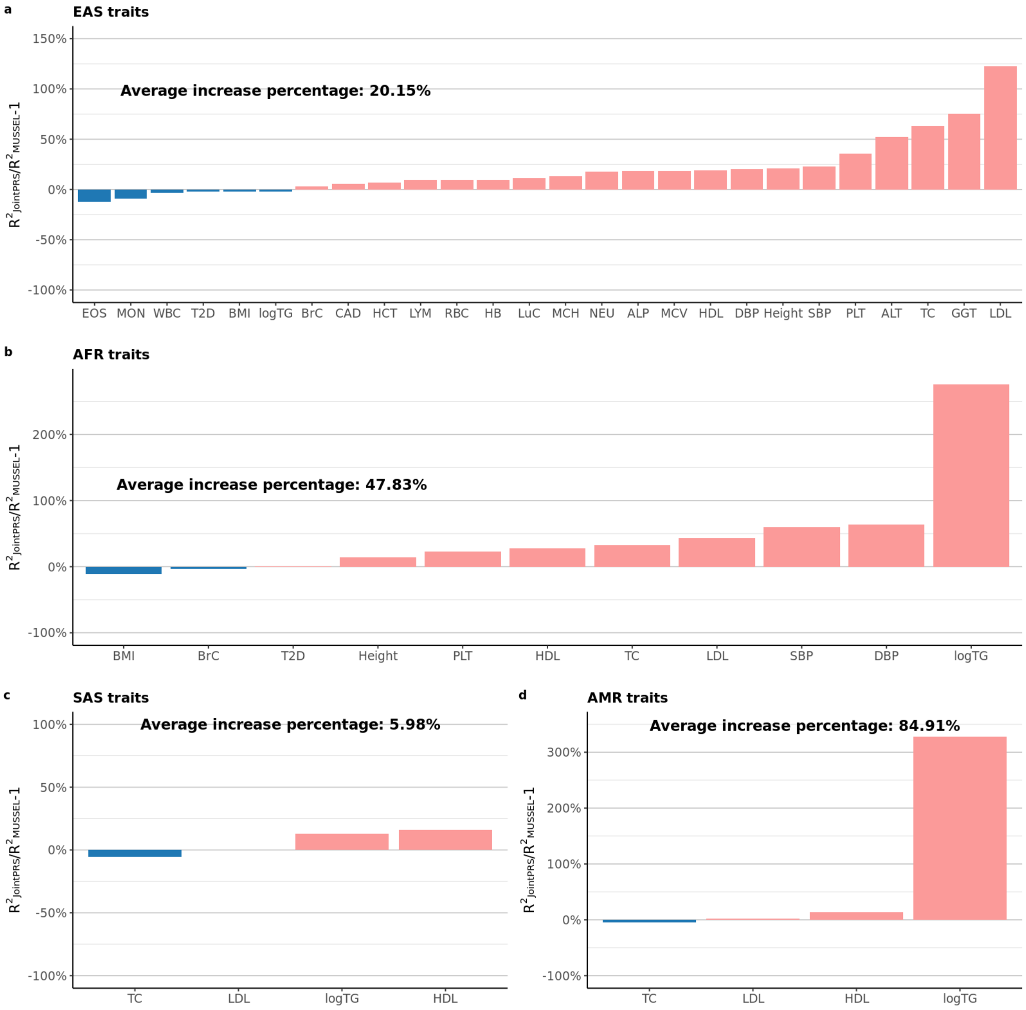

**Figure S22: Relative Improvement of JointPRS over PROSPER across 26 Traits and Four Non-European Populations When the Tuning and Testing Data Come From the Same Cohort (UKBB).** 26 traits from four populations were considered for the data scenarios when the tuning and testing data come from the same cohort. The whole UKBB data was spilt into five folds, and we performed 5-fold cross-validation with each fold being treated as the testing data and the rest data being treated as the tuning data. The relative improvement of the performance for JointPRS over that of PROSPER measured in mean of $R^2$ and AUC across five folds were calculated for quantitative traits and binary traits. The results were presented in the bar with the pink color represent positive improvement and blue color representing negative value.

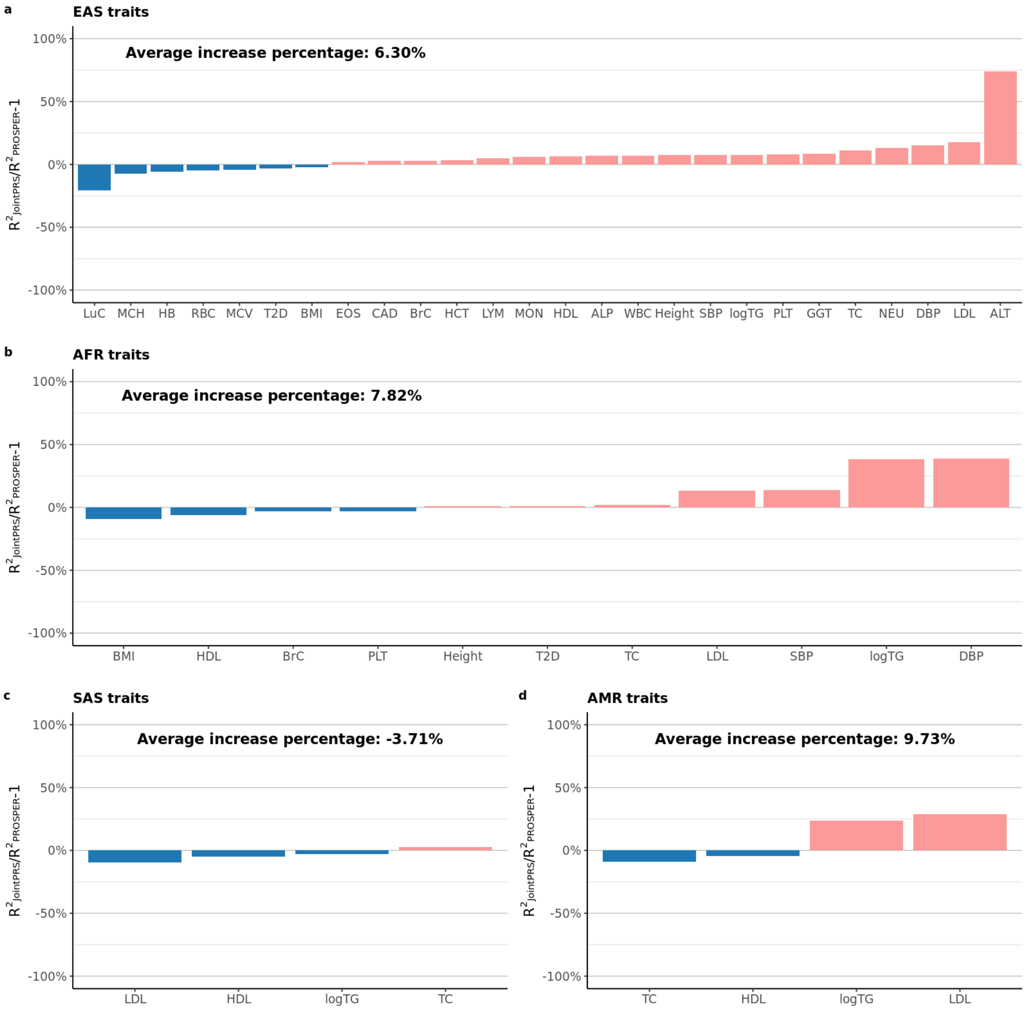

**Figure S23: Relative Improvement of JointPRS over BridgePRS across 26 Traits and Four Non-European Populations When the Tuning and Testing Data Come From the Same Cohort (UKBB).** 26 traits from four populations were considered for the data scenarios when the tuning and testing data come from the same cohort. The whole UKBB data was spilt into five folds, and we performed 5-fold cross-validation with each fold being treated as the testing data and the rest data being treated as the tuning data. The relative improvement of the performance for JointPRS over that of BridgePRS measured in mean of $R^2$ and AUC across five folds were calculated for quantitative traits and binary traits. The results were presented in the bar with the pink color represent positive improvement and blue color representing negative value.

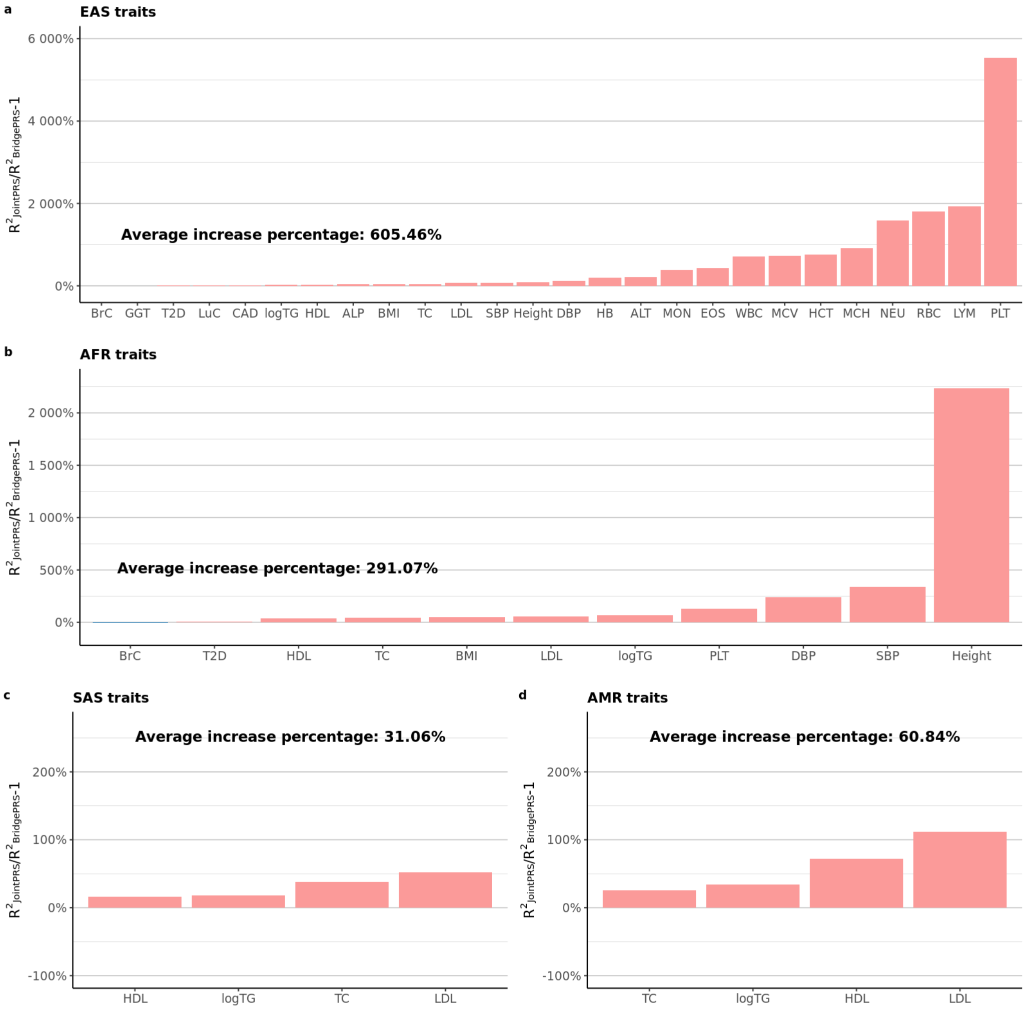

**Figure S24: Relative Improvement of the JointPRS Selected Version Over the Meta Version and the Tune Version across 26 Traits and Four Non-European Populations When the Tuning and Testing Data Come From the Same Cohort (UKBB).** 26 traits from EAS, AFR, SAS, and AMR populations were considered for the data scenarios when the tuning and testing data come from the same cohort. The whole UKBB data was spilt into five folds, and we performed 5-fold cross-validation with each fold being treated as the testing data and the rest data being treated as the tuning data. The relative improvement of the performance for the JointPRS selected version by the data-adaptive approach over that of the meta version and the tune version measured in mean of $R^2$ and AUC across five folds were calculated for quantitative traits and binary traits. The mean value across traits and five folds were presented as the black crossbar for each version

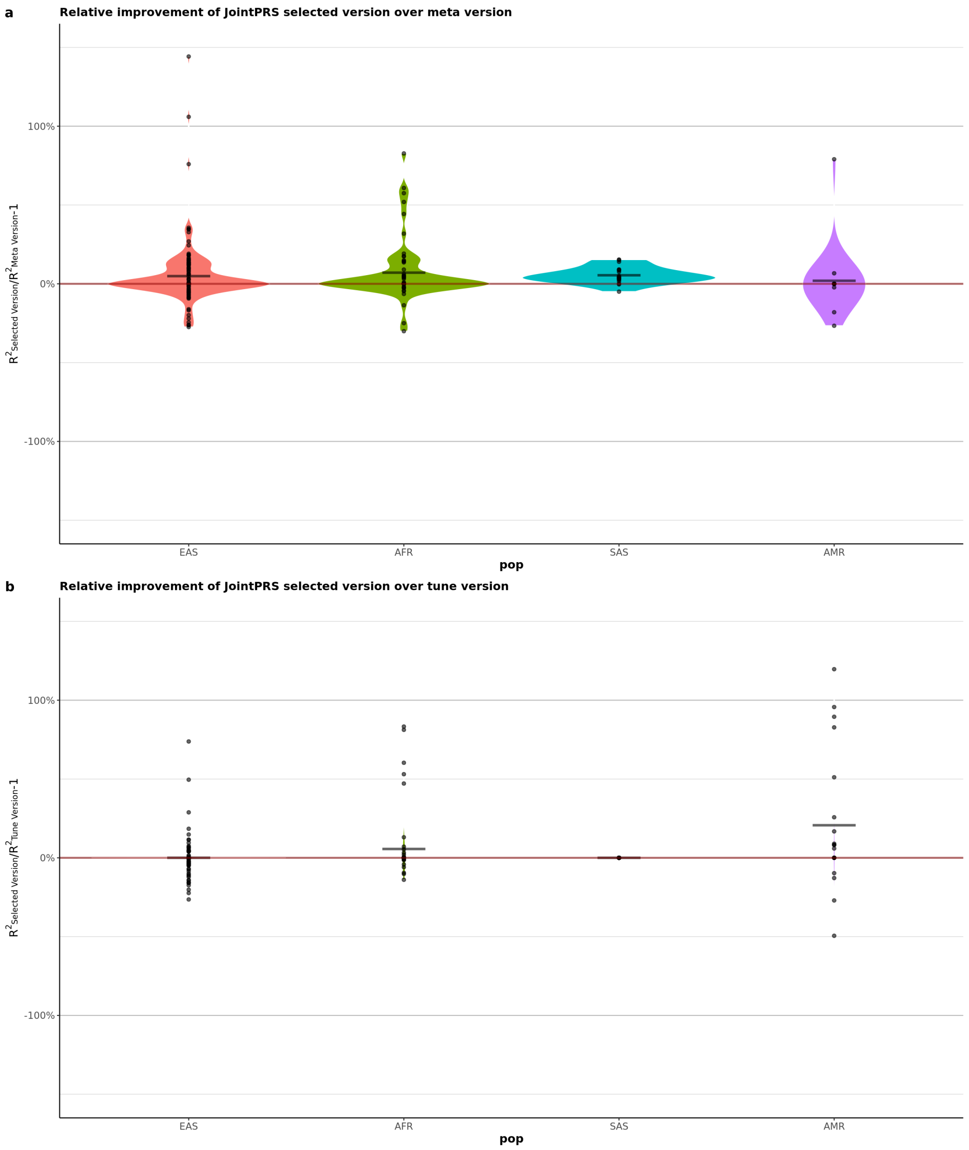

**Figure S25: Prediction Accuracy of the Meta Version and the Tune Version across GLGC Traits and Four Non-European Populations When the Tuning and Testing Data Come From the Same Cohort (UKBB).** GLGC traits from EAS, AFR, SAS, and AMR populations were considered for the data scenarios when the tuning and testing data come from the same cohort. The whole UKBB data was spilt into five folds, and we performed 5-fold cross-validation with each fold being treated as the testing data and the rest data being treated as the tuning data. The $R^2$ was used as evaluation metric for GLGC traits. The selected version obtained by the data-adaptive approach were slashed in the bar plot

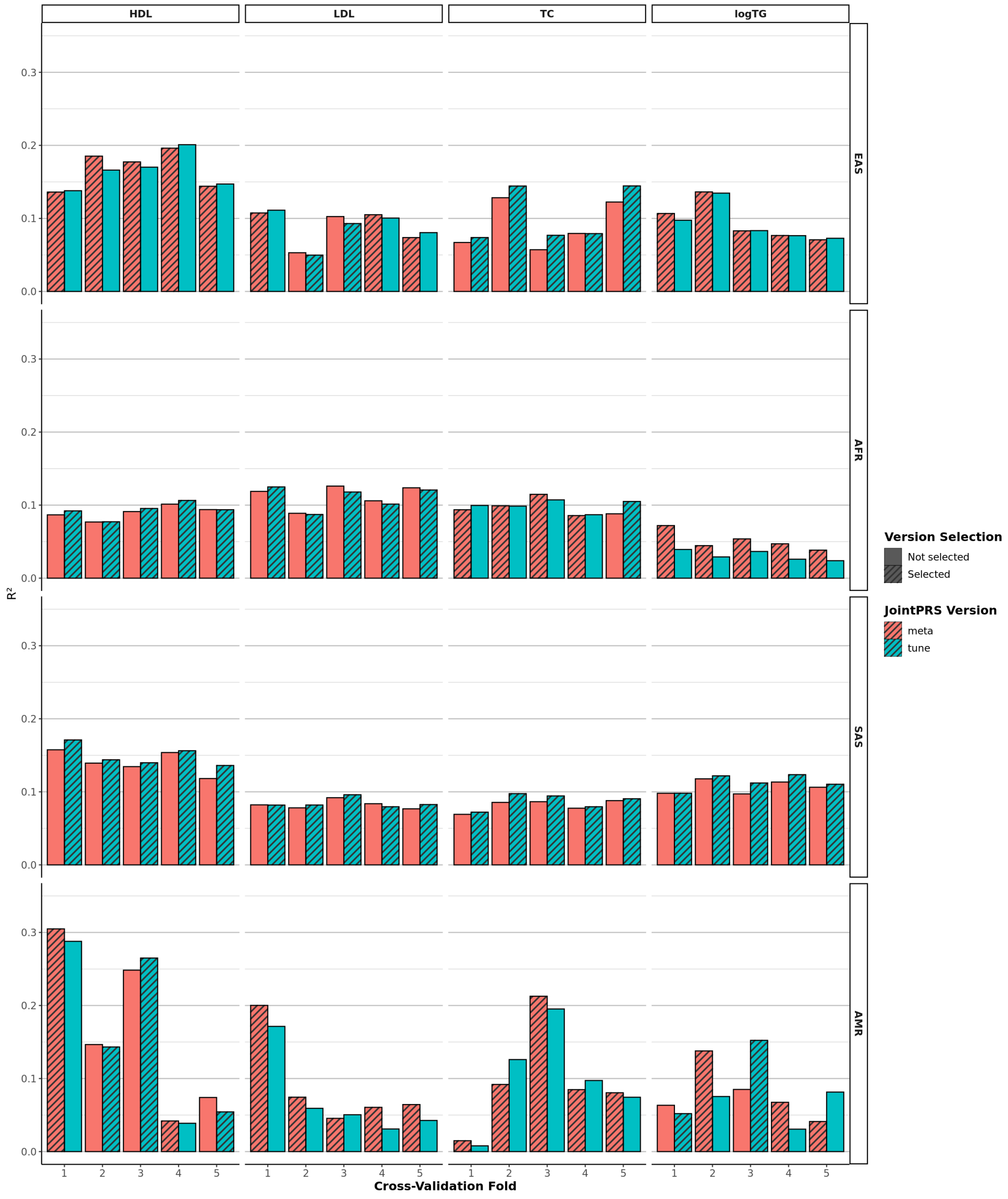

**Figure S26: Prediction Accuracy of the Meta Version and the Tune Version across PAGE Traits and Two Non-European Populations When the Tuning and Testing Data Come From the Same Cohort (UKBB).** PAGE traits from EAS and AFR populations were considered for the data scenarios when the tuning and testing data come from the same cohort. The whole UKBB data was spilt into five folds, and we performed 5-fold cross-validation with each fold being treated as the testing data and the rest data being treated as the tuning data. The $R^2$ was used as evaluation metric for PAGE traits. The selected version obtained by the data-adaptive approach were slashed in the bar plot

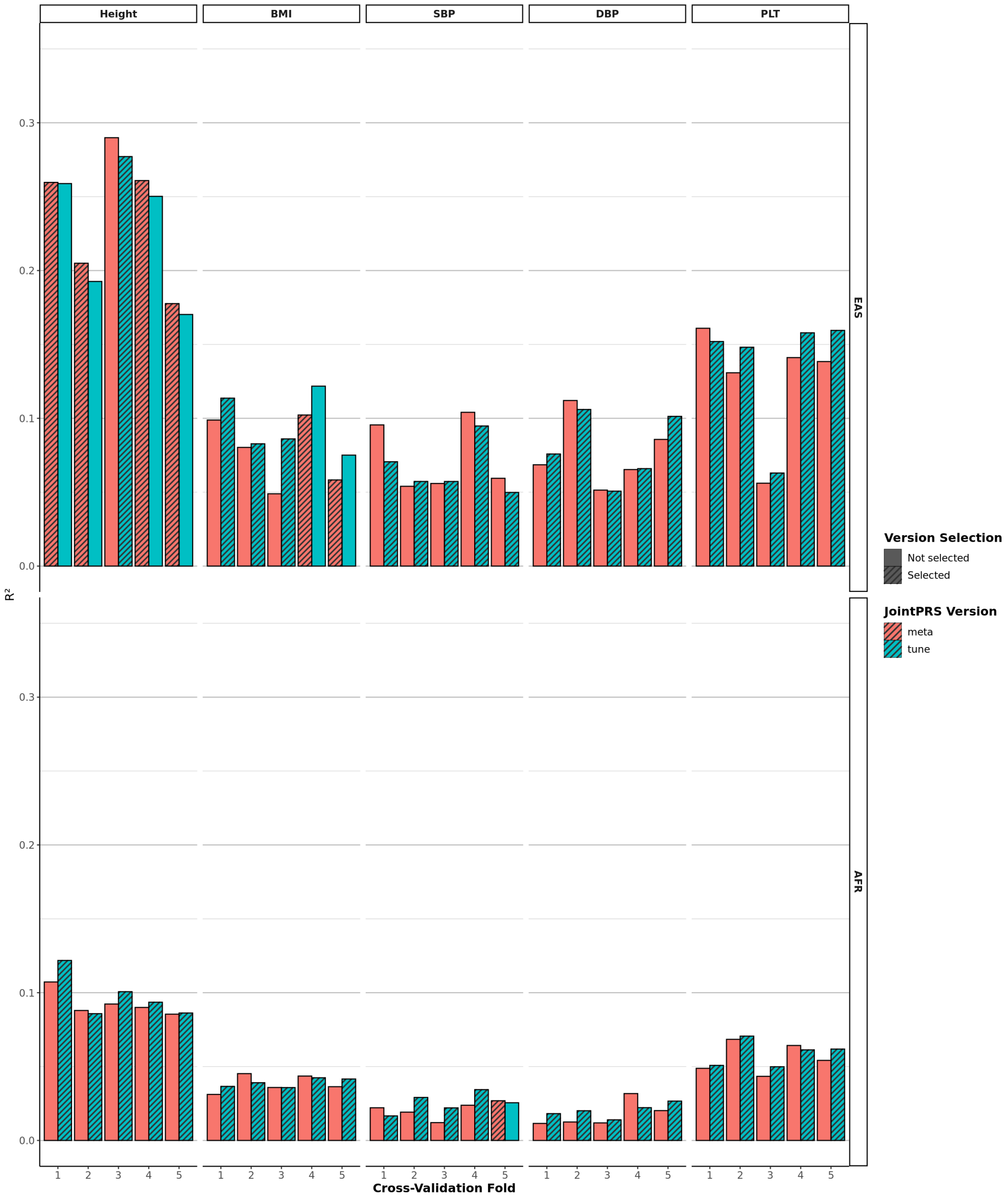

**Figure S27: Prediction Accuracy of the Meta Version and the Tune Version across BBJ Traits in EAS Populations When the Tuning and Testing Data Come From the Same Cohort (UKBB).** BBJ traits from EAS populations were considered for the data scenarios when the tuning and testing data come from the same cohort. The whole UKBB data was spilt into five folds, and we performed 5-fold cross-validation with each fold being treated as the testing data and the rest data being treated as the tuning data. The $R^2$ was used as evaluation metric for BBJ traits. The selected version obtained by the data-adaptive approach were slashed in the bar plot

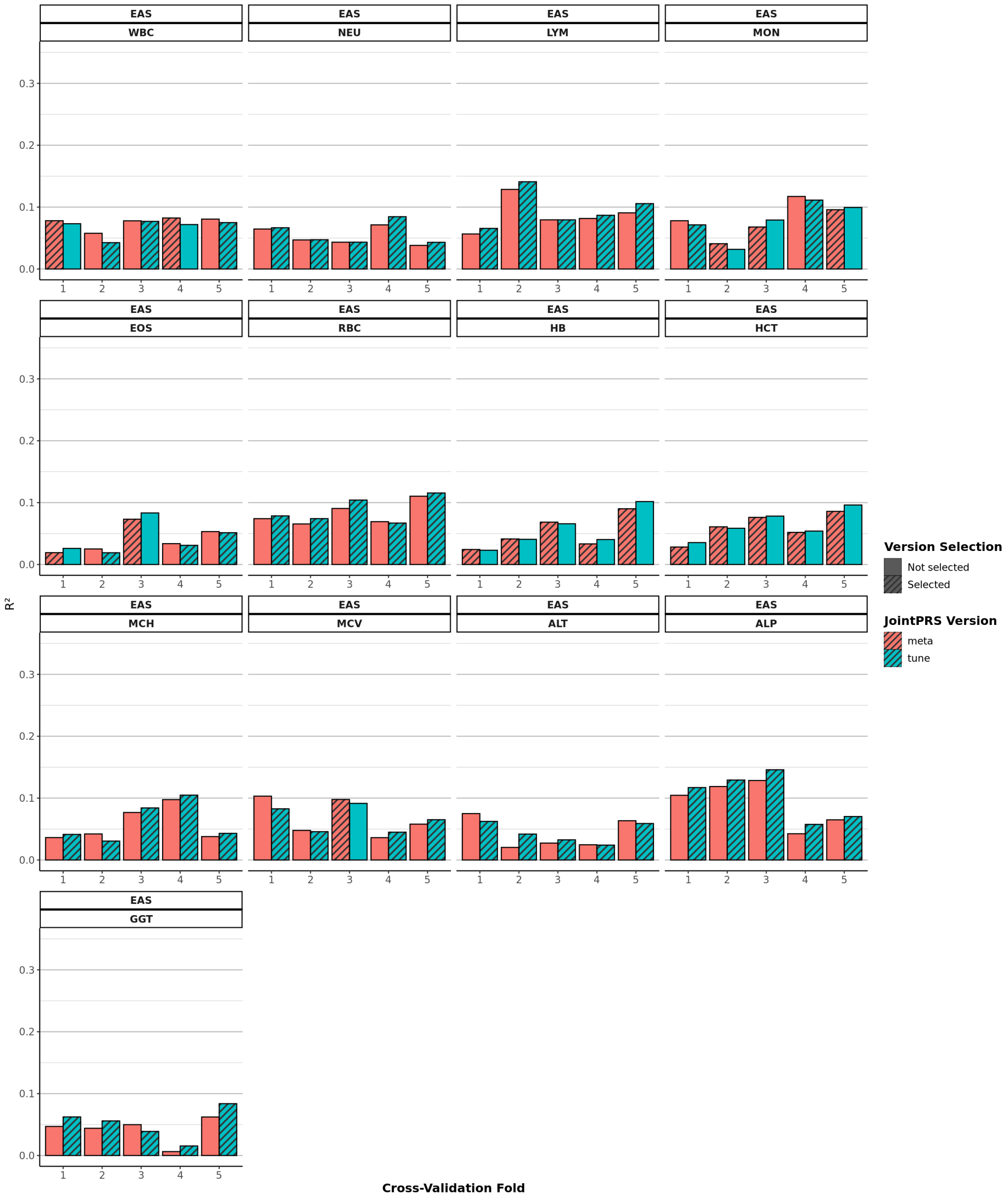

**Figure S28: Prediction Accuracy of the Meta Version and the Tune Version across Binary Traits and Two Non-European Populations When the Tuning and Testing Data Come From the Same Cohort (UKBB).** Binary traits from EAS and AFR populations were considered for the data scenarios when the tuning and testing data come from the same cohort. The whole UKBB data was spilt into five folds, and we performed 5-fold cross-validation with each fold being treated as the testing data and the rest data being treated as the tuning data. The AUC was used as evaluation metric for Binary traits. The selected version obtained by the data-adaptive approach were slashed in the bar plot

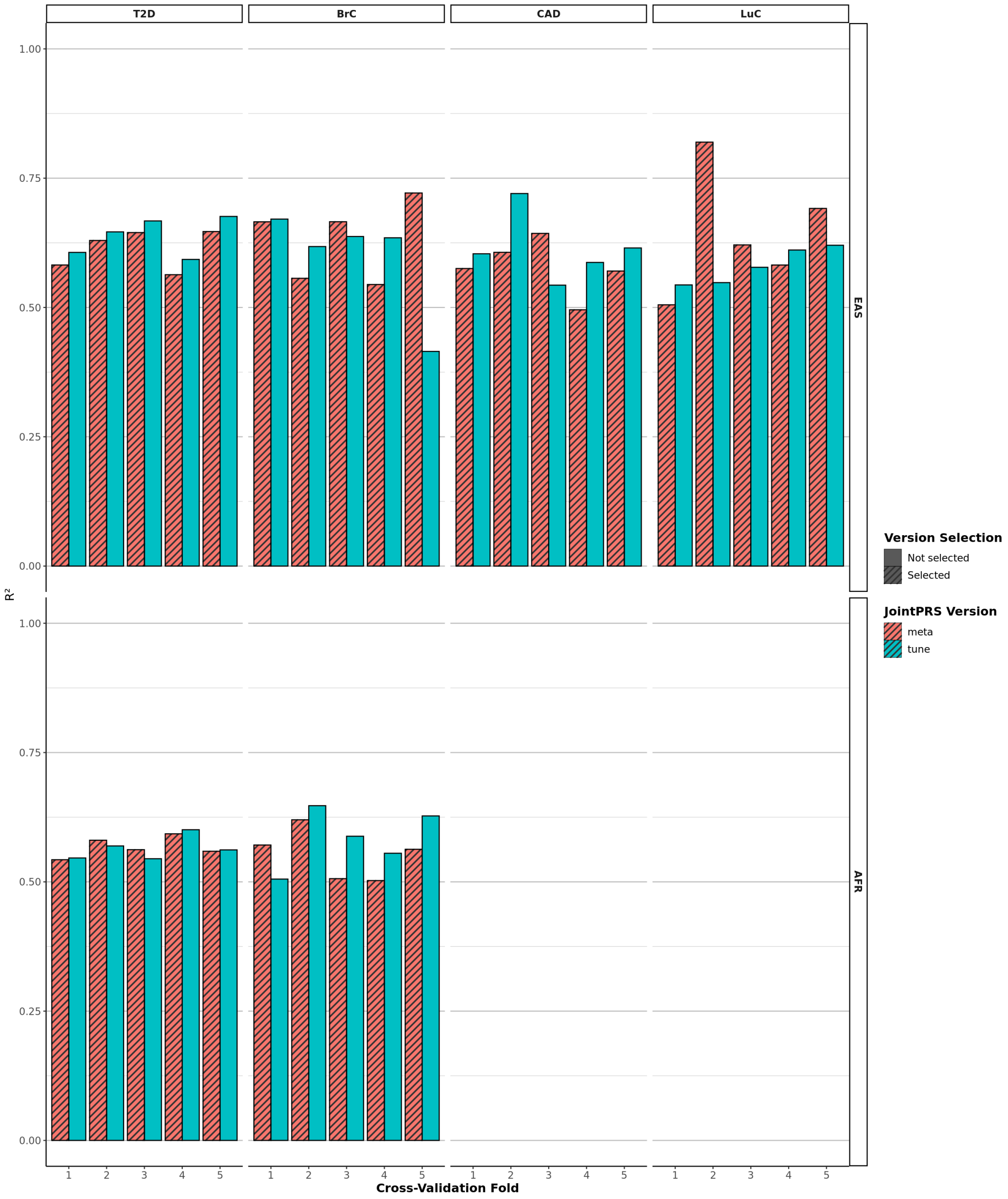

**Figure S29: Prediction Accuracy of Multi-population PRS Methods across Nine Traits and Two Non-European Populations When the Tuning and Testing Data Come From Different Cohorts (UKBB and AoU).** Nine traits from two categories we predefined were considered for the data scenarios when the tuning and testing data come from different cohorts (UKBB and AoU). The UKBB data was used as a tuning dataset, while the AoU data was used as a testing dataset. The $R^2$ were used as evaluation metric for quantitative traits and the value were presented in the bar plots for the two non-European populations (AFR and AMR). All seven methods were considered here, with the best and second-best method denoted by two stars and one star, respectively, in the corresponding bar plots. Here, SDPRX cannot provide predictions for AMR as it does not provide the corresponding LD reference panels.

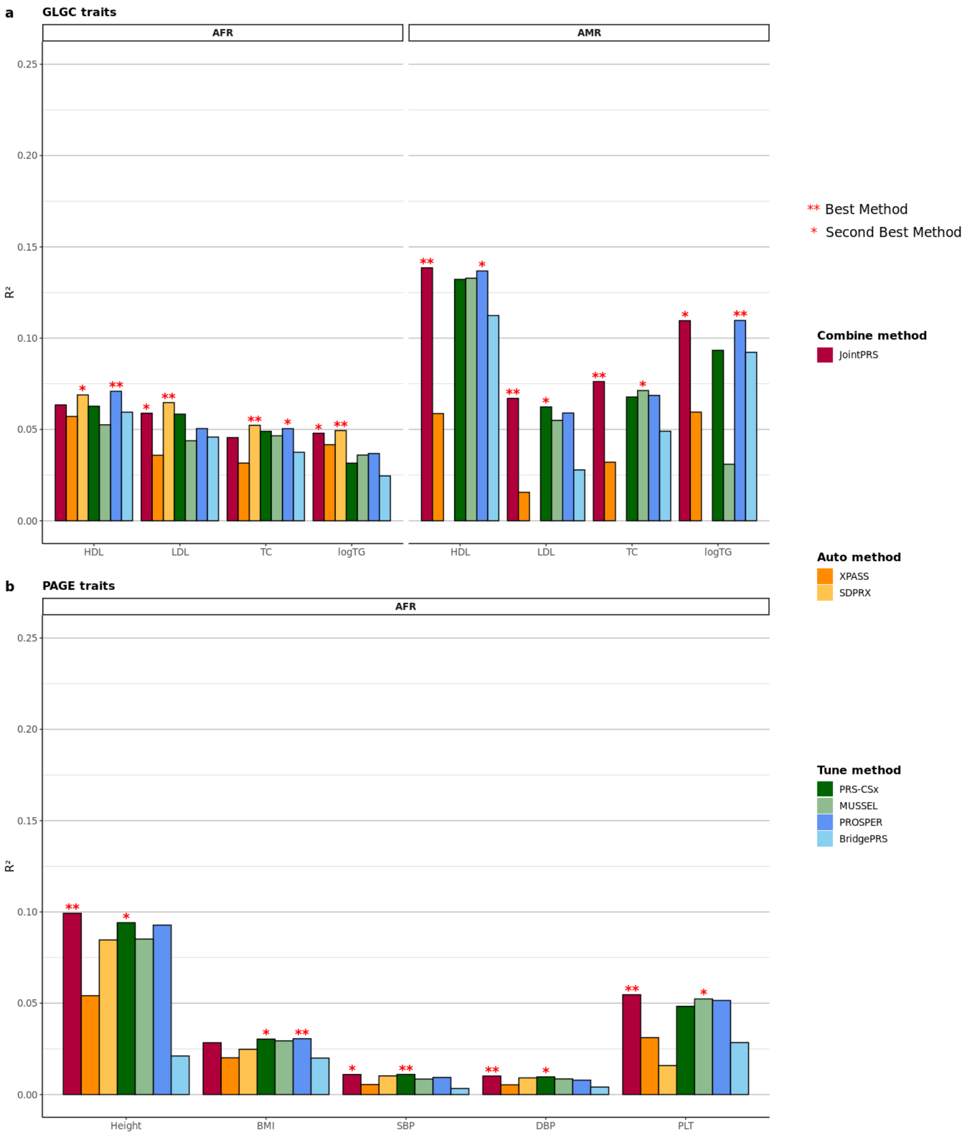

**Figure S30: Relative Improvement of JointPRS over XPASS across Nine Traits and Two Non-European Populations When the Tuning and Testing Data Come From Different Cohorts (UKBB and AoU).** Nine traits from two populations were considered for the data scenarios when the tuning and testing data come from the same cohort. The UKBB data was used as a tuning dataset, while the AoU data was used as a testing dataset. The relative improvement of the performance for JointPRS over that of XPASS measured in $R^2$ were calculated for quantitative traits. The results were presented in the bar with the pink color represent positive improvement and blue color representing negative value.

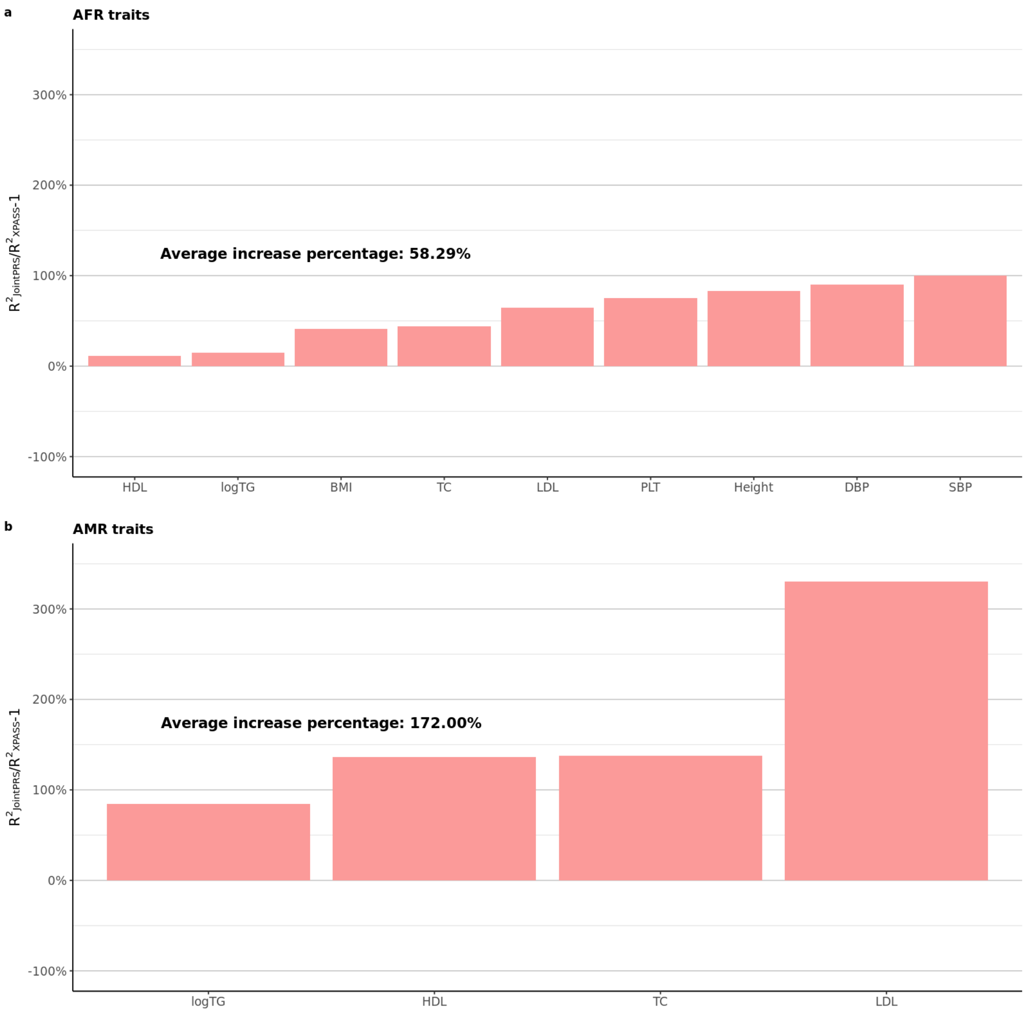

**Figure S31: Relative Improvement of JointPRS over SDPRX across Nine Traits in the AFR Population When the Tuning and Testing Data Come From Different Cohorts (UKBB and AoU).** Nine traits from the AFR population were considered for the data scenarios when the tuning and testing data come from the same cohort. The UKBB data was used as a tuning dataset, while the AoU data was used as a testing dataset. The relative improvement of the performance for JointPRS over that of SDPRX measured in $R^2$ were calculated for quantitative traits. The results were presented in the bar with the pink color represent positive improvement and blue color representing negative value. Here, SDPRX cannot provide predictions for AMR as it does not provide the corresponding LD reference panels.

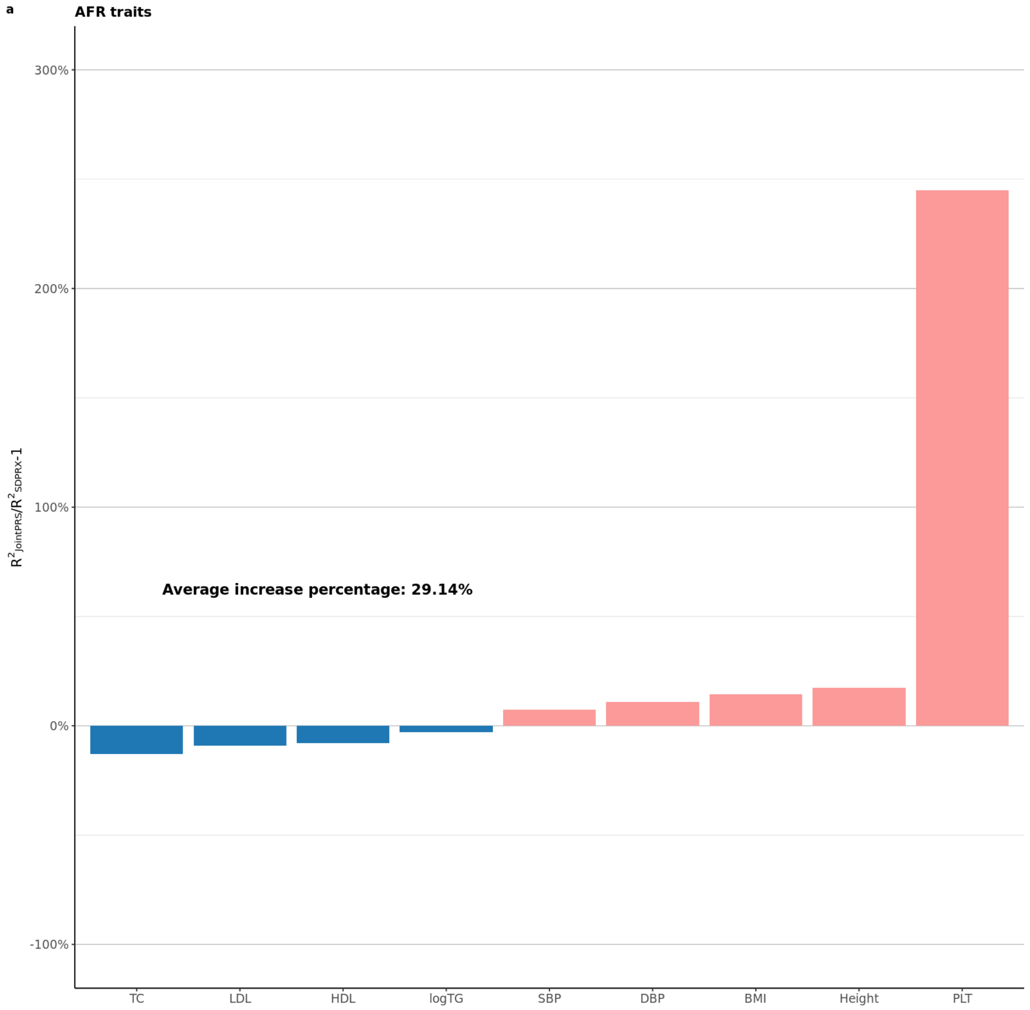

**Figure S32: Relative Improvement of JointPRS over PRS-CSx across Nine Traits and Two Non-European Populations When the Tuning and Testing Data Come From Different Cohorts (UKBB and AoU).** Nine traits from two populations were considered for the data scenarios when the tuning and testing data come from the same cohort. The UKBB data was used as a tuning dataset, while the AoU data was used as a testing dataset. The relative improvement of the performance for JointPRS over that of PRS-CSx measured in $R^2$ were calculated for quantitative traits. The results were presented in the bar with the pink color represent positive improvement and blue color representing negative value.

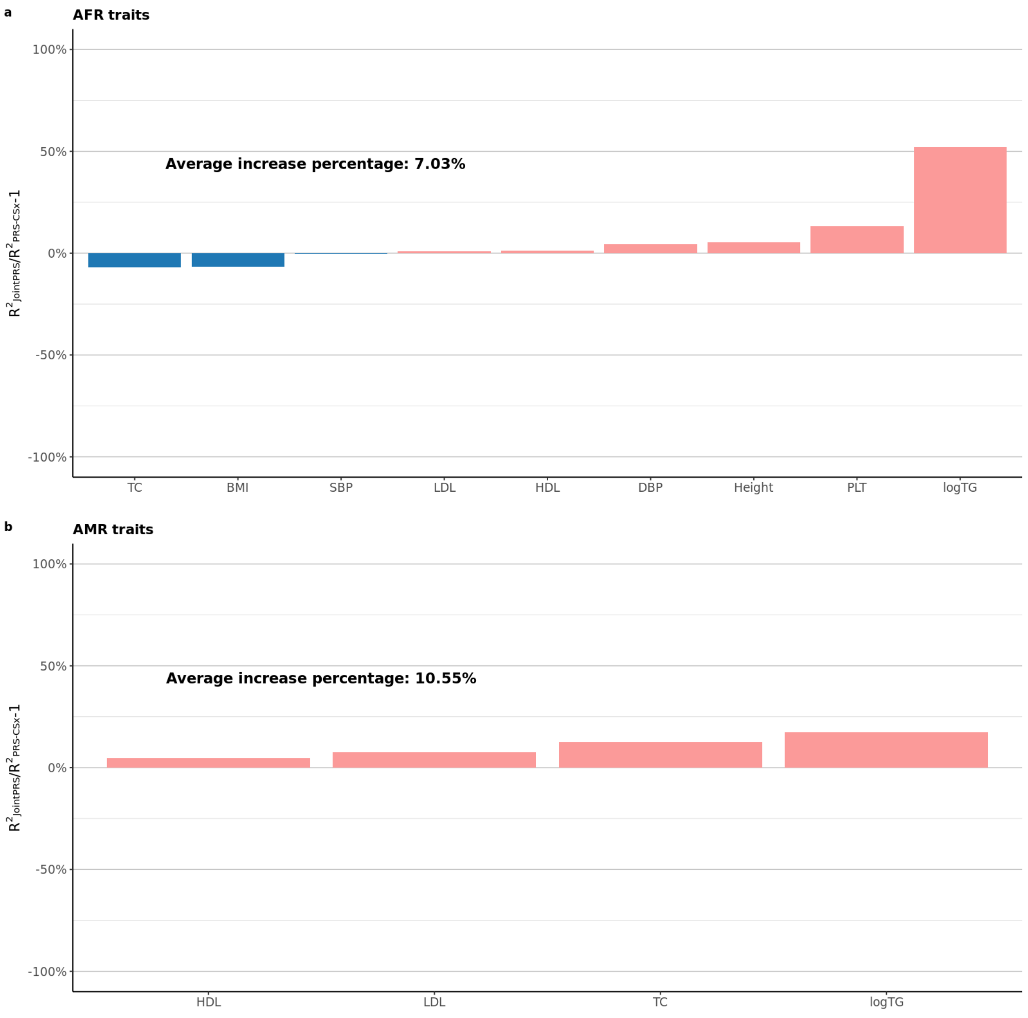

**Figure S33: Relative Improvement of JointPRS over MUSSEL across Nine Traits and Two Non-European Populations When the Tuning and Testing Data Come From Different Cohorts (UKBB and AoU).** Nine traits from two populations were considered for the data scenarios when the tuning and testing data come from the same cohort. The UKBB data was used as a tuning dataset, while the AoU data was used as a testing dataset. The relative improvement of the performance for JointPRS over that of MUSSEL measured in $R^2$ were calculated for quantitative traits. The results were presented in the bar with the pink color represent positive improvement and blue color representing negative value.

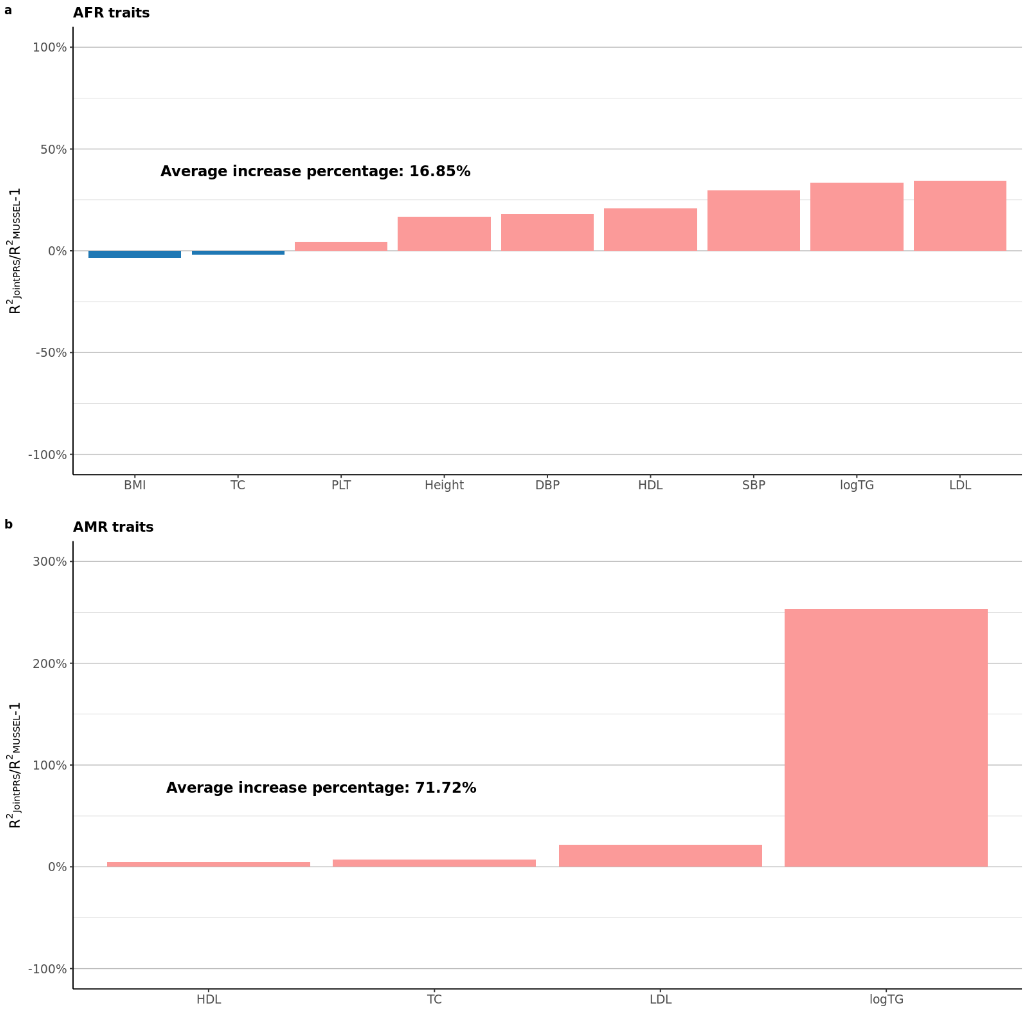

**Figure S34: Relative Improvement of JointPRS over PROSPER across Nine Traits and Two Non-European Populations When the Tuning and Testing Data Come From Different Cohorts (UKBB and AoU).** Nine traits from two populations were considered for the data scenarios when the tuning and testing data come from the same cohort. The UKBB data was used as a tuning dataset, while the AoU data was used as a testing dataset. The relative improvement of the performance for JointPRS over that of PROSPER measured in $R^2$ were calculated for quantitative traits. The results were presented in the bar with the pink color represent positive improvement and blue color representing negative value.

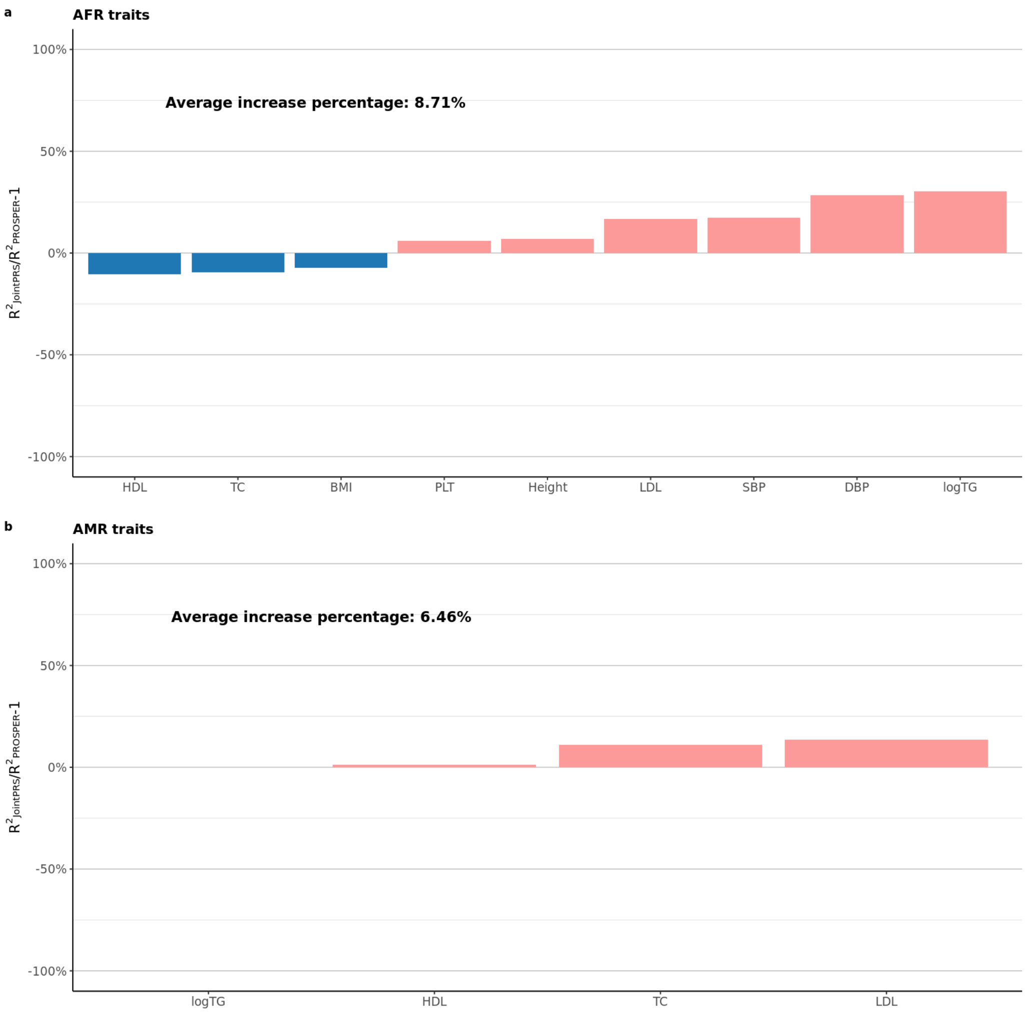

**Figure S35: Relative Improvement of JointPRS over BridgePRS across Nine Traits and Two Non-European Populations When the Tuning and Testing Data Come From Different Cohorts (UKBB and AoU).** Nine traits from two populations were considered for the data scenarios when the tuning and testing data come from the same cohort. The UKBB data was used as a tuning dataset, while the AoU data was used as a testing dataset. The relative improvement of the performance for JointPRS over that of BridgePRS measured in $R^2$ were calculated for quantitative traits. The results were presented in the bar with the pink color represent positive improvement and blue color representing negative value.

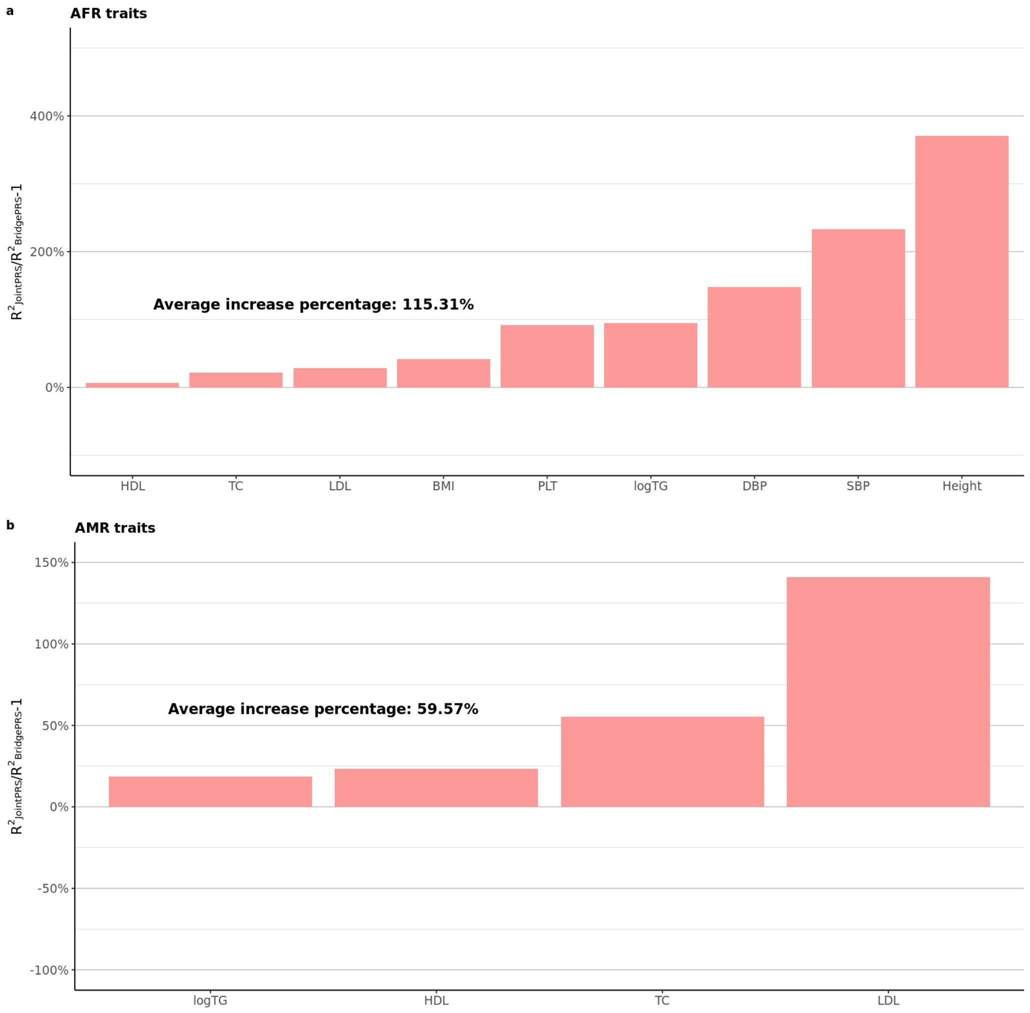

**Table S1: Simulation Results for the Mean and Standard Deviation $R^2$ Values across Five Replications When a Tuning Dataset is Not Available in EAS, AFR, SAS, and AMR Populations.**

| p | Non-European Sample Sizes | Method | EAS | | AFR | | SAS | | AMR | |
|---|---|---|---|---|---|---|---|---|---|---|
| | | | Mean | SD | Mean | SD | Mean | SD | Mean | SD |
| 0.1 | 80,000 | JointPRS-auto | 0.173 | 0.007 | 0.126 | 0.004 | 0.175 | 0.003 | 0.175 | 0.004 |
| | | XPASS | 0.123 | 0.008 | 0.087 | 0.004 | 0.136 | 0.006 | 0.087 | 0.006 |
| | | SDPRX | 0.148 | 0.007 | 0.112 | 0.005 | N/A | N/A | N/A | N/A |
| | | PRS-CSx-auto | 0.124 | 0.005 | 0.085 | 0.005 | 0.126 | 0.003 | 0.123 | 0.002 |
| | 15,000 | JointPRS-auto | 0.082 | 0.005 | 0.049 | 0.005 | 0.088 | 0.003 | 0.091 | 0.006 |
| | | XPASS | 0.093 | 0.006 | 0.064 | 0.004 | 0.100 | 0.006 | 0.093 | 0.005 |
| | | SDPRX | 0.100 | 0.005 | 0.068 | 0.006 | N/A | N/A | N/A | N/A |
| | | PRS-CSx-auto | 0.040 | 0.003 | 0.022 | 0.004 | 0.041 | 0.005 | 0.045 | 0.004 |
| 0.01 | 80,000 | JointPRS-auto | 0.233 | 0.013 | 0.206 | 0.009 | 0.238 | 0.005 | 0.239 | 0.006 |
| | | XPASS | 0.096 | 0.007 | 0.065 | 0.004 | 0.112 | 0.010 | 0.037 | 0.004 |
| | | SDPRX | 0.206 | 0.013 | 0.173 | 0.014 | N/A | N/A | N/A | N/A |
| | | PRS-CSx-auto | 0.204 | 0.011 | 0.177 | 0.009 | 0.203 | 0.006 | 0.205 | 0.005 |
| | 15,000 | JointPRS-auto | 0.143 | 0.011 | 0.112 | 0.009 | 0.147 | 0.002 | 0.150 | 0.009 |
| | | XPASS | 0.058 | 0.005 | 0.039 | 0.002 | 0.065 | 0.007 | 0.057 | 0.006 |
| | | SDPRX | 0.147 | 0.009 | 0.117 | 0.009 | N/A | N/A | N/A | N/A |
| | | PRS-CSx-auto | 0.088 | 0.007 | 0.065 | 0.004 | 0.089 | 0.002 | 0.092 | 0.008 |
| 0.001 | 80,000 | JointPRS-auto | 0.307 | 0.013 | 0.288 | 0.012 | 0.310 | 0.009 | 0.305 | 0.006 |
| | | XPASS | 0.227 | 0.018 | 0.217 | 0.011 | 0.224 | 0.010 | 0.224 | 0.005 |
| | | SDPRX | 0.310 | 0.014 | 0.296 | 0.011 | N/A | N/A | N/A | N/A |
| | | PRS-CSx-auto | 0.306 | 0.015 | 0.291 | 0.009 | 0.309 | 0.005 | 0.299 | 0.008 |
| | 15,000 | JointPRS-auto | 0.250 | 0.012 | 0.221 | 0.012 | 0.252 | 0.007 | 0.246 | 0.009 |
| | | XPASS | 0.026 | 0.010 | 0.023 | 0.008 | 0.029 | 0.004 | 0.025 | 0.006 |
| | | SDPRX | 0.250 | 0.013 | 0.225 | 0.012 | N/A | N/A | N/A | N/A |
| | | PRS-CSx-auto | 0.221 | 0.012 | 0.196 | 0.009 | 0.224 | 0.008 | 0.217 | 0.008 |
| $5 \times 10^{-4}$ | 80,000 | JointPRS-auto | 0.325 | 0.007 | 0.293 | 0.012 | 0.320 | 0.010 | 0.320 | 0.004 |
| | | XPASS | 0.278 | 0.011 | 0.264 | 0.010 | 0.265 | 0.014 | 0.283 | 0.007 |
| | | SDPRX | 0.329 | 0.012 | 0.305 | 0.008 | N/A | N/A | N/A | N/A |
| | | PRS-CSx-auto | 0.324 | 0.008 | 0.295 | 0.013 | 0.319 | 0.011 | 0.314 | 0.004 |
| | 15,000 | JointPRS-auto | 0.273 | 0.009 | 0.241 | 0.010 | 0.273 | 0.010 | 0.272 | 0.008 |
| | | XPASS | 0.083 | 0.011 | 0.075 | 0.004 | 0.082 | 0.009 | 0.087 | 0.009 |
| | | SDPRX | 0.278 | 0.012 | 0.244 | 0.010 | N/A | N/A | N/A | N/A |
| | | PRS-CSx-auto | 0.252 | 0.012 | 0.226 | 0.010 | 0.254 | 0.009 | 0.251 | 0.008 |

**Table S2: Simulation Results for the Mean and Standard Deviation $R^2$ Values across Five Replications When a Tuning Dataset is Available in EAS, AFR, SAS, and AMR Populations for Causal SNP Proportions $p = 0.1$.**

| p | minor_sample | minor_val_sample | method | EAS | | AFR | | SAS | | AMR | |
|---|---|---|---|---|---|---|---|---|---|---|---|
| | | | | Mean | SD | Mean | SD | Mean | SD | Mean | SD |
| 0.1 | 80,000 | 500 | JointPRS | 0.173 | 0.007 | 0.126 | 0.004 | 0.175 | 0.003 | 0.175 | 0.004 |
| | | | XPASS | 0.123 | 0.008 | 0.087 | 0.004 | 0.136 | 0.006 | 0.087 | 0.006 |
| | | | SDPRX | 0.148 | 0.007 | 0.112 | 0.005 | N/A | N/A | N/A | N/A |
| | | | PRS-CSx | 0.160 | 0.008 | 0.119 | 0.008 | 0.169 | 0.010 | 0.164 | 0.007 |
| | | | MUSSEL | 0.143 | 0.021 | 0.126 | 0.010 | 0.142 | 0.013 | 0.159 | 0.009 |
| | | | PROSPER | 0.148 | 0.009 | 0.115 | 0.006 | 0.154 | 0.011 | 0.153 | 0.011 |
| | | | BridgePRS | 0.099 | 0.015 | 0.069 | 0.011 | 0.114 | 0.006 | 0.116 | 0.012 |
| | | 2,000 | JointPRS | 0.173 | 0.007 | 0.126 | 0.004 | 0.175 | 0.003 | 0.176 | 0.003 |
| | | | XPASS | 0.123 | 0.008 | 0.087 | 0.004 | 0.136 | 0.006 | 0.087 | 0.006 |
| | | | SDPRX | 0.148 | 0.007 | 0.112 | 0.005 | N/A | N/A | N/A | N/A |
| | | | PRS-CSx | 0.172 | 0.006 | 0.132 | 0.004 | 0.176 | 0.006 | 0.177 | 0.002 |
| | | | MUSSEL | 0.134 | 0.075 | 0.103 | 0.058 | 0.123 | 0.067 | 0.135 | 0.075 |
| | | | PROSPER | 0.161 | 0.008 | 0.120 | 0.014 | 0.165 | 0.007 | 0.167 | 0.005 |
| | | | BridgePRS | 0.120 | 0.006 | 0.079 | 0.011 | 0.119 | 0.012 | 0.129 | 0.004 |
| | | 5,000 | JointPRS | 0.176 | 0.007 | 0.128 | 0.005 | 0.179 | 0.002 | 0.181 | 0.003 |
| | | | XPASS | 0.123 | 0.007 | 0.088 | 0.005 | 0.138 | 0.004 | 0.087 | 0.011 |
| | | | SDPRX | 0.151 | 0.006 | 0.112 | 0.005 | N/A | N/A | N/A | N/A |
| | | | PRS-CSx | 0.173 | 0.006 | 0.132 | 0.004 | 0.178 | 0.004 | 0.179 | 0.002 |
| | | | MUSSEL | 0.100 | 0.089 | 0.080 | 0.072 | 0.090 | 0.081 | 0.104 | 0.093 |
| | | | PROSPER | 0.132 | 0.073 | 0.104 | 0.046 | 0.167 | 0.004 | 0.138 | 0.077 |
| | | | BridgePRS | 0.122 | 0.005 | 0.085 | 0.004 | 0.125 | 0.006 | 0.131 | 0.004 |
| | | 10,000 | JointPRS | 0.180 | 0.006 | 0.131 | 0.005 | 0.182 | 0.004 | 0.181 | 0.001 |
| | | | XPASS | 0.127 | 0.007 | 0.090 | 0.005 | 0.140 | 0.007 | 0.082 | 0.010 |
| | | | SDPRX | 0.154 | 0.007 | 0.115 | 0.006 | N/A | N/A | N/A | N/A |
| | | | PRS-CSx | 0.174 | 0.006 | 0.133 | 0.004 | 0.178 | 0.004 | 0.179 | 0.002 |
| | | | MUSSEL | 0.167 | 0.009 | 0.133 | 0.005 | 0.150 | 0.009 | 0.173 | 0.005 |
| | | | PROSPER | 0.122 | 0.074 | 0.094 | 0.054 | 0.127 | 0.069 | 0.131 | 0.075 |
| | | | BridgePRS | 0.123 | 0.005 | 0.089 | 0.004 | 0.126 | 0.006 | 0.132 | 0.004 |
| | 15,000 | 500 | JointPRS | 0.098 | 0.018 | 0.058 | 0.012 | 0.113 | 0.011 | 0.105 | 0.018 |
| | | | XPASS | 0.093 | 0.006 | 0.064 | 0.004 | 0.100 | 0.006 | 0.093 | 0.005 |
| | | | SDPRX | 0.100 | 0.005 | 0.068 | 0.006 | N/A | N/A | N/A | N/A |
| | | | PRS-CSx | 0.111 | 0.004 | 0.074 | 0.014 | 0.120 | 0.005 | 0.113 | 0.012 |
| | | | MUSSEL | 0.091 | 0.019 | 0.072 | 0.009 | 0.105 | 0.006 | 0.116 | 0.008 |
| | | | PROSPER | 0.096 | 0.014 | 0.070 | 0.008 | 0.109 | 0.006 | 0.066 | 0.050 |
| | | | BridgePRS | 0.070 | 0.014 | 0.050 | 0.007 | 0.081 | 0.005 | 0.088 | 0.006 |
| | | 2,000 | JointPRS | 0.113 | 0.006 | 0.077 | 0.008 | 0.121 | 0.004 | 0.122 | 0.005 |
| | | | XPASS | 0.093 | 0.006 | 0.064 | 0.004 | 0.100 | 0.006 | 0.093 | 0.005 |
| | | | SDPRX | 0.100 | 0.005 | 0.068 | 0.006 | N/A | N/A | N/A | N/A |
| | | | PRS-CSx | 0.114 | 0.005 | 0.080 | 0.007 | 0.122 | 0.005 | 0.124 | 0.007 |
| | | | MUSSEL | 0.106 | 0.006 | 0.077 | 0.006 | 0.109 | 0.005 | 0.116 | 0.008 |
| | | | PROSPER | 0.095 | 0.025 | 0.073 | 0.005 | 0.114 | 0.006 | 0.109 | 0.028 |
| | | | BridgePRS | 0.082 | 0.004 | 0.057 | 0.004 | 0.091 | 0.003 | 0.095 | 0.004 |
| | | 5,000 | JointPRS | 0.114 | 0.006 | 0.078 | 0.008 | 0.121 | 0.003 | 0.124 | 0.004 |
| | | | XPASS | 0.095 | 0.005 | 0.066 | 0.005 | 0.102 | 0.005 | 0.094 | 0.005 |
| | | | SDPRX | 0.106 | 0.003 | 0.072 | 0.005 | N/A | N/A | N/A | N/A |
| | | | PRS-CSx | 0.115 | 0.005 | 0.081 | 0.008 | 0.123 | 0.004 | 0.126 | 0.006 |
| | | | MUSSEL | 0.099 | 0.026 | 0.067 | 0.020 | 0.100 | 0.029 | 0.106 | 0.034 |
| | | | PROSPER | 0.099 | 0.026 | 0.062 | 0.029 | 0.100 | 0.041 | 0.117 | 0.008 |
| | | | BridgePRS | 0.087 | 0.006 | 0.060 | 0.002 | 0.093 | 0.004 | 0.098 | 0.005 |
| | | 10,000 | JointPRS | 0.114 | 0.005 | 0.078 | 0.008 | 0.121 | 0.003 | 0.124 | 0.004 |
| | | | XPASS | 0.097 | 0.005 | 0.067 | 0.005 | 0.106 | 0.007 | 0.095 | 0.005 |
| | | | SDPRX | 0.112 | 0.002 | 0.074 | 0.008 | N/A | N/A | N/A | N/A |
| | | | PRS-CSx | 0.116 | 0.005 | 0.081 | 0.008 | 0.123 | 0.004 | 0.126 | 0.006 |
| | | | MUSSEL | 0.094 | 0.028 | 0.056 | 0.022 | 0.094 | 0.021 | 0.087 | 0.030 |
| | | | PROSPER | 0.087 | 0.047 | 0.045 | 0.041 | 0.097 | 0.042 | 0.090 | 0.052 |
| | | | BridgePRS | 0.090 | 0.004 | 0.061 | 0.003 | 0.094 | 0.004 | 0.099 | 0.004 |

**Table S3: Simulation Results for the Mean and Standard Deviation $R^2$ Values across Five Replications When a Tuning Dataset is Available in EAS, AFR, SAS, and AMR Populations for Causal SNP Proportions $p = 0.01$.**

| p | minor_sample | minor_val_sample | method | EAS | | AFR | | SAS | | AMR | |
|---|---|---|---|---|---|---|---|---|---|---|---|
| | | | | Mean | SD | Mean | SD | Mean | SD | Mean | SD |
| 0.01 | 80,000 | 500 | JointPRS | 0.233 | 0.013 | 0.206 | 0.009 | 0.238 | 0.005 | 0.239 | 0.006 |
| | | | XPASS | 0.096 | 0.007 | 0.065 | 0.004 | 0.112 | 0.010 | 0.037 | 0.004 |
| | | | SDPRX | 0.206 | 0.013 | 0.173 | 0.014 | N/A | N/A | N/A | N/A |
| | | | PRS-CSx | 0.219 | 0.011 | 0.188 | 0.009 | 0.222 | 0.008 | 0.220 | 0.012 |
| | | | MUSSEL | 0.190 | 0.015 | 0.196 | 0.012 | 0.158 | 0.040 | 0.200 | 0.007 |
| | | | PROSPER | 0.190 | 0.023 | 0.153 | 0.008 | 0.195 | 0.012 | 0.195 | 0.005 |
| | | | BridgePRS | 0.134 | 0.026 | 0.095 | 0.019 | 0.146 | 0.014 | 0.142 | 0.012 |
| | | 2,000 | JointPRS | 0.234 | 0.013 | 0.206 | 0.009 | 0.237 | 0.005 | 0.239 | 0.006 |
| | | | XPASS | 0.096 | 0.007 | 0.065 | 0.004 | 0.112 | 0.010 | 0.037 | 0.004 |
| | | | SDPRX | 0.206 | 0.013 | 0.173 | 0.014 | N/A | N/A | N/A | N/A |
| | | | PRS-CSx | 0.221 | 0.012 | 0.195 | 0.011 | 0.224 | 0.008 | 0.228 | 0.005 |
| | | | MUSSEL | 0.205 | 0.021 | 0.201 | 0.008 | 0.186 | 0.008 | 0.227 | 0.014 |
| | | | PROSPER | 0.201 | 0.013 | 0.165 | 0.006 | 0.207 | 0.007 | 0.207 | 0.005 |
| | | | BridgePRS | 0.138 | 0.016 | 0.108 | 0.004 | 0.152 | 0.007 | 0.158 | 0.006 |
| | | 5,000 | JointPRS | 0.237 | 0.013 | 0.208 | 0.012 | 0.240 | 0.005 | 0.242 | 0.006 |
| | | | XPASS | 0.098 | 0.008 | 0.063 | 0.006 | 0.116 | 0.008 | 0.041 | 0.003 |
| | | | SDPRX | 0.207 | 0.011 | 0.173 | 0.011 | N/A | N/A | N/A | N/A |
| | | | PRS-CSx | 0.223 | 0.011 | 0.195 | 0.009 | 0.225 | 0.007 | 0.227 | 0.004 |
| | | | MUSSEL | 0.206 | 0.009 | 0.193 | 0.022 | 0.187 | 0.006 | 0.199 | 0.005 |
| | | | PROSPER | 0.208 | 0.015 | 0.176 | 0.013 | 0.213 | 0.005 | 0.214 | 0.007 |
| | | | BridgePRS | 0.141 | 0.016 | 0.107 | 0.004 | 0.158 | 0.006 | 0.159 | 0.006 |
| | | 10,000 | JointPRS | 0.238 | 0.012 | 0.209 | 0.011 | 0.239 | 0.005 | 0.240 | 0.005 |
| | | | XPASS | 0.102 | 0.009 | 0.066 | 0.008 | 0.120 | 0.008 | 0.039 | 0.006 |
| | | | SDPRX | 0.212 | 0.013 | 0.173 | 0.009 | N/A | N/A | N/A | N/A |
| | | | PRS-CSx | 0.223 | 0.011 | 0.195 | 0.010 | 0.226 | 0.006 | 0.228 | 0.004 |
| | | | MUSSEL | 0.202 | 0.013 | 0.201 | 0.009 | 0.190 | 0.008 | 0.214 | 0.018 |
| | | | PROSPER | 0.211 | 0.015 | 0.181 | 0.009 | 0.218 | 0.007 | 0.218 | 0.006 |
| | | | BridgePRS | 0.142 | 0.015 | 0.105 | 0.006 | 0.159 | 0.008 | 0.160 | 0.006 |
| | 15,000 | 500 | JointPRS | 0.151 | 0.002 | 0.113 | 0.013 | 0.157 | 0.010 | 0.155 | 0.011 |
| | | | XPASS | 0.058 | 0.005 | 0.039 | 0.002 | 0.065 | 0.007 | 0.057 | 0.006 |
| | | | SDPRX | 0.147 | 0.009 | 0.117 | 0.009 | N/A | N/A | N/A | N/A |
| | | | PRS-CSx | 0.149 | 0.009 | 0.112 | 0.011 | 0.150 | 0.011 | 0.153 | 0.009 |
| | | | MUSSEL | 0.139 | 0.011 | 0.122 | 0.012 | 0.128 | 0.011 | 0.149 | 0.014 |
| | | | PROSPER | 0.117 | 0.008 | 0.069 | 0.025 | 0.126 | 0.010 | 0.129 | 0.010 |
| | | | BridgePRS | 0.094 | 0.007 | 0.064 | 0.006 | 0.109 | 0.009 | 0.113 | 0.004 |
| | | 2,000 | JointPRS | 0.159 | 0.009 | 0.124 | 0.010 | 0.166 | 0.003 | 0.167 | 0.007 |
| | | | XPASS | 0.058 | 0.005 | 0.039 | 0.002 | 0.065 | 0.007 | 0.057 | 0.006 |
| | | | SDPRX | 0.147 | 0.009 | 0.117 | 0.009 | N/A | N/A | N/A | N/A |
| | | | PRS-CSx | 0.152 | 0.008 | 0.117 | 0.008 | 0.158 | 0.003 | 0.158 | 0.007 |
| | | | MUSSEL | 0.143 | 0.008 | 0.124 | 0.010 | 0.141 | 0.009 | 0.152 | 0.004 |
| | | | PROSPER | 0.127 | 0.009 | 0.087 | 0.013 | 0.131 | 0.009 | 0.136 | 0.005 |
| | | | BridgePRS | 0.095 | 0.009 | 0.067 | 0.006 | 0.110 | 0.009 | 0.116 | 0.004 |
| | | 5,000 | JointPRS | 0.159 | 0.009 | 0.125 | 0.010 | 0.168 | 0.004 | 0.169 | 0.006 |
| | | | XPASS | 0.062 | 0.005 | 0.041 | 0.003 | 0.069 | 0.007 | 0.057 | 0.006 |
| | | | SDPRX | 0.154 | 0.011 | 0.124 | 0.010 | N/A | N/A | N/A | N/A |
| | | | PRS-CSx | 0.153 | 0.009 | 0.118 | 0.008 | 0.160 | 0.004 | 0.162 | 0.005 |
| | | | MUSSEL | 0.142 | 0.009 | 0.124 | 0.009 | 0.139 | 0.008 | 0.151 | 0.005 |
| | | | PROSPER | 0.115 | 0.026 | 0.085 | 0.015 | 0.132 | 0.013 | 0.119 | 0.046 |
| | | | BridgePRS | 0.098 | 0.011 | 0.069 | 0.006 | 0.113 | 0.005 | 0.117 | 0.002 |
| | | 10,000 | JointPRS | 0.160 | 0.009 | 0.125 | 0.010 | 0.168 | 0.004 | 0.169 | 0.007 |
| | | | XPASS | 0.065 | 0.006 | 0.043 | 0.002 | 0.074 | 0.006 | 0.055 | 0.009 |
| | | | SDPRX | 0.161 | 0.012 | 0.132 | 0.011 | N/A | N/A | N/A | N/A |
| | | | PRS-CSx | 0.154 | 0.009 | 0.119 | 0.009 | 0.160 | 0.004 | 0.163 | 0.005 |
| | | | MUSSEL | 0.145 | 0.012 | 0.125 | 0.010 | 0.143 | 0.009 | 0.153 | 0.010 |
| | | | PROSPER | 0.126 | 0.016 | 0.079 | 0.026 | 0.135 | 0.014 | 0.119 | 0.047 |
| | | | BridgePRS | 0.101 | 0.009 | 0.070 | 0.004 | 0.115 | 0.007 | 0.119 | 0.001 |

**Table S4: Simulation Results for the Mean and Standard Deviation $R^2$ Values across Five Replications When a Tuning Dataset is Available in EAS, AFR, SAS, and AMR Populations for Causal SNP Proportions $p = 0.001$.**

| p | minor_sample | minor_val_sample | method | EAS Mean | EAS SD | AFR Mean | AFR SD | SAS Mean | SAS SD | AMR Mean | AMR SD |
|---|---|---|---|---|---|---|---|---|---|---|---|
| 0.001 | 80,000 | 500 | JointPRS | 0.306 | 0.013 | 0.287 | 0.011 | 0.305 | 0.016 | 0.302 | 0.006 |
| | | | XPASS | 0.227 | 0.018 | 0.217 | 0.011 | 0.224 | 0.010 | 0.224 | 0.005 |
| | | | SDPRX | 0.310 | 0.014 | 0.296 | 0.011 | N/A | N/A | N/A | N/A |
| | | | PRS-CSx | 0.307 | 0.017 | 0.288 | 0.007 | 0.307 | 0.013 | 0.302 | 0.010 |
| | | | MUSSEL | 0.260 | 0.049 | 0.286 | 0.051 | 0.198 | 0.037 | 0.256 | 0.032 |
| | | | PROSPER | 0.283 | 0.012 | 0.263 | 0.011 | 0.283 | 0.017 | 0.282 | 0.011 |
| | | | BridgePRS | 0.230 | 0.008 | 0.201 | 0.023 | 0.247 | 0.009 | 0.232 | 0.009 |
| | | 2,000 | JointPRS | 0.313 | 0.015 | 0.293 | 0.012 | 0.315 | 0.008 | 0.308 | 0.007 |
| | | | XPASS | 0.227 | 0.018 | 0.217 | 0.011 | 0.224 | 0.010 | 0.224 | 0.005 |
| | | | SDPRX | 0.310 | 0.014 | 0.296 | 0.011 | N/A | N/A | N/A | N/A |
| | | | PRS-CSx | 0.313 | 0.016 | 0.293 | 0.009 | 0.313 | 0.007 | 0.308 | 0.005 |
| | | | MUSSEL | 0.264 | 0.025 | 0.306 | 0.015 | 0.160 | 0.092 | 0.246 | 0.011 |
| | | | PROSPER | 0.294 | 0.011 | 0.272 | 0.007 | 0.298 | 0.009 | 0.294 | 0.006 |
| | | | BridgePRS | 0.241 | 0.017 | 0.204 | 0.022 | 0.254 | 0.010 | 0.232 | 0.008 |
| | | 5,000 | JointPRS | 0.313 | 0.016 | 0.295 | 0.011 | 0.315 | 0.007 | 0.310 | 0.007 |
| | | | XPASS | 0.234 | 0.021 | 0.224 | 0.013 | 0.227 | 0.007 | 0.232 | 0.006 |
| | | | SDPRX | 0.313 | 0.013 | 0.297 | 0.014 | N/A | N/A | N/A | N/A |
| | | | PRS-CSx | 0.313 | 0.016 | 0.295 | 0.008 | 0.315 | 0.005 | 0.309 | 0.006 |
| | | | MUSSEL | 0.284 | 0.043 | 0.308 | 0.016 | 0.171 | 0.086 | 0.262 | 0.038 |
| | | | PROSPER | 0.298 | 0.013 | 0.278 | 0.006 | 0.304 | 0.006 | 0.298 | 0.006 |
| | | | BridgePRS | 0.241 | 0.017 | 0.206 | 0.013 | 0.251 | 0.009 | 0.234 | 0.007 |
| | | 10,000 | JointPRS | 0.313 | 0.016 | 0.295 | 0.011 | 0.316 | 0.007 | 0.310 | 0.007 |
| | | | XPASS | 0.238 | 0.022 | 0.230 | 0.011 | 0.235 | 0.008 | 0.236 | 0.005 |
| | | | SDPRX | 0.313 | 0.016 | 0.299 | 0.012 | N/A | N/A | N/A | N/A |
| | | | PRS-CSx | 0.313 | 0.016 | 0.295 | 0.009 | 0.316 | 0.005 | 0.309 | 0.006 |
| | | | MUSSEL | 0.283 | 0.045 | 0.308 | 0.015 | 0.208 | 0.017 | 0.253 | 0.011 |
| | | | PROSPER | 0.302 | 0.012 | 0.280 | 0.007 | 0.307 | 0.006 | 0.302 | 0.007 |
| | | | BridgePRS | 0.241 | 0.016 | 0.206 | 0.011 | 0.251 | 0.009 | 0.233 | 0.005 |
| | 15,000 | 500 | JointPRS | 0.250 | 0.012 | 0.221 | 0.012 | 0.253 | 0.009 | 0.248 | 0.009 |
| | | | XPASS | 0.026 | 0.010 | 0.023 | 0.008 | 0.029 | 0.004 | 0.025 | 0.006 |
| | | | SDPRX | 0.250 | 0.013 | 0.225 | 0.012 | N/A | N/A | N/A | N/A |
| | | | PRS-CSx | 0.250 | 0.015 | 0.212 | 0.010 | 0.257 | 0.007 | 0.253 | 0.013 |
| | | | MUSSEL | 0.190 | 0.026 | 0.222 | 0.014 | 0.172 | 0.069 | 0.219 | 0.016 |
| | | | PROSPER | 0.201 | 0.019 | 0.169 | 0.010 | 0.206 | 0.023 | 0.203 | 0.013 |
| | | | BridgePRS | 0.120 | 0.018 | 0.082 | 0.025 | 0.159 | 0.018 | 0.153 | 0.008 |
| | | 2,000 | JointPRS | 0.258 | 0.014 | 0.221 | 0.012 | 0.263 | 0.009 | 0.257 | 0.011 |
| | | | XPASS | 0.026 | 0.010 | 0.023 | 0.008 | 0.029 | 0.004 | 0.025 | 0.006 |
| | | | SDPRX | 0.250 | 0.013 | 0.225 | 0.012 | N/A | N/A | N/A | N/A |
| | | | PRS-CSx | 0.258 | 0.012 | 0.228 | 0.016 | 0.264 | 0.006 | 0.258 | 0.011 |
| | | | MUSSEL | 0.198 | 0.069 | 0.229 | 0.012 | 0.163 | 0.080 | 0.222 | 0.027 |
| | | | PROSPER | 0.213 | 0.011 | 0.190 | 0.012 | 0.223 | 0.015 | 0.221 | 0.010 |
| | | | BridgePRS | 0.136 | 0.004 | 0.108 | 0.013 | 0.167 | 0.011 | 0.162 | 0.004 |
| | | 5,000 | JointPRS | 0.261 | 0.012 | 0.233 | 0.009 | 0.265 | 0.007 | 0.261 | 0.007 |
| | | | XPASS | 0.053 | 0.010 | 0.048 | 0.011 | 0.050 | 0.006 | 0.042 | 0.005 |
| | | | SDPRX | 0.263 | 0.006 | 0.241 | 0.014 | N/A | N/A | N/A | N/A |
| | | | PRS-CSx | 0.259 | 0.012 | 0.230 | 0.012 | 0.264 | 0.006 | 0.260 | 0.007 |
| | | | MUSSEL | 0.202 | 0.014 | 0.230 | 0.014 | 0.165 | 0.086 | 0.243 | 0.030 |
| | | | PROSPER | 0.222 | 0.012 | 0.195 | 0.009 | 0.231 | 0.011 | 0.226 | 0.009 |
| | | | BridgePRS | 0.136 | 0.008 | 0.106 | 0.012 | 0.168 | 0.009 | 0.165 | 0.005 |
| | | 10,000 | JointPRS | 0.261 | 0.012 | 0.232 | 0.011 | 0.265 | 0.007 | 0.261 | 0.006 |
| | | | XPASS | 0.075 | 0.012 | 0.074 | 0.009 | 0.076 | 0.011 | 0.074 | 0.006 |
| | | | SDPRX | 0.272 | 0.011 | 0.250 | 0.011 | N/A | N/A | N/A | N/A |
| | | | PRS-CSx | 0.259 | 0.013 | 0.231 | 0.013 | 0.265 | 0.006 | 0.260 | 0.007 |
| | | | MUSSEL | 0.231 | 0.027 | 0.229 | 0.015 | 0.175 | 0.056 | 0.216 | 0.011 |
| | | | PROSPER | 0.227 | 0.012 | 0.199 | 0.010 | 0.235 | 0.013 | 0.231 | 0.011 |
| | | | BridgePRS | 0.139 | 0.005 | 0.106 | 0.011 | 0.168 | 0.010 | 0.166 | 0.006 |

**Table S5: Simulation Results for the Mean and Standard Deviation $R^2$ Values across Five Replications When a Tuning Dataset is Available in EAS, AFR, SAS, and AMR Populations for Causal SNP Proportions $p = 5 \times 10^{-4}$.**

| p | minor_sample | minor_val_sample | method | EAS | | AFR | | SAS | | AMR | |
|---|---|---|---|---|---|---|---|---|---|---|---|
| | | | | Mean | SD | Mean | SD | Mean | SD | Mean | SD |
| | | | JointPRS | 0.324 | 0.009 | 0.294 | 0.011 | 0.318 | 0.013 | 0.320 | 0.004 |
| | | | XPASS | 0.278 | 0.011 | 0.264 | 0.010 | 0.265 | 0.014 | 0.283 | 0.007 |
| | | | SDPRX | 0.329 | 0.012 | 0.305 | 0.008 | N/A | N/A | N/A | N/A |
| | | 500 | PRS-CSx | 0.322 | 0.007 | 0.293 | 0.009 | 0.316 | 0.011 | 0.316 | 0.012 |
| | | | MUSSEL | 0.262 | 0.049 | 0.313 | 0.009 | 0.206 | 0.016 | 0.239 | 0.015 |
| | | | PROSPER | 0.296 | 0.020 | 0.265 | 0.018 | 0.289 | 0.029 | 0.299 | 0.010 |
| | | | BridgePRS | 0.268 | 0.011 | 0.234 | 0.017 | 0.276 | 0.013 | 0.261 | 0.007 |
| | | | JointPRS | 0.329 | 0.010 | 0.296 | 0.009 | 0.325 | 0.013 | 0.324 | 0.004 |
| | | | XPASS | 0.278 | 0.011 | 0.264 | 0.010 | 0.265 | 0.014 | 0.283 | 0.007 |
| | | | SDPRX | 0.329 | 0.012 | 0.305 | 0.008 | N/A | N/A | N/A | N/A |
| | | 2,000 | PRS-CSx | 0.329 | 0.007 | 0.300 | 0.013 | 0.324 | 0.011 | 0.323 | 0.004 |
| | | | MUSSEL | 0.274 | 0.042 | 0.293 | 0.064 | 0.216 | 0.006 | 0.262 | 0.039 |
| | | | PROSPER | 0.310 | 0.010 | 0.286 | 0.014 | 0.310 | 0.012 | 0.309 | 0.007 |
| | | | BridgePRS | 0.269 | 0.009 | 0.245 | 0.013 | 0.279 | 0.016 | 0.264 | 0.007 |
| | 80,000 | | JointPRS | 0.331 | 0.008 | 0.299 | 0.011 | 0.326 | 0.013 | 0.325 | 0.004 |
| | | | XPASS | 0.284 | 0.009 | 0.267 | 0.009 | 0.268 | 0.011 | 0.289 | 0.010 |
| | | | SDPRX | 0.332 | 0.009 | 0.306 | 0.007 | N/A | N/A | N/A | N/A |
| | | 5,000 | PRS-CSx | 0.330 | 0.007 | 0.301 | 0.013 | 0.325 | 0.011 | 0.324 | 0.004 |
| | | | MUSSEL | 0.258 | 0.051 | 0.320 | 0.010 | 0.200 | 0.019 | 0.248 | 0.011 |
| | | | PROSPER | 0.316 | 0.008 | 0.290 | 0.011 | 0.314 | 0.011 | 0.314 | 0.007 |
| | | | BridgePRS | 0.266 | 0.009 | 0.241 | 0.007 | 0.277 | 0.014 | 0.263 | 0.006 |
| | | | JointPRS | 0.330 | 0.009 | 0.298 | 0.011 | 0.326 | 0.013 | 0.325 | 0.004 |
| | | | XPASS | 0.286 | 0.011 | 0.271 | 0.007 | 0.269 | 0.012 | 0.291 | 0.009 |
| | | | SDPRX | 0.332 | 0.008 | 0.308 | 0.008 | N/A | N/A | N/A | N/A |
| | | 10,000 | PRS-CSx | 0.331 | 0.007 | 0.301 | 0.013 | 0.325 | 0.011 | 0.325 | 0.004 |
| | | | MUSSEL | 0.258 | 0.006 | 0.319 | 0.010 | 0.175 | 0.097 | 0.248 | 0.011 |
| | | | PROSPER | 0.318 | 0.008 | 0.294 | 0.011 | 0.317 | 0.013 | 0.317 | 0.006 |
| | | | BridgePRS | 0.267 | 0.013 | 0.241 | 0.010 | 0.277 | 0.012 | 0.265 | 0.007 |
| $5 \times 10^{-4}$ | | | JointPRS | 0.273 | 0.009 | 0.239 | 0.011 | 0.275 | 0.011 | 0.275 | 0.010 |
| | | | XPASS | 0.083 | 0.011 | 0.075 | 0.004 | 0.082 | 0.009 | 0.087 | 0.009 |
| | | | SDPRX | 0.278 | 0.012 | 0.244 | 0.010 | N/A | N/A | N/A | N/A |
| | | 500 | PRS-CSx | 0.285 | 0.011 | 0.244 | 0.015 | 0.279 | 0.010 | 0.280 | 0.012 |
| | | | MUSSEL | 0.218 | 0.003 | 0.247 | 0.011 | 0.205 | 0.014 | 0.220 | 0.015 |
| | | | PROSPER | 0.212 | 0.026 | 0.175 | 0.035 | 0.219 | 0.020 | 0.216 | 0.031 |
| | | | BridgePRS | 0.183 | 0.013 | 0.126 | 0.038 | 0.189 | 0.010 | 0.173 | 0.015 |
| | | | JointPRS | 0.285 | 0.011 | 0.241 | 0.010 | 0.279 | 0.009 | 0.279 | 0.007 |
| | | | XPASS | 0.083 | 0.011 | 0.075 | 0.004 | 0.082 | 0.009 | 0.087 | 0.009 |
| | | | SDPRX | 0.278 | 0.012 | 0.244 | 0.010 | N/A | N/A | N/A | N/A |
| | | 2,000 | PRS-CSx | 0.288 | 0.011 | 0.252 | 0.014 | 0.285 | 0.008 | 0.287 | 0.006 |
| | | | MUSSEL | 0.236 | 0.034 | 0.225 | 0.048 | 0.164 | 0.092 | 0.252 | 0.024 |
| | | | PROSPER | 0.240 | 0.018 | 0.217 | 0.018 | 0.244 | 0.015 | 0.251 | 0.006 |
| | | | BridgePRS | 0.176 | 0.011 | 0.143 | 0.022 | 0.195 | 0.015 | 0.190 | 0.012 |
| | 15,000 | | JointPRS | 0.288 | 0.009 | 0.253 | 0.011 | 0.287 | 0.009 | 0.288 | 0.005 |
| | | | XPASS | 0.122 | 0.008 | 0.112 | 0.016 | 0.119 | 0.006 | 0.132 | 0.012 |
| | | | SDPRX | 0.287 | 0.009 | 0.260 | 0.007 | N/A | N/A | N/A | N/A |
| | | 5,000 | PRS-CSx | 0.288 | 0.011 | 0.253 | 0.013 | 0.285 | 0.008 | 0.288 | 0.007 |
| | | | MUSSEL | 0.255 | 0.040 | 0.259 | 0.012 | 0.205 | 0.012 | 0.227 | 0.016 |
| | | | PROSPER | 0.256 | 0.015 | 0.226 | 0.013 | 0.254 | 0.013 | 0.258 | 0.007 |
| | | | BridgePRS | 0.176 | 0.011 | 0.142 | 0.018 | 0.192 | 0.011 | 0.194 | 0.012 |
| | | | JointPRS | 0.289 | 0.009 | 0.258 | 0.008 | 0.287 | 0.009 | 0.289 | 0.005 |
| | | | XPASS | 0.153 | 0.010 | 0.136 | 0.019 | 0.149 | 0.011 | 0.159 | 0.011 |
| | | | SDPRX | 0.297 | 0.009 | 0.266 | 0.005 | N/A | N/A | N/A | N/A |
| | | 10,000 | PRS-CSx | 0.289 | 0.011 | 0.253 | 0.013 | 0.285 | 0.008 | 0.288 | 0.007 |
| | | | MUSSEL | 0.227 | 0.008 | 0.259 | 0.011 | 0.208 | 0.013 | 0.226 | 0.021 |
| | | | PROSPER | 0.261 | 0.014 | 0.228 | 0.013 | 0.257 | 0.013 | 0.263 | 0.008 |
| | | | BridgePRS | 0.176 | 0.013 | 0.140 | 0.016 | 0.192 | 0.011 | 0.197 | 0.012 |

#### Table S6: GWAS Summary Statistics SNP Number Information for 26 Traits across Five Populations.

| | | GWAS SNP number | | | | |
|---|---|---|---|---|---|---|
| **GLGC traits** | | | | | | |
| Trait | | **EUR** | **EAS** | **AFR** | **SAS** | **AMR** |
| HDL | HDL-cholesterol | 800,281 | 735,249 | 827,727 | 1,085,452 | 1,107,923 |
| LDL | LDL-cholesterol | 800,283 | 797,861 | 827,727 | 1,088,264 | 1,104,517 |
| TC | Total cholesterol | 800,281 | 461,893 | 827,727 | 1,085,270 | 1,111,066 |
| logTG | Triglycerides | 800,286 | 797,898 | 827,727 | 1,088,215 | 1,109,126 |
| **PAGE traits** | | | | | | |
| Trait | | **EUR** | **EAS** | **AFR** | **SAS** | **AMR** |
| Height | Height | 724,431 | 790,675 | 827,738 | N/A | N/A |
| BMI | Body mass index | 725,221 | 782,322 | 827,738 | N/A | N/A |
| SBP | Systolic blood pressure | 797,661 | 747,306 | 827,738 | N/A | N/A |
| DBP | Diastolic blood pressure | 798,292 | 747,306 | 827,738 | N/A | N/A |
| PLT | Platelet | 800,321 | 747,306 | 827,738 | N/A | N/A |
| **BBJ traits** | | | | | | |
| Trait | | **EUR** | **EAS** | **AFR** | **SAS** | **AMR** |
| WBC | White blood cell | 800,320 | 747,306 | N/A | N/A | N/A |
| NEU | Neutrophil | 800,319 | 747,306 | N/A | N/A | N/A |
| LYM | Lymphocyte | 800,320 | 747,306 | N/A | N/A | N/A |
| MON | Monocyte | 800,320 | 747,306 | N/A | N/A | N/A |
| EOS | Eosinophil | 800,319 | 747,306 | N/A | N/A | N/A |
| RBC | Red blood cell | 800,318 | 747,306 | N/A | N/A | N/A |
| HCT | Hematocrit | 800,319 | 747,306 | N/A | N/A | N/A |
| MCH | Mean corpuscular hemoglobin | 800,319 | 747,306 | N/A | N/A | N/A |
| MCV | Mean corpuscular volume | 800,317 | 747,306 | N/A | N/A | N/A |
| HB | Hemoglobin | 800,322 | 747,306 | N/A | N/A | N/A |
| ALT | Alanine aminotransferase | 787,866 | 747,306 | N/A | N/A | N/A |
| ALP | Alkaline phosphatase | 799,800 | 747,306 | N/A | N/A | N/A |
| GGT | $\gamma$-glutamyl transpeptidase | 799,800 | 747,306 | N/A | N/A | N/A |
| **Binary traits** | | | | | | |
| Trait | | **EUR** | **EAS** | **AFR** | | |
| T2D | Type 2 diabetes | 800,320 | 799,544 | 827,738 | | |
| BrC | Breast cancer | 800,245 | 747,306 | 837,774 | | |
| CAD | Coronary artery disease | 653,825 | 786,215 | N/A | | |
| LuC | Lung cancer | 756,305 | 786,215 | N/A | | |

**Table S7: Pairwise Cross-Population Genetic Correlation for 26 Traits across Five Populations Using Popcorn.**

**GLGC traits**

| Trait | Cross-population genetic correlation | | | | |
|---|---|---|---|---|---|
| | **EUR_EAS** | **EUR_AFR** | **EUR_SAS** | **EUR_AMR** | **EAS_AFR** |
| HDL | 0.99 | 0.93 | 0.99 | 0.95 | 0.71 |
| LDL | 0.90 | 0.70 | 0.99 | 0.92 | 0.61 |
| TC | 0.99 | 0.74 | 0.99 | 0.92 | 0.60 |
| logTG | 0.99 | 0.93 | 0.91 | 0.96 | 0.92 |

| Trait | Cross-population genetic correlation | | | | |
|---|---|---|---|---|---|
| | **EAS_SAS** | **EAS_AMR** | **AFR_SAS** | **AFR_AMR** | **SAS_AMR** |
| HDL | 0.99 | 0.85 | 0.99 | 0.96 | 0.99 |
| LDL | 0.91 | 0.83 | 0.53 | 0.63 | 0.85 |
| TC | 0.99 | 0.82 | 0.58 | 0.67 | 0.86 |
| logTG | 0.99 | 0.96 | 0.78 | 0.99 | 0.96 |

**PAGE traits**

| Trait | Cross-population genetic correlation | | | | |
|---|---|---|---|---|---|
| | **EUR_EAS** | **EUR_AFR** | **EUR_SAS** | **EUR_AMR** | **EAS_AFR** |
| Height | 0.87 | 0.99 | N/A | N/A | 0.97 |
| BMI | 0.84 | 0.99 | N/A | N/A | 0.92 |
| SBP | 0.83 | 0.99 | N/A | N/A | 0.93 |
| DBP | 0.72 | 0.99 | N/A | N/A | 0.99 |
| PLT | 0.68 | 0.99 | N/A | N/A | 0.99 |

**BBJ traits**

| Trait | Cross-population genetic correlation | | | | |
|---|---|---|---|---|---|
| | **EUR_EAS** | **EUR_AFR** | **EUR_SAS** | **EUR_AMR** | **EAS_AFR** |
| WBC | 0.77 | N/A | N/A | N/A | N/A |
| NEU | 0.80 | N/A | N/A | N/A | N/A |
| LYM | 0.78 | N/A | N/A | N/A | N/A |
| MON | 0.80 | N/A | N/A | N/A | N/A |
| EOS | 0.86 | N/A | N/A | N/A | N/A |
| RBC | 0.92 | N/A | N/A | N/A | N/A |
| HCT | 0.85 | N/A | N/A | N/A | N/A |
| MCH | 0.87 | N/A | N/A | N/A | N/A |
| MCV | 0.89 | N/A | N/A | N/A | N/A |
| HB | 0.84 | N/A | N/A | N/A | N/A |
| ALT | 0.69 | N/A | N/A | N/A | N/A |
| ALP | 0.52 | N/A | N/A | N/A | N/A |
| GGT | 0.79 | N/A | N/A | N/A | N/A |

**Binary traits**

| Trait | Cross-population genetic correlation | | | | |
|---|---|---|---|---|---|
| | **EUR_EAS** | **EUR_AFR** | **EUR_SAS** | **EUR_AMR** | **EAS_AFR** |
| T2D | 0.87 | 0.99 | N/A | N/A | 0.94 |
| BrC | 0.88 | 0.17 | N/A | N/A | 0.15 |
| CAD | 0.79 | N/A | N/A | N/A | N/A |
| LuC | 0.99 | N/A | N/A | N/A | N/A |

#### Table S8: UKBB Sample Sizes Information for 26 Traits across Five Populations.

**GLGC traits**

| Trait | | UKBB sample size | | | | UKBB Field |
|---|---|---|---|---|---|---|
| | | EAS | AFR | SAS | AMR | |
| HDL | HDL-cholesterol | 1,812 | 5,927 | 6,784 | 561 | 30760 |
| LDL | LDL-cholesterol | 1,911 | 6,171 | 7,062 | 583 | 30780 |
| TC | Total cholesterol | 1,912 | 6,183 | 7,080 | 583 | 30690 |
| logTG | Triglycerides | 1,993 | 6,400 | 7,444 | 607 | 30870 |

**PAGE traits**

| Trait | | UKBB sample size | | | | UKBB Field |
|---|---|---|---|---|---|---|
| | | EAS | AFR | SAS | AMR | |
| Height | Height | 2,080 | 6,574 | N/A | N/A | 50 |
| BMI | Body mass index | 2,077 | 6,713 | N/A | N/A | 21001 |
| SBP | Systolic blood pressure | 2,001 | 6,688 | N/A | N/A | 93; 4080 |
| DBP | Diastolic blood pressure | 2,001 | 6,574 | N/A | N/A | 94; 4090 |
| PLT | Platelet | 2,027 | 6,445 | N/A | N/A | 30080 |

**BBJ traits**

| Trait | | UKBB sample size | | | | UKBB Field |
|---|---|---|---|---|---|---|
| | | EAS | AFR | SAS | AMR | |
| WBC | White blood cell | 2,027 | N/A | N/A | N/A | 30000 |
| NEU | Neutrophil | 2,025 | N/A | N/A | N/A | 30140 |
| LYM | Lymphocyte | 2,025 | N/A | N/A | N/A | 30120 |
| MON | Monocyte | 2,025 | N/A | N/A | N/A | 30130 |
| EOS | Eosinophil | 2,025 | N/A | N/A | N/A | 30150 |
| RBC | Red blood cell | 2,027 | N/A | N/A | N/A | 30010 |
| HCT | Hematocrit | 2,027 | N/A | N/A | N/A | 30030 |
| MCH | Mean corpuscular hemoglobin | 2,027 | N/A | N/A | N/A | 30050 |
| MCV | Mean corpuscular volume | 2,027 | N/A | N/A | N/A | 30040 |
| HB | Hemoglobin | 2,027 | N/A | N/A | N/A | 30020 |
| ALT | Alanine aminotransferase | 1,993 | N/A | N/A | N/A | 30620 |
| ALP | Alkaline phosphatase | 1,994 | N/A | N/A | N/A | 30610 |
| GGT | $\gamma$-glutamyl transpeptidase | 1,991 | N/A | N/A | N/A | 30730 |

**Binary traits**

| Trait | | UKBB sample size | | | |
|---|---|---|---|---|---|
| | | EAS | | AFR | |
| | | case | control | case | control |
| T2D | Type 2 diabetes | 203 | 1,887 | 1,292 | 5,535 |
| BrC | Breast cancer | 89 | 2,001 | 173 | 6,654 |
| CAD | Coronary artery disease | 56 | 2,034 | N/A | N/A |
| LuC | Lung cancer | 21 | 2,069 | N/A | N/A |

#### Table S9: AoU Sample Sizes Information for Nine Traits across Two Populations.

| **GLGC traits** | | | | |
|---|---|---|---|---|
| **Trait** | | **AoU sample size** | | **AoU Concept Id** |
| | | **AFR** | **AMR** | |
| HDL | HDL-cholesterol | 12822 | 7004 | 3007070 |
| LDL | LDL-cholesterol | 9359 | 5457 | 3028288 |
| TC | Total cholesterol | 12429 | 7016 | 3027114 |
| logTG | Triglycerides | 12395 | 7020 | 3022192 |
| **PAGE traits** | | | | |
| **Trait** | | **AoU sample size** | | **AoU Concept Id** |
| | | **AFR** | **AMR** | |
| Height | Height | 51790 | N/A | 903133 |
| BMI | Body mass index | 50747 | N/A | 903124 |
| SBP | Systolic blood pressure | 51342 | N/A | 903118 |
| DBP | Diastolic blood pressure | 51395 | N/A | 903115 |
| PLT | Platelet | 24517 | N/A | 37037425 |

**Table S10: Real Data Results for $R^2$ of Nine Traits in UKBB When a Tuning Dataset is Not Available in EAS, AFR, SAS, and AMR.**

| GLGC traits | | | | | |
|---|---|---|---|---|---|
| Trait | pop | R2 value | | | |
| | | JointPRS-auto | XPASS | SDPRX | PRS-CSx-auto |
| HDL | EAS | 0.168 | 0.131 | 0.143 | 0.144 |
| | AFR | 0.090 | 0.071 | 0.087 | 0.077 |
| | SAS | 0.133 | 0.100 | N/A | 0.109 |
| | AMR | 0.140 | 0.063 | N/A | 0.125 |
| LDL | EAS | 0.087 | 0.074 | 0.091 | 0.082 |
| | AFR | 0.106 | 0.066 | 0.119 | 0.102 |
| | SAS | 0.077 | 0.030 | N/A | 0.061 |
| | AMR | 0.081 | 0.057 | N/A | 0.065 |
| TC | EAS | 0.088 | 0.068 | 0.092 | 0.076 |
| | AFR | 0.094 | 0.068 | 0.110 | 0.088 |
| | SAS | 0.075 | 0.041 | N/A | 0.060 |
| | AMR | 0.085 | 0.044 | N/A | 0.073 |
| logTG | EAS | 0.093 | 0.072 | 0.098 | 0.078 |
| | AFR | 0.045 | 0.034 | 0.044 | 0.036 |
| | SAS | 0.108 | 0.086 | N/A | 0.091 |
| | AMR | 0.078 | 0.040 | N/A | 0.060 |
| **PAGE traits** | | | | | |
| Trait | pop | R2 value | | | |
| | | JointPRS-auto | XPASS | SDPRX | PRS-CSx-auto |
| Height | EAS | 0.234 | 0.175 | 0.217 | 0.220 |
| | AFR | 0.091 | 0.053 | 0.077 | 0.062 |
| BMI | EAS | 0.075 | 0.060 | 0.081 | 0.059 |
| | AFR | 0.039 | 0.028 | 0.034 | 0.028 |
| SBP | EAS | 0.072 | 0.045 | 0.077 | 0.055 |
| | AFR | 0.016 | 0.009 | 0.024 | 0.009 |
| DBP | EAS | 0.073 | 0.046 | 0.075 | 0.055 |
| | AFR | 0.014 | 0.007 | 0.019 | 0.009 |
| PLT | EAS | 0.128 | 0.095 | 0.090 | 0.106 |
| | AFR | 0.047 | 0.021 | 0.014 | 0.038 |

**Table S11: Real Data Results for $R^2$ or AUC of 17 Traits in UKBB When a Tuning Dataset is Not Available in EAS and AFR.**

| BBJ traits | | | | | |
|---|---|---|---|---|---|
| | | R2 value | | | |
| **Trait** | **pop** | **JointPRS-auto** | **XPASS** | **SDPRX** | **PRS-CSx-auto** |
| WBC | EAS | 0.075 | 0.040 | 0.073 | 0.056 |
| NEU | EAS | 0.050 | 0.032 | 0.053 | 0.036 |
| LYM | EAS | 0.091 | 0.049 | 0.092 | 0.073 |
| MON | EAS | 0.077 | 0.047 | 0.080 | 0.069 |
| EOS | EAS | 0.036 | 0.019 | 0.035 | 0.029 |
| RBC | EAS | 0.078 | 0.062 | 0.087 | 0.065 |
| HB | EAS | 0.046 | 0.021 | 0.050 | 0.040 |
| HCT | EAS | 0.059 | 0.026 | 0.058 | 0.044 |
| MCH | EAS | 0.055 | 0.051 | 0.052 | 0.051 |
| MCV | EAS | 0.064 | 0.048 | 0.056 | 0.058 |
| ALT | EAS | 0.033 | 0.021 | 0.033 | 0.028 |
| ALP | EAS | 0.091 | 0.060 | 0.075 | 0.079 |
| GGT | EAS | 0.030 | 0.018 | 0.030 | 0.023 |
| **Binary traits** | | | | | |
| | | **AUC value** | | | |
| **Trait** | **pop** | **JointPRS-auto** | **XPASS** | **SDPRX** | **PRS-CSx-auto** |
| T2D | EAS | 0.619 | 0.599 | 0.549 | 0.614 |
| | AFR | 0.567 | 0.543 | 0.555 | 0.554 |
| BrC | EAS | 0.644 | 0.613 | 0.637 | 0.612 |
| | AFR | 0.551 | 0.551 | 0.532 | 0.530 |
| CAD | EAS | 0.577 | 0.558 | 0.602 | 0.564 |
| LuC | EAS | 0.558 | 0.517 | 0.594 | 0.555 |

**Table S12: Real Data Results for the Mean of $R^2$ across Five Folds of Nine Traits When the Tuning and Testing Data Come From the Same Cohort (UKBB) in EAS, AFR, SAS, and AMR.**

| GLGC traits | | | | | | | | |
|---|---|---|---|---|---|---|---|---|
| Trait | pop | Mean of R2 value across five folds | | | | | | |
| | | **JointPRS** | **XPASS** | **SDPRX** | **PRS-CSx** | **MUSSEL** | **PROSPER** | **BridgePRS** |
| HDL | EAS | 0.168 | 0.132 | 0.146 | 0.154 | 0.147 | 0.158 | 0.134 |
| | AFR | 0.093 | 0.057 | 0.087 | 0.091 | 0.074 | 0.100 | 0.068 |
| | SAS | 0.149 | 0.084 | N/A | 0.144 | 0.130 | 0.158 | 0.129 |
| | AMR | 0.162 | 0.068 | N/A | 0.154 | 0.133 | 0.154 | 0.097 |
| LDL | EAS | 0.086 | 0.074 | 0.092 | 0.081 | 0.043 | 0.075 | 0.051 |
| | AFR | 0.110 | 0.059 | 0.124 | 0.113 | 0.078 | 0.098 | 0.073 |
| | SAS | 0.084 | 0.034 | N/A | 0.081 | 0.085 | 0.093 | 0.056 |
| | AMR | 0.089 | 0.060 | N/A | 0.056 | 0.090 | 0.073 | 0.043 |
| TC | EAS | 0.104 | 0.069 | 0.092 | 0.088 | 0.074 | 0.096 | 0.071 |
| | AFR | 0.100 | 0.052 | 0.112 | 0.098 | 0.079 | 0.100 | 0.069 |
| | SAS | 0.087 | 0.039 | N/A | 0.084 | 0.092 | 0.085 | 0.064 |
| | AMR | 0.097 | 0.066 | N/A | 0.093 | 0.098 | 0.113 | 0.072 |
| logTG | EAS | 0.095 | 0.072 | 0.098 | 0.089 | 0.097 | 0.089 | 0.080 |
| | AFR | 0.051 | 0.032 | 0.052 | 0.020 | 0.015 | 0.039 | 0.031 |
| | SAS | 0.113 | 0.080 | N/A | 0.114 | 0.101 | 0.117 | 0.096 |
| | AMR | 0.090 | 0.046 | N/A | 0.073 | 0.064 | 0.077 | 0.081 |
| **PAGE traits** | | | | | | | | |
| Trait | pop | Mean of R2 value across five folds | | | | | | |
| | | **JointPRS** | **XPASS** | **SDPRX** | **PRS-CSx** | **MUSSEL** | **PROSPER** | **BridgePRS** |
| Height | EAS | 0.236 | 0.182 | 0.224 | 0.229 | 0.197 | 0.220 | 0.128 |
| | AFR | 0.098 | 0.049 | 0.076 | 0.087 | 0.085 | 0.097 | 0.015 |
| BMI | EAS | 0.089 | 0.063 | 0.084 | 0.097 | 0.090 | 0.092 | 0.064 |
| | AFR | 0.039 | 0.028 | 0.027 | 0.041 | 0.044 | 0.043 | 0.027 |
| SBP | EAS | 0.066 | 0.045 | 0.078 | 0.066 | 0.058 | 0.064 | 0.047 |
| | AFR | 0.026 | 0.011 | 0.028 | 0.026 | 0.017 | 0.024 | 0.008 |
| DBP | EAS | 0.080 | 0.047 | 0.075 | 0.077 | 0.069 | 0.070 | 0.039 |
| | AFR | 0.020 | 0.007 | 0.020 | 0.020 | 0.014 | 0.015 | 0.006 |
| PLT | EAS | 0.136 | 0.094 | 0.091 | 0.121 | 0.099 | 0.125 | 0.003 |
| | AFR | 0.059 | 0.025 | 0.019 | 0.057 | 0.049 | 0.061 | 0.028 |

**Table S13: Real Data Results for the Mean of $R^2$ or AUC across Five Folds of 17 Traits When the Tuning and Testing Data Come From the Same Cohort (UKBB) in EAS and AFR.**

| BBJ traits | | | | | | | | |
|---|---|---|---|---|---|---|---|---|
| Trait | pop | Mean of R2 value across five folds | | | | | | |
| | | **JointPRS** | **XPASS** | **SDPRX** | **PRS-CSx** | **MUSSEL** | **PROSPER** | **BridgePRS** |
| WBC | EAS | 0.071 | 0.045 | 0.076 | 0.066 | 0.073 | 0.070 | 0.013 |
| NEU | EAS | 0.057 | 0.035 | 0.058 | 0.053 | 0.049 | 0.052 | 0.006 |
| LYM | EAS | 0.096 | 0.044 | 0.088 | 0.091 | 0.087 | 0.090 | 0.006 |
| MON | EAS | 0.077 | 0.049 | 0.083 | 0.080 | 0.086 | 0.078 | 0.018 |
| EOS | EAS | 0.039 | 0.024 | 0.038 | 0.038 | 0.043 | 0.040 | 0.014 |
| RBC | EAS | 0.088 | 0.066 | 0.093 | 0.088 | 0.080 | 0.092 | 0.006 |
| HB | EAS | 0.051 | 0.024 | 0.055 | 0.055 | 0.045 | 0.055 | 0.017 |
| HCT | EAS | 0.061 | 0.028 | 0.058 | 0.061 | 0.056 | 0.059 | 0.011 |
| MCH | EAS | 0.061 | 0.053 | 0.053 | 0.064 | 0.051 | 0.063 | 0.010 |
| MCV | EAS | 0.067 | 0.052 | 0.058 | 0.068 | 0.059 | 0.070 | 0.008 |
| ALT | EAS | 0.044 | 0.028 | 0.041 | 0.045 | 0.034 | 0.037 | 0.022 |
| ALP | EAS | 0.104 | 0.061 | 0.078 | 0.107 | 0.088 | 0.099 | 0.075 |
| GGT | EAS | 0.051 | 0.024 | 0.041 | 0.053 | 0.035 | 0.054 | 0.048 |
| **Binary traits** | | | | | | | | |
| Trait | pop | Mean of AUC value across five folds | | | | | | |
| | | **JointPRS** | **XPASS** | **SDPRX** | **PRS-CSx** | **MUSSEL** | **PROSPER** | **BridgePRS** |
| T2D | EAS | 0.613 | 0.596 | 0.547 | 0.646 | 0.628 | 0.635 | 0.583 |
| | AFR | 0.568 | 0.545 | 0.556 | 0.558 | 0.564 | 0.561 | 0.537 |
| BrC | EAS | 0.631 | 0.614 | 0.635 | 0.568 | 0.612 | 0.615 | 0.639 |
| | AFR | 0.553 | 0.555 | 0.544 | 0.586 | 0.576 | 0.575 | 0.574 |
| CAD | EAS | 0.578 | 0.576 | 0.568 | 0.587 | 0.559 | 0.564 | 0.515 |
| LuC | EAS | 0.644 | 0.575 | 0.623 | 0.659 | 0.591 | 0.710 | 0.592 |

**Table S14: JointPRS Version Selection across Five Folds of 26 Traits When the Tuning and Testing Data Come From the Same Cohort (UKBB) in Four non-European Populations.**

| | **EAS** | | **AFR** | | **SAS** | | **AMR** | |
|---|---|---|---|---|---|---|---|---|
| **Trait** | tune | meta | tune | meta | tune | meta | tune | meta |
| HDL | 0 | 5 | 5 | 0 | 5 | 0 | 3 | 2 |
| LDL | 2 | 3 | 5 | 0 | 5 | 0 | 0 | 5 |
| TC | 5 | 0 | 1 | 4 | 5 | 0 | 0 | 5 |
| logTG | 0 | 5 | 0 | 5 | 5 | 0 | 2 | 3 |
| Height | 1 | 4 | 5 | 0 | N/A | N/A | N/A | N/A |
| BMI | 3 | 2 | 5 | 0 | N/A | N/A | N/A | N/A |
| SBP | 5 | 0 | 4 | 1 | N/A | N/A | N/A | N/A |
| DBP | 5 | 0 | 5 | 0 | N/A | N/A | N/A | N/A |
| PLT | 5 | 0 | 5 | 0 | N/A | N/A | N/A | N/A |
| WBC | 3 | 2 | N/A | N/A | N/A | N/A | N/A | N/A |
| NEU | 5 | 0 | N/A | N/A | N/A | N/A | N/A | N/A |
| LYM | 5 | 0 | N/A | N/A | N/A | N/A | N/A | N/A |
| MON | 2 | 3 | N/A | N/A | N/A | N/A | N/A | N/A |
| EOS | 3 | 2 | N/A | N/A | N/A | N/A | N/A | N/A |
| RBC | 5 | 0 | N/A | N/A | N/A | N/A | N/A | N/A |
| HB | 0 | 5 | N/A | N/A | N/A | N/A | N/A | N/A |
| HCT | 0 | 5 | N/A | N/A | N/A | N/A | N/A | N/A |
| MCH | 5 | 0 | N/A | N/A | N/A | N/A | N/A | N/A |
| MCV | 4 | 1 | N/A | N/A | N/A | N/A | N/A | N/A |
| ALT | 5 | 0 | N/A | N/A | N/A | N/A | N/A | N/A |
| ALP | 5 | 0 | N/A | N/A | N/A | N/A | N/A | N/A |
| GGT | 5 | 0 | N/A | N/A | N/A | N/A | N/A | N/A |
| T2D | 0 | 5 | 0 | 5 | N/A | N/A | N/A | N/A |
| BrC | 0 | 5 | 0 | 5 | N/A | N/A | N/A | N/A |
| CAD | 0 | 5 | N/A | N/A | N/A | N/A | N/A | N/A |
| LuC | 0 | 5 | N/A | N/A | N/A | N/A | N/A | N/A |

**Table S15: Real Data Results for $R^2$ of Nine Traits When the Tuning and Testing Data Come From Different Cohorts (UKBB and AoU) in AFR and AMR.**

| GLGC traits | | | | | | | | |
|---|---|---|---|---|---|---|---|---|
| Trait | pop | R2 value | | | | | | |
| | | JointPRS | XPASS | SDPRX | PRS-CSx | MUSSEL | PROSPER | BridgePRS |
| HDL | AFR | 0.063 | 0.057 | 0.069 | 0.063 | 0.071 | 0.071 | 0.059 |
| | AMR | 0.139 | 0.059 | N/A | 0.132 | 0.137 | 0.137 | 0.112 |
| LDL | AFR | 0.059 | 0.036 | 0.065 | 0.058 | 0.051 | 0.050 | 0.046 |
| | AMR | 0.067 | 0.016 | N/A | 0.062 | 0.060 | 0.059 | 0.028 |
| TC | AFR | 0.046 | 0.032 | 0.052 | 0.049 | 0.009 | 0.050 | 0.037 |
| | AMR | 0.076 | 0.032 | N/A | 0.068 | 0.007 | 0.069 | 0.049 |
| logTG | AFR | 0.048 | 0.042 | 0.049 | 0.032 | 0.037 | 0.037 | 0.025 |
| | AMR | 0.110 | 0.060 | N/A | 0.093 | 0.099 | 0.110 | 0.092 |
| PAGE traits | | | | | | | | |
| Trait | pop | R2 value | | | | | | |
| | | JointPRS | XPASS | SDPRX | PRS-CSx | MUSSEL | PROSPER | BridgePRS |
| Height | AFR | 0.099 | 0.054 | 0.085 | 0.094 | 0.093 | 0.093 | 0.021 |
| BMI | AFR | 0.028 | 0.020 | 0.025 | 0.030 | 0.030 | 0.031 | 0.020 |
| SBP | AFR | 0.011 | 0.005 | 0.010 | 0.011 | 0.009 | 0.009 | 0.003 |
| DBP | AFR | 0.010 | 0.005 | 0.009 | 0.010 | 0.008 | 0.008 | 0.004 |
| PLT | AFR | 0.055 | 0.031 | 0.016 | 0.048 | 0.052 | 0.052 | 0.028 |

**Table S16: Computational Time and Memory Comparison in JointPRS and PRS-CSx.** We used the computational time and memory result from chromosome 2 which takes the longest computational time in JointPRS and PRS-CSx using the auto version across all quantitative traits with different number of available populations. The computational time is in hours and the computational memory usage is measured in the unit of Gigabytes (GB).

| Number of Available Populations | Trait | Computational Time (Hour) | | Computational Memory (GB) | |
|---|---|---|---|---|---|
| | | JointPRS | PRS-CSx | JointPRS | PRS-CSx |
| 5 | HDL | 8 | 10 | 2 | 2 |
| | LDL | 7 | 7 | 2 | 2 |
| | TC | 7 | 6 | 2 | 2 |
| | logTG | 8 | 7 | 2 | 2 |
| 3 | Height | 2 | 2 | 1 | 1 |
| | BMI | 2 | 2 | 1 | 1 |
| | SBP | 2 | 2 | 1 | 1 |
| | DBP | 2 | 2 | 1 | 1 |
| | PLT | 2 | 2 | 1 | 1 |
| 2 | WBC | 1 | 1 | 1 | 1 |
| | NEU | 1 | 1 | 1 | 1 |
| | LYM | 1 | 1 | 1 | 1 |
| | MON | 1 | 1 | 1 | 1 |
| | EOS | 1 | 1 | 1 | 1 |
| | RBC | 1 | 1 | 1 | 1 |
| | HCT | 1 | 1 | 1 | 1 |
| | MCH | 1 | 1 | 1 | 1 |
| | MCV | 1 | 1 | 1 | 1 |
| | HB | 1 | 1 | 1 | 1 |
| | ALT | 1 | 1 | 1 | 1 |
| | ALP | 1 | 1 | 1 | 1 |
| | GGT | 1 | 1 | 1 | 1 |

### References

1. Ruan, Y. *et al.* Improving polygenic prediction in ancestrally diverse populations. *Nature Genetics* **54,** 573–580 (2022).

2. Sudlow, C. *et al.* UK biobank: an open access resource for identifying the causes of a wide range of complex diseases of middle and old age. *PLoS medicine* **12,** e1001779 (2015).

3. Zhou, G., Chen, T. & Zhao, H. SDPRX: A statistical method for cross-population prediction of complex traits. *The American Journal of Human Genetics* **110,** 13–22 (2023).

4. Evangelou, E. *et al.* Genetic analysis of over 1 million people identifies 535 new loci associated with blood pressure traits. *Nature Genetics* **50,** 1412–1425 (2018).

5. Graham, S. E. *et al.* The power of genetic diversity in genome-wide association studies of lipids. *Nature* **600,** 675–679 (2021).

6. Of Us Research Program Investigators, A. The "All of Us" research program. *New England Journal of Medicine* **381,** 668–676 (2019).

7. Venner, E. *et al.* The frequency of pathogenic variation in the All of Us cohort reveals ancestry-driven disparities. *Communications Biology* **7,** 174 (2024).
